# Genome evolution of a non-parasitic secondary heterotroph, the diatom *Nitzschia putrida*

**DOI:** 10.1101/2021.01.24.427197

**Authors:** Ryoma Kamikawa, Takako Mochizuki, Mika Sakamoto, Yasuhiro Tanizawa, Takuro Nakayama, Ryo Onuma, Ugo Cenci, Daniel Moog, Samuel Speak, Krisztina Sarkozi, Andrew Toseland, Cock van Oosterhout, Kaori Oyama, Misako Kato, Keitaro Kume, Motoki Kayama, Tomonori Azuma, Ken-ichiro Ishii, Hideaki Miyashita, Bernard Henrissat, Vincent Lombard, Joe Win, Sophien Kamoun, Yuichiro Kashiyama, Shigeki Mayama, Shin-ya Miyagishima, Goro Tanifuji, Thomas Mock, Yasukazu Nakamura

**Author notes:** Corresponding author (RK): Graduate School of Agriculture, Kyoto University, Kitashirakawa oiwake cho, Sakyo ku, Kyoto, Kyoto 606-8502, Japan. Max Planck Institute for Terrestrial Microbiology, Karl-von-Frisch-Str. 10, 35043, Marburg, Germany (Daniel Moog).

## Abstract

Secondary loss of photosynthesis is observed across almost all plastid-bearing branches of the eukaryotic tree of life. However, genome-based insights into the transition from a phototroph into a secondary heterotroph have so far only been revealed for parasitic species. Free-living organisms can yield unique insights into the evolutionary consequence of the loss of photosynthesis, as the parasitic lifestyle requires specific adaptations to host environments. Here we report on the diploid genome of the free-living diatom *Nitzschia putrida* (35 Mbp), a non-photosynthetic osmotroph whose photosynthetic relatives contribute ca. 40% of net oceanic primary production. Comparative analyses with photosynthetic diatoms revealed that a combination of genes loss, the horizontal acquisition of genes involved in organic carbon degradation, a unique secretome and the rapid divergence of conserved gene families involved in cell wall and extracellular metabolism appear to have facilitated the lifestyle of a non-parasitic, free-living secondary heterotroph.

---

The loss of photosynthesis in photoautotrophs seems to be accomplished if such loss is compensated by a competitive advantage arising from the availability of an extracellular energy source. Some secondary heterotrophs have evolved as parasites (Freese and Lane 2017; Hadariová et al. 2018; Janouškovec et al. 2019), relying on sufficient resources provided by their hosts. Well studied examples are the Apicomplexa (e.g., Kissinger et al. 2002), which have lost photosynthesis secondarily. However, examples of loss of photosynthesis found in free-living secondary heterotrophs are as common as those of parasites (Kamikawa et al. 2015; Hadariová et al. 2018; Dorrell et al. 2019; Kayama et al. 2020a; Kayama et al. 2020b), and thereby such examples provide novel insights into evolutionary processes required to thrive without photosynthesis and independently of a resource-providing host. Given that a parasitic lifestyle accelerates the rate of evolution (*cf*. Red Queens hypothesis, Van Valen 1974) and of loss of conserved orthologous genes (e.g., Sun et al. 2018), the genome analysis of a non-parasitic secondary heterotroph can provide insights uncompromised by parasite-specific adaptations. The diatom *Nitzschia putrida* distributing in the mangrove estuaries is the ideal model to test these hypotheses because it is a rare example of a free-living secondary heterotroph (Li and Volcani 1987; Kamikawa et al. 2015a) within the diverse group of largely photoautotrophic diatoms (Field et al. 1998; Mann 1999). As several genomes of the latter have recently become available including close phylogenetic relatives (Armbrust et al. 2004; Bowler et al. 2008; Mock et al. 2012), a genome-based comparative metabolic reconstruction of *N. putrida* promises to reveal novel insights into what is required to thrive as a free-living secondary heterotroph. Here, we have analysed the draft genome sequence of a non-photosynthetic, obligately heterotrophic diatom species (Bacillariophyceae). We provide insight into evolutionary processes underpinning lifestyle shifts from photoautotrophy to free-living heterotrophy in the context of the surface ocean ecosystem.

## Results

### Genome assembly

K-mer-based GenomeScope analysis (Vurture et al. 2017) with 150 bp-long Illumina short reads suggested the genome of *Nitzschia putrida* (Fig. 1A) to be diploid (Supplementary Fig. 1A). To provide a high-quality genome with long-range contiguity, PacBio sequencing (RSII platform) was performed resulting in ≥ 40-fold coverage. Due to the confirmed diploid nature of the *N. putrida* genome, we have applied the Falcon assembler and Falcon_unzip ver. 0.5 (Chin et al. 2016) to provide a first draft genome of this species. Based on this assembly, we estimated a genome size of 35 Mbps, including 87 scaffolds with an N50 of 860.9 kb. The longest scaffold was 3.8 Mbps. The heterozygous regions of the genome (alternate contigs) estimated by the Falcon assembler resulted in 12 Mbps, with an N50 of 121 kbps (Table S1). The Falcon assembly was error corrected and polished by approximately 150-fold coverage of Illumina short reads, which were subsequently used for generating the final assembly with Pilon 1.2.2 (Walker et al. 2016) including manual curation.

**Fig. 1.**
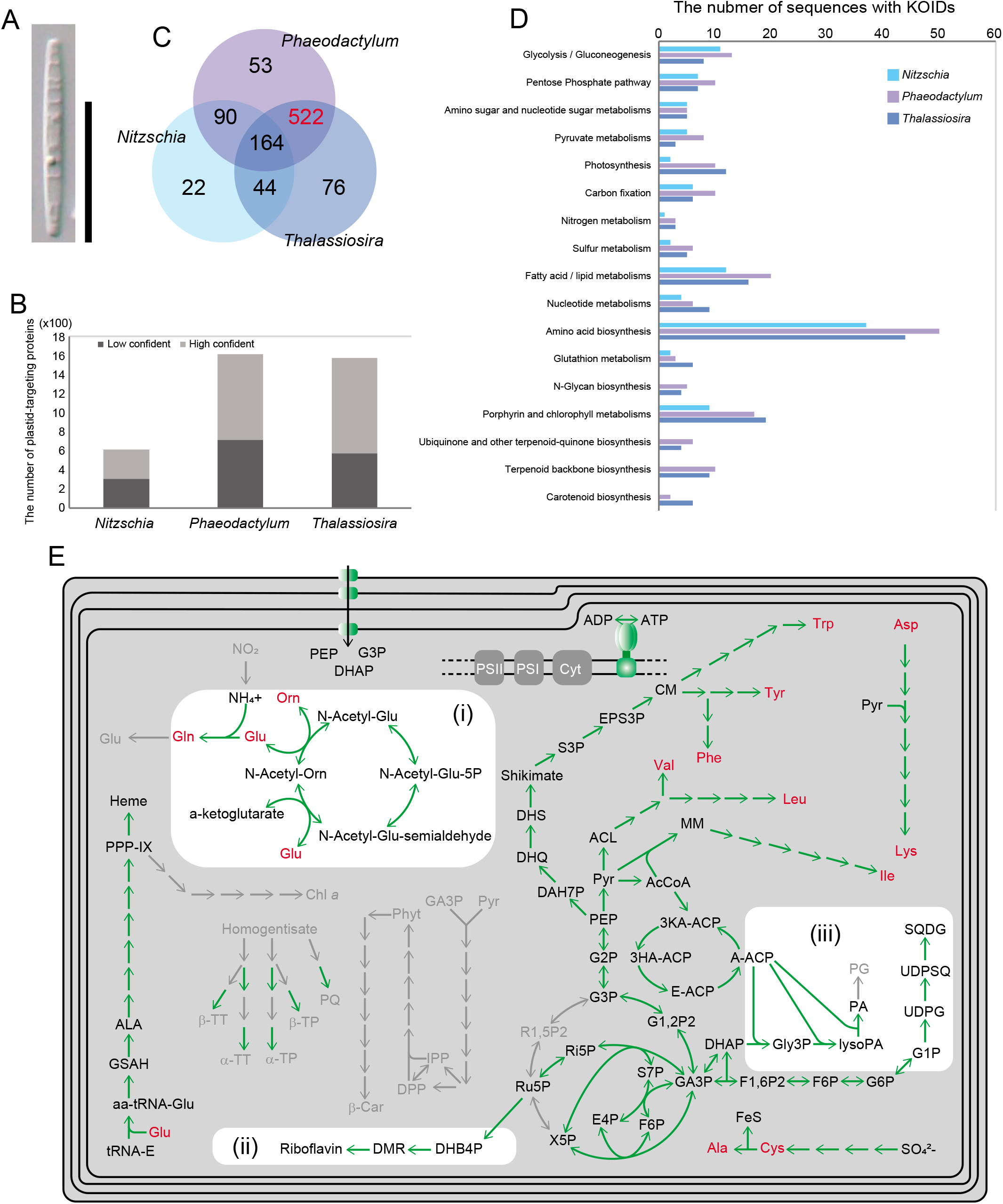
The heterotrophic diatom *Nitzschia putrida* and its plastid proteome. **A.** The frustule view of *N. putrida*. Bar = 20 μm. **B.** Estimated plastid proteome size in three diatoms. Light and dark grey bars show low and high confident plastid-targeted proteins identified by ASAFind, respectively. Data of two photosynthetic diatoms *Phaeodactylum tricornutum* and *Thalassisira pseudonana* are derived from Gruber et al. (2007). **C.** Unique and shared plastid-targeted orthogroups. High lighted in red is the orthogroup exclusively shared by the two photosynthetic diatoms. **D.** Comparison of KO ID numbers among plastids of the three diatoms. Each bar indicates numbers of unique KO IDs in each functional category. **E.** Predicted metabolic map of the non-photosynthetic plastid. Representative pathways found in photosynthetic diatom species are shown. Green and light grey arrows show presence and absence of the responsible protein sequences for the reactions in the genome, respectively. Amino acids are highlighted in red. Glu: Glutamate, Gln: Glutamine, Orn: Ornithine, NH^4+^: ammonium, NO2: nitrite, N-Acetyl-Glu: N-Acetylglutamate, N-Acetyl-Glu-5P: N-Acetylglutamyl-5phosphate, N-Acetyl-Glu-semialdehyde: N-Acetylglutamyl-semialdehyde, a-ketoglutarate: alpha-ketoglutarate, PPP-IX: protoporphyrin-IX, ALA: 5-aminolevulinic acid, GSAH: Glutamate-1-semialdehyde, aa-tRNA-Glu: amino acyl-tRNA-glutamate, tRNA-E: glutamyl tRNA, Chl *a*: chlorophyll *a*, α-TT: alpha-Tocotrierol, β-TT: beta-tocotrierol, α-TP: alpha-tocopherol, β-TP: beta-tocopherol, PQ: plastoquinone/plastoquinol, GA3P: glyceroaldehyde-3phosphate, Pyr: Pyruvate, IPP: Isopentenylpyrophosphate, DPP: Dimethylallylpyrophosphate, Phy: Phytoene, β-Car: beta-carotene, DHB4P: 3,4-Dihydroxy 2 butanone 4phosphate, DMR: 6,7-Dimethyl-8-ribityl umazine, Asp: Aspartate, Lys: Lysine, Ile: Isoleucine, Leu: Leucine, Val: Valine, MM: Methylmalate, ACL: Acetolactate, AcCoA: Acetyl-CoA, ACP: acyl-carrier protein, 3KA-ACP: 3ketoacyl-ACP, A-ACP: acyl-ACP, E-ACP: enoyl-ACP, 3HA-ACP: 3hydroxyacyl-ACP, PEP: phosphoenolpyruvate, G2P: 2 phosphoglycerate, G3P: 3 phosphoglycerate, DAH7P: 3-deoxy-7-phosphoheptulonate, DHQ: 3-dehydroquinic acid, DHS: 3-dehydroshikimate, S3P: shikimate 3phosphate, EPS3P: 5-enolpyruvylshikimate −3-phosphate, CM: Chorismate, Trp: Tryptophan, Tyr: Tyrosine, Phe: Phenylalanine, G1,2P2: Glycerol 1,2 bisphosphate, R1,5P2: Ribulose 1,5 bisphosphate, Ru5P: Ribulose 5phosphate, Ri5P: Ribose 5phosphate, S7P: sedoheptulose 7-phosphate, E4P: Erythrose 4phosphate, F6P: Fructose 6phosphate, X5P: Xylulose 5phosphate, F1,6P2: Fructose 1,6 bisphosphate, Gly3P: Glycerate 3phosphate, LysoPA: Lysophosphatidic acid, PA: Phosphatidic acid, PG: Phosphatidyl glycerol, G6P: Glucose 6phosphate, G1P: Glucose 1phosphate, UDPG: UDP-glucose, UDPSQ: UDP-sulfoquinovose, SQDG: Sulfoquinovosyl diacylglycerol, SO_4_^2-^: sulfate, Cys: Cysteine, FeS: Iron-sulfur cluster, Ala: Alanine, Cyt: cytochrome *b6/f* complex, PSI: photosystem I, PSII: Photosystem II.

According to the k-mer assessed diploid nature of the *N. putrida* genome, the read coverage of the homozygous regions is approximately two-fold higher than the read coverage for the heterozygous regions, suggesting the presence of diverged alleles as previously identified in the genome of the photoautotroph diatom *Fragilariopsis cylindrus* (Supplementary Figs. 1A & 1B). Thus, most of the diverged allelic variants can be found in the heterozygous regions characterized by the presence of alternate contigs (Supplementary Fig. 1B). On the basis of the analysis with Braker2 v.2.0.3 (Hoff et al. 2016), the *Nitzschia* genome comprises 15,003 and 5,767 inferred protein-coding loci on the primary and alternate contigs, respectively (Table S1). Almost 40% of loci in the genome of *N. putrida* appear to be characterised by diverged alleles. A BUSCOv3 analysis (Waterhouse et al. 2018) revealed the genome to be complete at a level of 90.1% based on the haploid set of genes.

### The loss of photosynthesis

The haploid set of genes was used to reconstruct the nuclear-encoded plastid proteome of *N. putrida* and therefore to reveal the extent of gene loss including key genes of photosynthesis. A comparative analysis of the *N. putrida* plastome (Gruber et al. 2015) with its photosynthetic counterparts revealed that more than 50% of nuclear encoded plastid proteins have been lost (Fig. 1B). More than 500 orthogroups (Orthofinder, Emms and Kelly 2015) of nuclear-encoded plastid proteins which are usually shared between photosynthetic diatoms (Gruber et al. 2015) are missing in the predicted plastid proteome of *N. putrida* (Fig. 1C). In the missing part of the plastid proteome were genes encoding for proteins of light-harvesting antenna including fucoxanthin-chlorophyll *a/c* protein *fcp*), photosystem II and I (e.g. *psbA, psbC, psbO, psaA, psaB*, and *psaD*), the cytochrome *b6/f* complex (e.g., *petA*) and carbon fixation (e.g. *rbcS, rbcL*) in addition to genes of the Calvin cycle (e.g. phosphoribulokinase (*prk*)). Furthermore, a significant number of key genes were missing for the biosynthesis of chlorophyll, carotenoids, and plastoquinones (Fig. 1D).

Despite the loss of some of these key photosynthesis genes, unexpectedly, there is still a significant number of genes left encoding common plastid metabolic pathways as known from photosynthetic diatoms, including the generation of ATP by a nuclear-encoded ATPase subunit (Kamikawa et al. 2015b). Almost all genes encoding for plastid enzymes to synthesize essential amino acids are still encoded in the nuclear genome of *N. putrida*. Furthermore, all genes of the heme pathway have been found, and *N. putrida* appears to be able to synthesise riboflavin. The presence of plastid-targeted transporters (Moog et al. 2020) enable the transport of phosphoenolpyruvate (PEP), 3-phosphoglycerate (G3P), and dihydroxyacetone-phosphate (DHAP) across the plastid membranes. Additionally, our genome-based reconstruction of plastid metabolism detects the biosynthesis pathway for lipids and the ornithine cycle in *N. putrida* (Fig. 1E; Supplementary Fig. 2). The latter has neither been reported in previous transcriptome-based studies with this species (Kamikawa et al. 2017) nor in any other secondary heterotrophs (e.g., Dorrell et al. 2019; Kayama et al. 2020).

### Communication between organelles and light-dependent gene expression

The lack of CO_2_ fixation in plastids of *N. putrida*, which reduces the amount of amino acids, lipids and other metabolites to be synthesised, appears to be partially compensated by the remodelling of metabolic interactions with mitochondria and peroxisomes as well as by the keeping nitrogen recycling (Fig. 1; Fig. 2A; Supplementary Fig. S3&S4). It appears that the non-photosynthetic plastid of *N. putrida* still exchanges glutamine and ornithine, both of which are important intermediates of the ornithin cycle. Indeed, all genes for the ornithine-urea cycle have been retained in the *N. putrida* genome. The ornithine-urea cycle is indispensable for nitrogen recycling in photosynthetic diatoms (Allen et al. 2011; Smith et al. 2019), and even after the loss of photosynthesis, nitrogen recycling appears to be essential in *N. putrida* (Fig. 2A) due to its osmotrophic lifestyle. Usually, the ornithine-urea cycle is tightly linked with TCA cycle and/or photorespiration in photosynthetic diatoms (Allen et al. 2011; Smith et al. 2019). However, *N. putrida* is not likely to perform photorespiration (Fig. 2A). The metabolic exchange with the peroxisome through glycolate likely has ceased as phosphoglycolate phosphatase and peroxisomal glycolate oxidase are missing. Thus, photorespiration is unlikely to take place in non-photosynthetic plastids of *N. putrid* due to the lack of Ribulose 1, 5-bisphosphate carboxylase/oxygenase (RuBisCO) and other key enzymes of the Calvin cycle (Fig. 1). Nevertheless, peroxisomes still appear to play a role in *N. putrida* for the production of malate or glyoxylate, which feed into respiratory pathways of the mitochondria to support ATP and NADPH production (Fig. 2A).

**Fig. 2.**
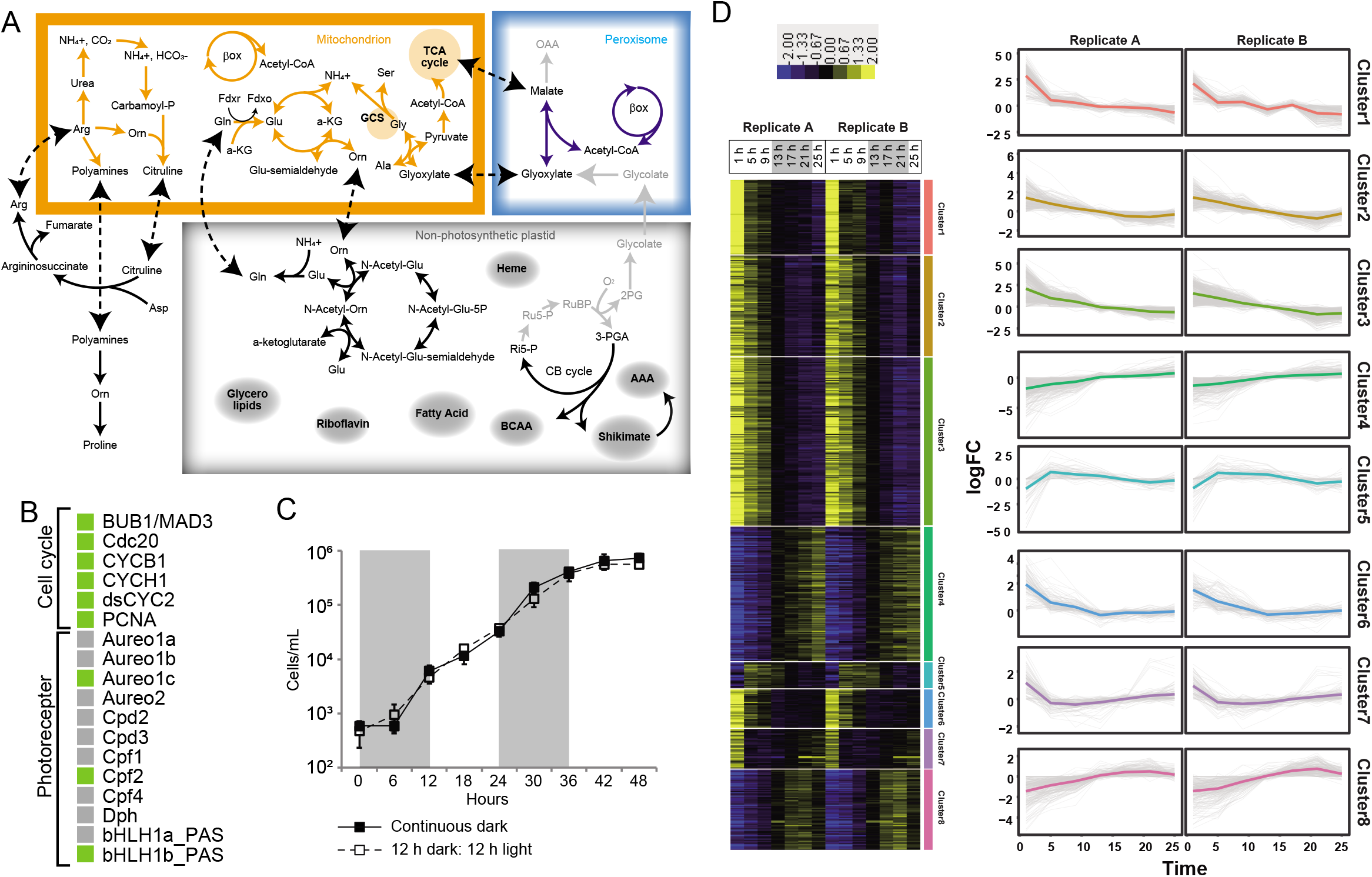
Loss of genes for the plastid-persoxisome metabolic flow and photoreceptors. **A.** Metabolic interactions between a mitochondrion and a non-photosynthetic plastid and between a mitochondrion and a peroxisome. Black, orange, and blue arrows show presence of responsible protein sequences for the reactions in a plastid, a mitochondrion, and a peroxisome, respectively, while light grey arrows show absence of responsible protein sequences. Dashed arrows show possible inter-organellar metabolic flows. NH4+: ammonium, CO2: carbon dioxide, HCO3-: bicarbonate, Arg: Arginine, Orn: Ornithine, Asp: Aspartate, βox: fatty acid β oxidation, Gln: Glutamine, Glu: Glutamate, a-KG: alpha-ketoglutarate, Ser: Serine, Gly: Glycine, GCS: Glycine cleavage system, Ala: Alanine, TCA: Tricarboxylic acid cycle, OAA: oxaloacetate, BCAA: Branched chain amino acid synthesis, AAA: aromatic amino acid synthesis. Other abbreviations are described in Fig. 1. B. Photoreceptor and cell cycle genes in the *N. putrida* genome. The other genes are shown in Supplementary Fig. S5. Light green and light grey boxes show presence and absence of corresponding genes, respectively. **C.** Growth of the heterotrophic diatom under the different light conditions. Closed boxes show growth in the continuous dark condition, while open boxes show growth in the light-dark condition. Shaded in grey are the dark periods in the light-dark cultivation conditions. **D.** (Left) Heatmap showing the reproducible expression patterns of genes (Pearson’s correlation coefficient < 0.9). *k*-means clustering was calculated for each gene based on RPKM +1 values, which were transformed to log2 and centred by median values. Yellow and blue indicate upregulation and downregulation of the gene, respectively. (Right) The line graphs showing expression pattern of genes in each cluster. The coloured line indicates the average value of the expression patterns.

Light in photosynthetic organisms does not only play a significant role for photosynthesis generating ATP and NADPH, it also regulates cell division, diel cycles and different signalling processes unlike in many heterotrophic organisms (Ashworth et al. 2013; Chauton et al. 2013; Smith et al. 2017). Thus, we identified remaining photoreceptors and cell-cycle regulators and their effect on regulating diel cycles and genome-wide light-dependent gene expression. Although we found that all the cyclin and cyclin-dependent kinases (Huysman et al. 2010) were still encoded and expressed in the genome of *N. putrida* (Fig. 2B; Supplementary Fig. S5A&S5B), we were unable to identify a diel cycle in cell division over a growth period between 12 hours light and 12 hours dark condition and 48 hours darkness with cells previously acclimatised to a periodic change of light and dark for 12 hours each (Fig. 2C). This suggests these cell-cycle regulators potentially have neo/subfunctionalized and therefore have a different redulatory role in *N. putrida* unrelated to the diel cycle. The loss of the transcription factor *bHLH-1a* (RITMO1), which has been identified as a master regulator of diel periodicity (Annunziata et al. 2019), corroborates our finding that *N. putrida* has lost the ability to perform diel cycles. In addition, most of the other photoreceptors known from photosynthetic diatoms have also been lost (Fig. 2B) such as the blue-light sensing Aureochromes 1a/b, both of which are transcription factors responsible for photoacclimation (Kroth et al. 2017). Despite the lack of light-dependent cell-cycle regulation, a few remaining photoreceptors were identified including bHLH1b_PAS, Aureochrome 1c, and Cryptochrome-DASH/CPF2 (Fig. 2B) (Coesel et al. 2009; Ashworth et al. 2013). Basic ZIP transcription factors possessing potentially light-sensitive PAS domains (bZIP-PAS) (Fortunato et al. 2016), were also identified in the *N. putrida* genome such as homologues to bZIP6 and bZIP7 of *Phaeodactylum tricornutum* (Rayko et al. 2010). The latter has been duplicated and diversified in *N. putrida* (Supplementary Fig. S5C). The presence of bZIP-PAS protein in a heterotrophic eukaryote is not unprecedented as some oomycetes, non-photosynthetic parasites, have been reported to encode them in their genomes (e.g., Kong et al. 2020). Although their role in regulating gene expression remains to be investigated in *N. putrida*, light still appears to influences the expression of some genes in this heterotrophic species. Comparative transcriptome analyses every four hours during a shift from a light phase to darkness (Fig. 2D) revealed eight clusters characterised by different expression patterns, and there is no cluster explicitly representing the light-dependent gene expression patterns as seen in photosynthetic algae (e.g., Ashworth et al. 2013; Fujiwara et al. 2020). However, one of the clusters suggested some genes were expressed only in the mid light phase: cluster 7 containing 90 genes (0.6% total). EuKaryotic Orthologous Groups (KOGs) analysis with these genes detected 44 genes with known functional domains, and of them, 21 are responsible for substrate import and carbon metabolisms (Supplementary Fig. S5D). However, the photoreceptor homologues above, bHLH1b_PAS, Aureochrome 1c and Cryptochrome-DASH/CPF2, were not part of this cluster, and there was no explicit trend in their gene expression patterns with respect to changes between light and dark conditions.

### The acquisition of genes through horizontal gene transfer

Recent studies estimated that 3-5% of genes in diatom genomes were acquired through species-specific horizontal gene transfer (HGT) (Vancaester et al. 2020). For *N. putrida*, we identified 73 genes potentially acquired via HGT based on phylogenetic tree reconstruction. Three of these genes were only shared with photosynthetic and non-photosynthetic species of the genus *Nitzschia*. Hence, they were likely acquired by the last common ancestor of the *Nitzschia* genus (Supplementary Fig. S6&S7). These three genes are one sulfate transporter and two serine hydrolases, which contribute to the sulfate assimilation and degradation of various compounds, respectively. Of the remaining 70 HGT genes, 25 were of bacterial origin and 34 appear to have been originated in other microbial eukaryotes such as fungi. The remaining 11 genes were either of viral or of ambiguous origin (Fig. 3A). As the majority of the HGT genes in *N. putrida* were not identified in any other diatom species, it suggests that they play a role in the heterotrophic lifestyle and therefore might have been essential for the transition from photoautotrophy to secondary heterotrophy. This hypothesis is corroborated by the fact that many of these genes (ca. 50%) are involved in catabolic processes such as carbohydrate metabolism (such as glucose 1P dehydrogenase, GH27 α-galactosidase) and degradation of amino acids (such as arginase). One HGT gene (trichothecene 3-O-acetyltransferase) even appears to convey tolerance against mycotoxin (Khatibi et al. 2011) (Figs. 3B–3E). Thus, the acquisition of these genes may have provided the ancestor of *N. putrida* with the potential to utilize new substrates and to compete fungi, both of which can be considered functional traits essential for a successful transition to becoming a secondary heterotroph. Only five HGT genes encode secreted proteins according to our *in-silico* analysis (see below), which suggests that most of the others contribute to intracellular metabolic processes.

**Fig. 3.**
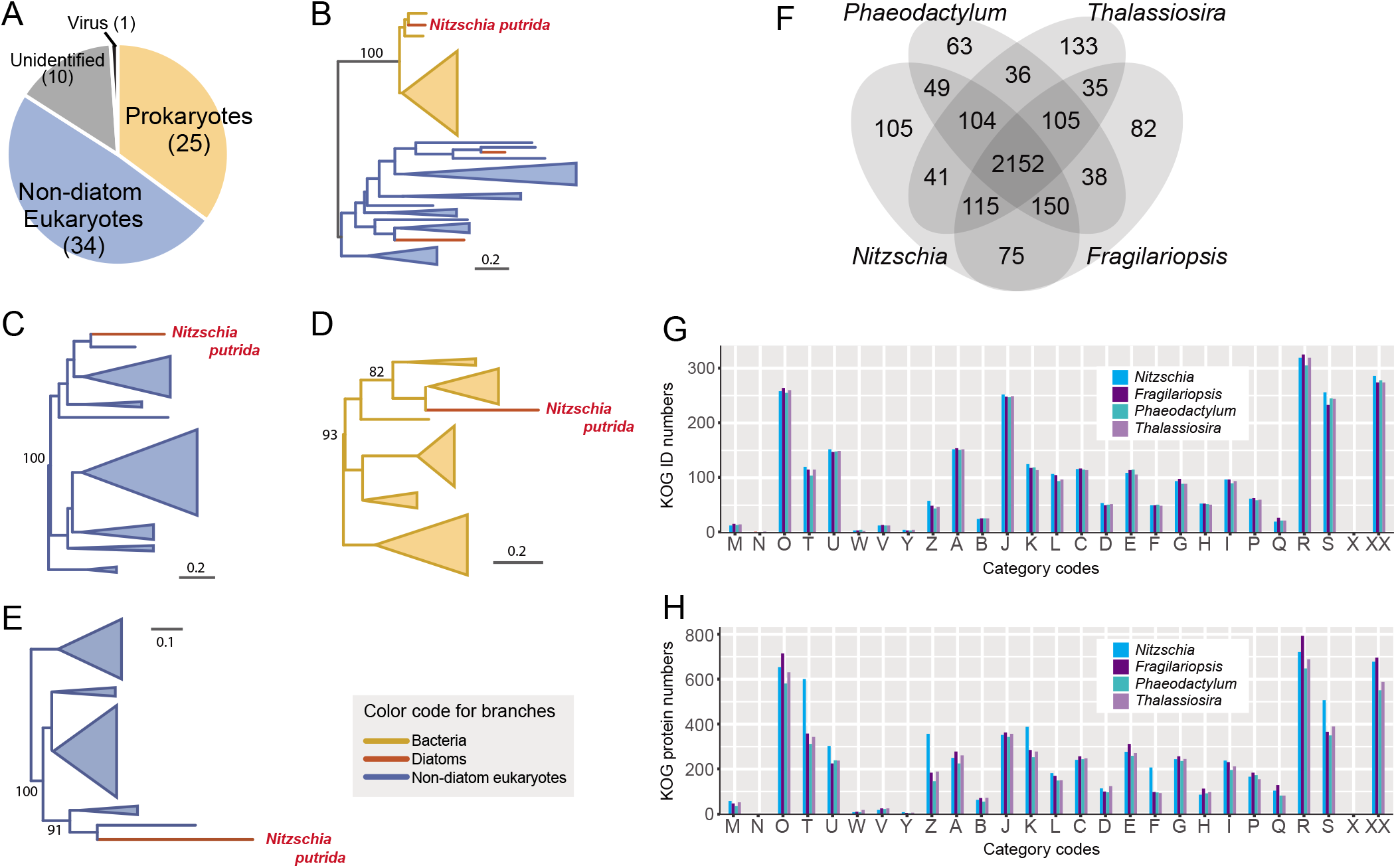
Horizontal gene transfers and gene family diversification in the heterotrophic diatom *Nitzschia putrida*. **A.** Proportion of origins of laterally transferred genes identified in the *N. putrida* genome. **B-E.** Phylogenetic trees of glucose 1P-dehydrogenase (NAA00P12280; B), arginase family protein (NAA12P00140;C), alpha-N-acetylgalactosidase (NAA03P01590; D), and trichothecene 3-O-acetyltransferase, self-protection against trichothecenes (NAA53P00570; E). The numbers on branches are maximum likelihood bootstrap values higher than 80. All the trees showing possible *N. putrida*-specific LGTs are provided in Supplementary Fig. 7. **F.** Venn diagram of KOG IDs shared by four diatoms. *Nitzschia: Nitzschia putrida, Phaeodactylum: Phaeodactylum tricornutum, Thalassiosira*: *Thalassiosira pseudonana*, *Fragilariopsis*: *Fragilariopsis cylindrus*. **G.** Comparison of the number of KOG ID among diatoms. KOG categories are as follows: A, RNA processing and modification; B, chromatin structure and dynamics; C, energy production and conversion; D, cell cycle control, cell division and chromosome partitioning; E, amino acid transport and metabolism; F, nucleotide transport and metabolism; G, carbohydrate transport and metabolism; H, coenzyme transport and metabolism; I, lipid transport and metabolism; J, translation, ribosomal structure and biogenesis; K, transcription; L, replication, recombination and repair; M, cell wall, membrane or envelope biogenesis; N, cell motility; O, post-translational modification, protein turnover, chaperones; P, inorganic ion transport and metabolism; Q, secondary metabolites biosynthesis, transport and catabolism; R, general function prediction only; S, function unknown; T, signal transduction; U, intracellular trafficking, secretion and vesicular transport; V, defence mechanisms; W, extracellular structures; Y, nuclear structure; Z, cytoskeleton. **H.** Comparison of the number of genes assigned to each KOG category. Other details are described above.

### The genetic toolkit for the evolution of non-parasitic secondary heterotrophy

Despite the loss of many nuclear genes and their families, the genome size of *N. putrida* is not significantly different to photosynthetic relatives such as *Fragilariopsis cylindrus*, *Phaedoctylum tricornutum* and the more distantly related diatom *Thalassiosira pseudonana* (Table S1). By comparing KOGs of paralog proteins, there was no significant difference in the number of unique KOG IDs between these four diatom species (Fig. 3F&3G). However, when we compared the number of paralog proteins assigned to each KOG ID, there were several KOG categories for which *N. putrida* had a higher number of paralogous proteins compared to the other diatom species: nucleotide transport (F), transcription (K), signal transduction (T), intracellular trafficking, secretion, vesicular transport (U), and cytoskeleton (Z) (Fig. 3H). Even after normalization by total gene numbers, nucleotide transport (F), signal transduction (T), and cytoskeleton (Z) genes were more abundant in the *N. putrida* genome (Supplementary Fig. S8). This observation was corroborated by *N. putrida-specific* enrichment of important Pfam domains in these functions such as Adenylate/Guanylate cyclase and Cyclic nucleotide esterase, Leucine rich repeat (LRR), and glycosyl/galactosyl transferase domains (Supplementary Fig. S8).

A microbial heterotroph either acquires nutrients by phagotrophy, the preferred nutrition of many parasites, or by osmotrophy, which is the uptake of dissolved organic compounds by osmosis as realised by bacteria and fungi, for instance (Richards and Talbot 2013; Richards and Talbot 2018). As *N. putrida* grows well under axenic conditions (Kamikawa et al. 2015; Ishii & Kamikawa 2017), it is likely an osmotroph, dependent on the uptake of dissolved organic compounds across the silicified cell wall and the plasma membrane. As realised by osmotrophic fungi, *N. putrida* may even be able to degrade higher molecular weight compounds extracellularly to be subsequently taken up as individual molecules by specific transporters or even osmosis (Richards and Talbot 2013; Richards and Talbot 2018). Thus, it is likely that cell wall, membrane, and secreted proteins diversified in *N. putirda* compared to photosynthetic diatoms to facilitate osmotrophy. We analysed the enrichment of paralog proteins and differences in nutrient transporters involved in the uptake of dissolved organic compounds such as solute carriers.

Although the *N. putrida* genome does not differ in the abundance of nutrient transporters relative to photosynthetic diatoms (Fig. 4A), we did find a significant difference in the composition of these genes. For instance, the number of genes encoding silicon transporters, solute symporters, and the resistance-nodulation-cell division superfamily were more than twice as large in *N. putrida* compared to photosynthetic diatom species (Fig. 4B; Supplementary Fig. S9).

**Fig. 4.**
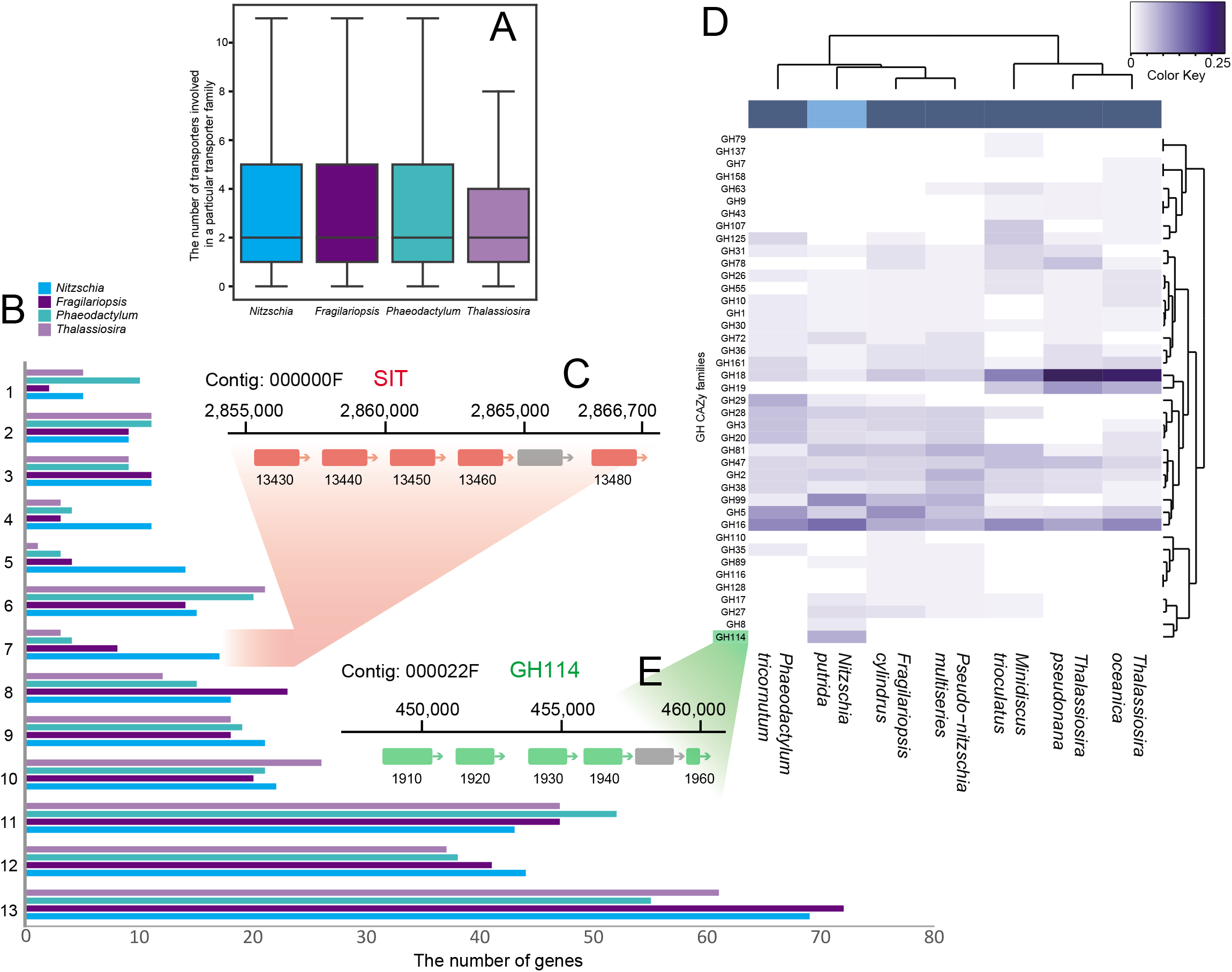
Diversity of transporters and carbohydrate active enzymes in *N. putrida*. **A.** Distribution of the number of transporters in each transporter family. There is no significant difference detectable by Wilcoxon signed rank test. **B.** The gene number of transporters in the 10 most abundant transporter families of *N. putrida*. 1. Resistance-Nodulation-Cell Division (RND) Superfamily, 2. Solute:Sodium Symporter (SSS) Family, 3. Voltage-gated Ion Channel (VIC) Superfamily, 4. Silicon Transporter (SIT) Family, 5. Amino Acid/Auxin Permease (AAAP) Family, 6. P-type ATPase (P-ATPase) Superfamily, 7. Drug/Metabolite Transporter (DMT) Superfamily, 8. Mitochondrial Carrier (MC) Family, 9. Major Facilitator Superfamily (MFS), 10. ATP-binding Cassette (ABC) Superfamily. **C.** Silicon transporter (SIT) genes tandemly located in the contig 000000F. SIT genes are highlighted in light red with the gene IDs. **D.** Glycoside Hydrolases (GH) families from the Carbohydrate Active enZyme (CAZy) database focused on Bacillariophyta. The diagram shows a heatmap of CAZyme prevalence in each taxon (number of a particular CAZyme family divided by the total number of CAZyme genes in the organism); the white to blue color scheme indicates low to high prevalence, respectively. Dendrograms (left and top) show respectively the relative taxa proximity with respect co-occurrence of CAZyme families and the co-occurrence of CAZyme families with one another within genomes (according to the method described in Cenci et al. 2018). **E.** GH114 genes tandemly located in the contig 000022F. GH114 genes are highlighted in light green with the gene IDs.

Expansion of those gene families may, at least partly, have been achieved by recent tandem duplications (Fig. 4C). To gain insight into when the expansion had occurred, we performed a coalescence analysis, which revealed that silicon transporters (SITs) in *N. putrida* began to expand around 3.3 Mya, while divergence from another non-photosynthetic diatom *N. alba* is estimated to have occurred around 6.67 Mya (Supplementary Fig. S10). Thus, their recent expansion suggests neo/subfunctionalisation of the gene family in response to the change in lifestyle. The divergence rate of SIT genes was much larger than that of control genes (e.g., myosin), indicating that SIT diversification might have contributed to the adaptation of the heterotrophic lifestyle. In support of this hypothesis, we detected several sites under positive selection in different members of the SIT family (Table S2), which implies that the evolution of those genes may have been driven by diversifying selection.

The solute sodium symporters are estimated to have diverged around 7.3 Mya, markedly earlier than the SIT gene family. Although the divergence rate is also larger than that of control genes (Supplementary Fig. S10), we did not find evidence of diversifying selection in this gene family. The differences between these two families of transporters suggest that their expansion might have occurred in a stepwise manner and driven by different evolutionary forces.

Furthermore, although the overall carbohydrate-active enzymes (CAZyme) family composition of *Nitzschia* was not remarkably different from that of photosynthetic diatoms (Supplementary Fig. S11), families encoding β-glycoside hydrolase (GH8), laminarinase (GH16_3), pectinase (GH28), β-glucanase (GH72), α-mannan hydrolyzing enzymes (GH99), and β-1,2-glucan hydrolytic enzymes (GH114) were also enriched in *N. putrida* compared to photosynthetic species (Fig. 4D). Expansion of these families might, at least partly, have been achieved by recent tandem duplications (Fig. 4E). Notably, more than one third of proteins assigned to the above six families are predicted to be secreted in *N. putrida*.

### The predicted secretome of the non-parasitic, free-living secondary heterotroph *N. putrida*

Given that the secretome plays an important role for substrate degradation and subsequent uptake of low-molecular weight compounds in osmotrophs (Richards and Talbot 2013), we conducted a comparative analysis to predict secreted proteins of *N. putrida in-silico* by idintifying proteins with N-terminal signal peptides and a lack of transmembrane domains. The resulting proteins were clustered using TribeMCL (Enright et al. 2002), and plastid- and lysosome-localised proteins were subsequently removed using ASAFind according to their characteristic targeting motifs (Gruber et al. 2005) and Pfam domains. The number of putatively secreted proteins is 978, 998, 596, and 718, in *N. putrida, F. cylindrus, P. tricornutum*, and *T. pseudonana*, respectively, which corresponds to between 5 and 7% of the total number of genes in their genomes (Supplementary Fig. S12A). Nevertheless, there were significant differences when we compared the diversity of proteins for each between the four diatom species (Figs. 5A & 5B); *N. putrida* on average had a significantly higher number of proteins per tribe than any of the other diatom species (Two-sided Wilcoxon signed rank test; *p* < 0.01; Fig. 5C). Especially, proteins involved in heterotrophy such as organic matter degradation/modification including CAZymes and peptidases were more abundant in *N. putrida* than in the photosynthetic diatom genomes (188 in *N. putrida*, 142 in *F. cylindrus*, 118 in *P. tricornutum*, and 101 in *T. pseudonana;* Supplementary Fig. S12A).

**Fig. 5.**
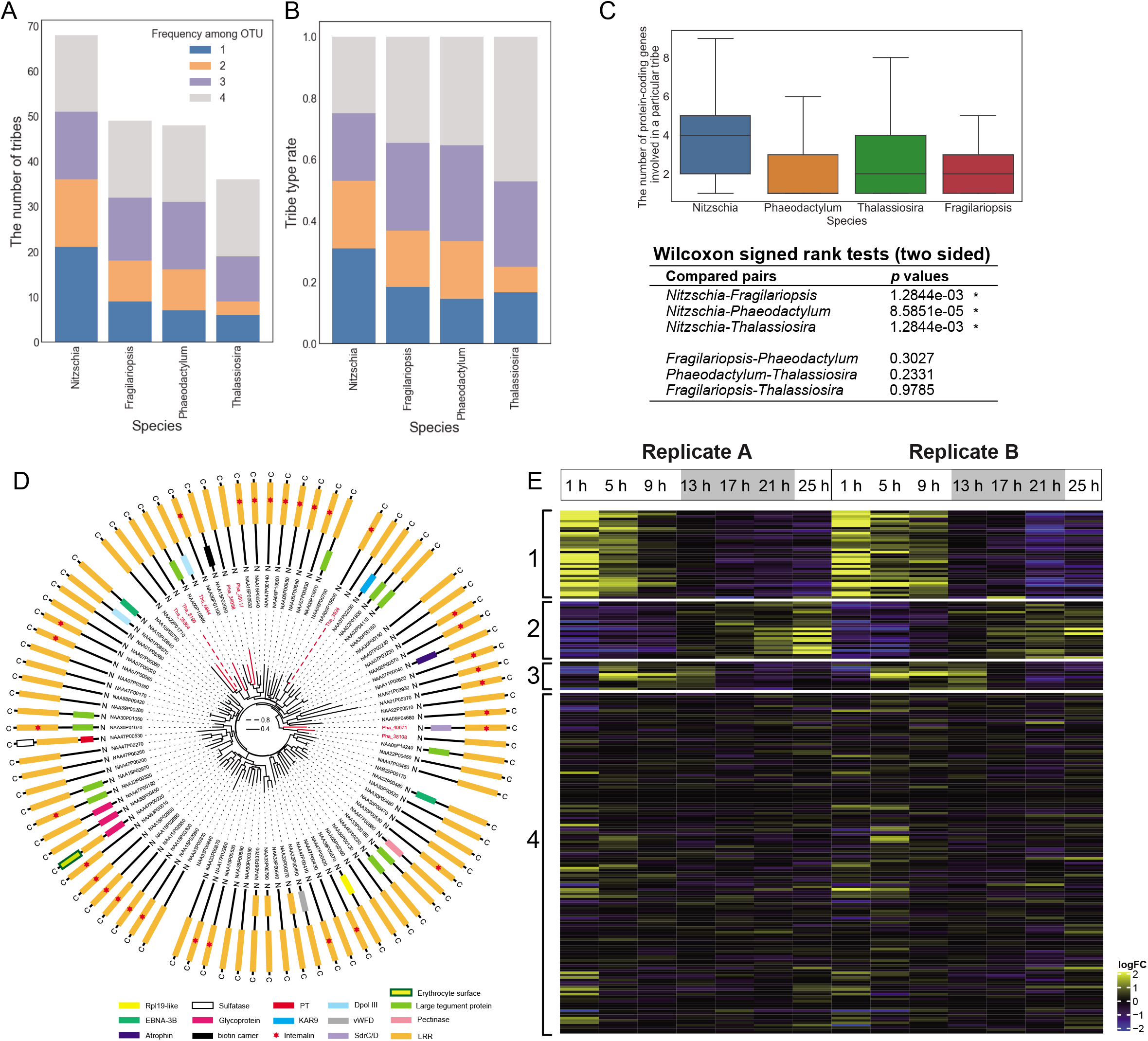
Secretome of *N. putrida*. **A.** The number of secretome tribes, including at least four sequences, clustered by TribeMCL (Enright et al. 2002). Different colours represent tribe categories as follows: 1. Species specific tribes, 2: tribes shared by two species, 3: tribes shared by three species, and 4: tribes shared by all the four diatoms. **B.** Proportion of each tribe category in diatoms. Details are described in **A. C.** Distribution of the number of protein sequences in each secretome tribe. Each box represents the interquartile range between the first and third quartiles (25th and 75th percentiles, respectively), and the median is represented by the vertical line inside the box. The lines protruding from either side of the box are the lowest and highest values within 1.5 times the interquartile range from the first and third quartiles, respectively. Outliers were omitted in the boxplot. The results of the Wilcoxon signed rank test was shown below the box plot. *p*-values for three pairs of comparisons that include *N. putrida* as a counterpart were adjusted by the Benjamini–Hochberg procedure. Asterisks show significantly different pairs under *p* < 0.01. **D.** Sequence diversity of Leucin-rich repeat protein sequences. Phylogenetic tree centred in the figure was reconstructed by IQ-TREE. Domains contained in each sequence were predicted by NCBI Conserved Domain Search and depicted next to the gene IDs; in only several sequences no domain was predicted regardless of apparent homologies (e^-30^ in TribeMCL) to LRR proteins. **E.** Expression of the 10 largest tribes in *N. putrida* during the 25 hours cultivation. Heatmap showing the expression patterns of genes in two independent experiments. *k*-means clustering was calculated for each gene based on RPKM + 1 values, which were transformed to log2 and centred by median values. Yellow and blue indicate upregulation and downregulation of the gene, respectively.

The most common secreted proteins in *N. putrida* are leucine rich repeat-containing (LRR) proteins (Supplementary Fig. 12B), many of which contain additional domains such as tegument and glycoprotein domains, suggesting an increased functional diversity (Fig. 5D). Interestingly, only very few LRR-containing proteins were identified in the predicted secretomes of the photosynthetic diatoms, indicating that signal-peptide-dependent secretion of abundant and diverse LRR-containing proteins maybe an essential requirement in this secondary heterotroph. In addition to LRR-containing proteins, the top ten most enriched proteins in *N. putrida* were Von Willebrand factor type D involved in adhesion or clotting, two types of endopeptidases, trypsin and leishmanolysin (cell surface peptidase of the human parasite *Leishmania*), intradiol ring-cleavage dioxygenase protein for degradation of aromatic compounds, methyltransferase, and four proteins with unknown function (Supplementary Fig. 12B). Transcriptional dynamics of the predicted secretome over a diel cycle (Fig. 2) revealed the presence of four different clusters. Genes in cluser 1 were transcribed at the beginning of the first light phase and genes in cluster 2 at the end of the dark phase and into the second light phase (Fig. 5E). Genes of clusters 3 were most strongly expressed in the middle and end of the first light phase, whereas genes in cluster 4 were relatively weakly expressed throughout day and night. These results suggest that some stimuli including light conditions or nutrient conditions play a role in the regulation of these genes, which might either be a relict from the photosynthetic ancestor or a response to diel cycles of organic substances in the aquatic system occupied by *N. putrida*. As only five of the secreted proteins potentially have been acquired via horizontal gene transfer (Supplementary Fig. 7), the origin of the secretome in *N. putrida* most likely is derived from a photosynthetic ancestor.

## Discussion

*N. putrida* experienced a series of genetic adaptations towards a heterotrophic lifestyle. *N. putrida* took a step backwards in one of the major evolutionary transitions, from photoautotrophs to heterotrophs, potentially relaxing selection on some of the now redundant gene networks and functions. As expected, more than 50% of nuclear encoded plastid proteins have been lost in the *N. putrida* plastid proteome in comparison to its photosynthetic counterparts (Gruber et al. 2015). However, the total number of genes (~15,000) fell within the range of photosynthetic microalgae, and we found no evidence of pseudogene formation, genome streamlining (e.g., Wolf and Koonin 2013), gene family contraction (cf. birth-and-death hypothesis, Nei and Kumar 2000), or reductive genome evolution (Black Queen Hypothesis, Morris et al. 2012). The relatively large genome size is not unexpected given that *N. putrida* is a free-living osmotroph. This free-living lifestyle in a complex and highly variable coastal marine environment likely is the reason why a significant number of genes including some photoreceptors, cell-cycle regulators, and common plastid metabolic pathways present in photosynthetic diatoms have remained. Even though some of the latter genes were still expressed, *N. putrida* appears to lack a diel growth cycle, which suggests that these cell-cycle regulators have neo/subfunctionalized. However, as a cetain number of genes still appears to be regulated by light, osmotrophy potentially benefits from diel fluctuations of resouces such as dissolved organic carbon in aquatic environments (Ottesen et al. 2014; Aylward et al. 2015; Frischkorn et al. 2018). For photoautotrophs, it is important to regulate the cell cycle in accordance with diel cycles for optimising photosynthesis and therefore cell proliferation (Ashworth et al. 2013; Chauton et al. 2013; Ottesen et al. 2013; Smith et al. 2017; Hernández Limón et al. 2020). Without being reliant on light as its primary energy source, the osmotroph *N. putrida* no longer requires coordinating its cell cycle with diel cycles. Thus, after the loss of photosynthesis, the strict light-dependent regulation of gene expression might have become less important and gene expression therefore may have become predominantly regulated by other stimuli. Indeed, many photoreceptors are missing but duplication of genes for bZIP transcription factors with PAS domains and genes for signal transduction and cellular regulatory roles such as adenyl/guanyl cyclase and cyclic nucleotide esterase domains were enriched in the *N. putrida* genome. Furthermore, the peroxisome-plastid interaction is no longer requried after the loss of photosynthesis giving rise to loss of carbon fixation in the context of glycolate recycling. In contrast, the ornithin-urea cycle likely remains to be functional to faciliate nitrogen recycling.

Gene family expansions and neo/subfunctionalizations appear to have played a prominent role in the adaptation to its new lifestyle given that many proteins predicted to be secreted have diversified in *N. putirda*, possibly to facilitate osmotrophy. Altogether, the drastic change of lifestyle associated with the “devolution” in light of autotrophic lifestyles did not result in the reductive genome evolution such as loss in gene number, predicted from non-photosynthetic plastid-bearing parasites.

## Methods

### Cultivation, DNA and RNA extraction, and sequencing

*Nitzschia putrida* NIES-4239 was cultivated in the daigo’ IMK medium (Wako) including 1% Luria–Bertani medium based on the artificial seawater made with MARINE ART SF-1 (Osaka Yakken Co.) at 20°C under the 12 hours light and 12 hours dark conditions: 50 μmol photons/m^2^/s with Plant cultivation LED light (BC-BML3, Biomedical Science). DNA was extracted with Extrap Soil DNA Kit Plus ver. 2 (Nippon Steel). Total DNA was subjected to library construction with TruSeq DNA PCR Free (350; Illumina) and to 151 bp paired-end sequencing by HiseqX, resulting in 660 million paired-end reads, and to PacBio RSII, with SMRT cell 8Pac V3 and DNA Polymerase Binding Kit P6 v2, in Macrogen, resulting in 1.3 Gb subreads. Total RNA was extracted with Trizol (Sigma) according to the manufacturer’s instruction and was subjected to library construction with TruSeq RNA Sample Prep Kit v2 (Illumina) and 101 bp paired-end sequencing by Hiseq2500, resulting in 107.5 million paired-end reads.

### Genome assembly and construction of gene models

PacBio reads were assembled into contigs using Falcon (ver. 0.7.0) with a length cutoff of 7,000 bp for seed reads and an estimated genome size of 33 Mbp. Genome size estimation was performed on the GenomeScope web server (http://qb.cshl.edu/genomescope/) based on the K-mer frequency distribution of Illumina reads calculated by JellyFish ver. 2.2.6 with a K-mer size of 21. The resultant primary and associate contigs were then subjected to Falcon_unzip (ver. 0.5.0), generating partially haplotype-phased contigs (primary contigs) and fully phased contigs (haplotigs). The assembly was polished using PacBio reads and Quiver program, followed by SNP and short indel error correction using Pilon (ver. 1.2.2) with Illumina reads mapped by BWA (ver. 0.7.15). Indel errors in the vicinity of hetero SNPs were further fixed manually, as they were difficult to be automatically corrected. Contigs derived from plastid and mitochondrial genomes were identified using BLASTN and separated from contigs derived from the nuclear genome.

RNA-seq reads were trimmed under the parameters of ILLUMINACLIP:TruSeq3-PE.fa:2:30:10, LEADING:20 TRAILING:20, SLIDINGWINDOW:4:15 and MINLEN:75 using Trimmomatic (ver. 0.36) (Bolger et al. 2014). The trimmed reads aligned to the assembled contigs using HISAT2 (ver. 2.0.4) (Kim et al. 2015). They were provided to BRAKER2 gene annotation pipeline (ver 2.0.3) as training data to be used for *ab initio* prediction of protein-coding genes. Supplementarily, PASA (ver. 2.3.3) (Haas et al., 2003) was used to generate transcript-based gene models by integrating *de novo* transcriptome assembly and genome-guided assembly using Trinity (ver. 2.5.0) (Grabherr et al. 2011). The genome-guided assembly used the mapping result from HISAT2 with --dta option. TransDecoder (ver. 5.0.2) (Haas et al. 2013) was employed to extract protein coding regions from PASA result with the alignment files from BlastP (ver 2.7.1) (Camacho et al. 2009) against UniRef90 with −evalue 1e-5 option and hmmscan (http://hmmer.org/, ver 3.1b2) against pfam (El-Gebali et al. 2019) database.

The gene models that overlapped with the results from Braker were removed using BlastP with evalue 1e-5 option, and remaining gene models were merged with the Braker gene models to generate the final gene annotation. Transposable elements in the NIES-4239 genome were searched by RepeatMasker (ver. 4.9.0) with using Dfam3.1 and RepBase-20170127 as reference repeat libraries.

The integrity of gene annotation was assessed by BUSCO (ver. 3.0.2) (Simão et al. 2015) and the Eukaryota odb9 (ver. 2) dataset. The manipulation of sam/bam file was used by samtools (ver. 1.9). The sequence files of gene region from gff file were used by gffread. (ver. 0.9.11) (Pertea and Pertea 2020).

Organellar genome annotation was performed as described in previous studies (Kamikawa et al. 2015b; Kamikawa et al. 2018).

Assembled genomes were deposited to DNA Data Bank of Japan under the accession numbers BLYE01000001-BLYE01000234 for the nuclear genome, LC600866 for the mitochondrial genome, and LC600867 for the plastid genome.

### Functional annotation

The predicted protein coding genes were annotated using InterProScan, and RPS-BLAST search was performed against KOG (EuKaryotic Orthologous Groups) database. KO identifiers for Kyoto Encyclopedia of Genes and Genomes (KEGG) metabolic pathways were assigned using KEGG Automatic Annotation Server (KAAS, Moriya et al., 2004). Transporter proteins were annotated with TransportTP (Li et al. 2009). Reference proteome datasets for three photosynthetic diatom species were obtained from the JGI Genome Portal: *Phaeodactylum tricornutum* CCAP 1055/1 v2.0 (Phatr2_bd_unmapped_GeneModels_FilteredModels1_aa.fasta, Phatr2_chromosomes_geneModels_FilteredModels2_aa.fasta, 10,402 protein sequences in total), *Thalassiosira pseudonana* CCMP 1335 (Thaps3_bd_unmapped_GeneModels_FilteredModels1_aa.fasta, Thaps3_chromosomes_geneModels_FilteredModels2_aa.fasta, 11,776 sequences), *Fragilariopsis cylindrus* CCMP 1102 (Fracy1_GeneModels_FilteredModels3_aa.fasta, 21,066 sequences). KEGG and KOG annotation was performed with them in the same manner as NIES-4239. For plastid and mitochondrial genomes assembled in the above procedures, annotation was performed by MFANNOT (insert ref).

### CAZyme annotation

We performed a manual annotation of CAZymes (Lombard et al. 2014) using a combination of BLAST (Altschul et al. 1997) and HMM searches (Mistry et al. 2013), similar to that done previously (Curtis et al. 2012; Cenci et al. 2018). To assess the similarity between the CAZyme family profiles of the two species, we generated heat maps derived from an average linkage hierarchical clustering based on Bray-Curtis dissimilarity matrix distances and Ward’s method (Bray and Curtis 1957; Ward 1963; Cenci et al. 2018). The heat maps were computed with Rstudio software (https://www.rstudio.com/) using vegan in the R package (http://cc.oulu.fi/~jarioksa/softhelp/vegan/html/vegdist.html) (Oksanen 2014) with vegdist and hclust commands.

### Annotation of Cyclins, Cyclin-dependent kinases, bZIP transcription factors, and photoreceptor proteins

Cyclins, cyclin-dependent kinases, transcription factors were retrieved from the results of Pfam annotation (above). In addition, those proteins were surveyed by homology-based search with homologues of *P. tricornutum* and *T. pseudonana* (Huysman et al. 2010; Rayko et al. 2010) as queries. We also specifically surveyed photoreceptor proteins with diel cycle-based expression, according to the list of Annunziata et al. (2019). Detected sequences were again confirmed as proteins of our interest by reciprocal blastP search against non-redundant database of GenBank. For Cyclins, CDKs, and bZIP transcription factors in *N. putrida, P. tricornutum*, and *T. pseudonana* were subjected to MAFFT (version 7.394; Kato and Standley 2013), followed by removal of ambiguously aligned sites by Bioedit (Hall 1999). The resultant datasets were subjected to the maximum likelihood analysis with IQ-TREE (Nguyen et al. 2015) under the LG+Γ+F model with 100 non-parametric bootstrap analyses.

### Annotation of peroxisomal proteins

To identify potential peroxisomal proteins, the protein models of the diatom were screened via KEGG annotation and BLAST analyses. First, annotation of all the protein sequences was performed via GhostKOALA (KEGG Automatic Annotation Server; https://www.kegg.jp/ghostkoala/). Peroxisomal protein candidates were subsequently identified using the “KEGG Mapper – Reconstruct Pathway” tool (https://www.genome.jp/kegg/tool/map_pathway.html). In addition, all the *Nitzschia* proteins were screened for the presence of a peroxisomal targeting signal of type 1 (PTS1, a C-terminal tri-peptide) using a local command line script identifying those entries, which contain the amino acids [SAC][KRHS][LM] within the last three positions of a protein sequence. Detected proteins were then functionally annotated with WebMGA (http://weizhongli-lab.org/metagenomic-analysis/server/kog/) as well as BlastKOALA (https://www.kegg.jp/blastkoala/) and their sequences were further investigated for the presence of additional targeting signals using SignalP 3.0 (http://www.cbs.dtu.dk/services/SignalP-3.0/) and 4.1 (http://www.cbs.dtu.dk/services/SignalP-4.1/), TargetP 1.1 (http://www.cbs.dtu.dk/services/TargetP-1.1/index.php), PredSL (http://aias.biol.uoa.gr/PredSL/), Predotar (https://urgi.versailles.inra.fr/predotar/) and TMHMM 2.0 (http://www.cbs.dtu.dk/services/TMHMM/). To identify factors for peroxisomal biogenesis/maintenance (peroxins) as well as for photorespiration and the glyoxylate cycle, manual BLAST analyses against the *Nitzschia* proteins were conducted using protein queries from the diatoms *Phaeodactylum tricornutum* and *Thalassiosira pseudonana*, the cryptophyte *Guillardia theta* (Davis et al. 2017, Gonzalez et al. 2011, Mix et al. 2018) as well as yeast (peroxin identification) and an e-value cut-off of e-4. For verification, identified candidates were analyzed via reciprocal BlastP against the NCBI nr database (https://blast.ncbi.nlm.nih.gov) and using NCBI Conserved Domain Search (https://www.ncbi.nlm.nih.gov/Structure/cdd/wrpsb.cgi) for identification of conserved domains within the protein sequences. Missing proteins present in peroxisomes of many organisms were especially surveyed in the transcriptome data (see above).

### Annotation of mitochondrial proteins

To identify mitochondrial proteins, we retrieved mitochondrial reference pathways and proteins of *Homo sapiens:* from a stand-alone reactome server (Sidiropoulos et al. 2017; Fabregat et al. 2018a; Fabregat et al. 2018b; Jassal et al. 2020) on our systems, Uniprot (UniProt Consortium 2019) ids of proteins which are annotated that its subcellar locations are ‘mitochondrial matrix’, ‘mitochondrial inner membrane’, ‘mitochondrial outer membrane’ or ‘mitochondrial intermembrane space’ were extracted using the cypher query language. To search and annotate for mitochondrial proteins, we performed PSI-BLAST (Altschul et al. 1990; Altschul et al. 1997; Camacho et al. 2009) search with proteins which have above uniprot ids as queries for the assemblies of genome and transcriptome. As validation of those annotations, we performed three analyses: 1) psi-blast search with above blast-hit sequences for swissprot (UniProt Consortium 2019), 2) KEGG orthology (KO) number assignment by kofamscan (Aramaki et al. 2020), 3) mitochondrial import signal analysis by Mitofates (Fukasawa et al. 2015) and Nommpred (Kume et al. 2018). We manually refined our annotations based on these results. Missing proteins present in mitochondria of many organisms and/or in each metabolic pathway were especially surveyed in the transcriptome data (see above).

### Annotation of plastid proteins

To identify plastid proteins, protein sequences were subjected to SignalP4.0 (Petersen et al. 2011) followed by ASAFind (Gruber et al. 2015). Functions and related metabolisms for proteins predicted to be localized in the plastid were estimated by their KEGG ID (see above) and KEGG mapper (Kanehisa et al. 2012). Missing proteins in each metablic pathway were especially surveyed in the transcriptome data (see above). Plastid proteins in the photosynthetic diatoms *Phaeodactylum tricornutum* and *Thalassiosira pseudonana* have been already published in Gruber et al. (2005). We retrieved them and clustered with the plastid proteins of *N. putrida* as orthogroups by Orthofinder with default settings (Emms and Kelly 2015).

### Secretome analysis

We detected protein sequences with N-terminal signal peptides and no internal transmembrane domain by evaluation with signalP4.0 (Petersen et al. 2011) and HMTMM (Sonnhammer et al., 1998; Krogh et al., 2001), respectively. Those sequences highly likely comprise secretome and plastid sequences. The sequences were clustered by TribeMCL (Enlight et al. 2002) into Tribes, with homology under e-30 criterion. If a tribe comprises sequences predicted to be localized in plastids with the “high confidence” category by ASAFind (Gruber et al. 2015), more than or equal to those predicted to be non-plastidal or localized in plastids with the “low confidence” category, we removed the tribe. Functional categorization was performed with domain annotation by Pfam, and we also removed tribes if the included sequences were predicted to have a domain for apparent organellar proteins such as plastid protein translocons, plastid heme biosynthesis, components of photosystems, organellar transporters, and lysosomal proteins by KO definition and/or Pfam. To evaluate whether the above procedure to detect secreted proteins is appropriate for diatoms, we made a benchmark dataset of diatoms including 21 secreted proteins experimentally confirmed (Bruckner et al. 2011; Buhmann et al. 2016; Lachnit et al. 2019; Dell’Aquila et al. 2020) and 62 of their homologues, 182 plastid proteins, 29 mitochondrial proteins, and 29 proteins localized in other compartments such as cytosol and nucleus. From the benchmark set, our method identified 77 secreted proteins as secretome proteins, indicating 92.8% recovery rate. But no non-secreted proteins was not identified as secretome proteins in our benchmark set, suggesting that our secretome dataset for whole protein data in the four diatoms less likely comprise high propotion of non-secreted proteins. In addition, the benchmark dataset included 17 secreted protein sequences, experimentally verified, of *Phaeodactyum*, and of them, 13 protein sequences were retrieved by our genome-wide secretome analysis. Four proteins not identified were appeared to lack signal peptides detactable either SignalP4.0 and SignalP3.0, indicating those sequences might be secreted by signal peptide-independnet ways.

Difference in the distributions of the number of protein sequences in each secretome tribe among species was tested by the Wilcoxon signed rank test. *p*-values for three pairs of comparisons that include *N. putrida* as a counterpart were adjusted by the Benjamini–Hochberg procedure for multiple-testing correction. The Wilcoxon signed rank test and the Benjamini–Hochberg procedure were conducted by using ‘wilcoxon’ function implemented in SciPy library (version 1.4.1) and ‘multipletests’ function implemented in Statsmodels library (version 0.11.1) for Python, respectively.

For LRR, additional domains were also searched by NCBI Conserved Domain Search. All the possible secreted LRR proteins in *N. putrida*, *P. tricornutum*, and *T. pseudonana* were subjected to MAFFT (version 7.394; Kato and Standley 2013) and ambiguously aligned sites were removed by Bioedit (Hall 1999). The resultant dataset was subjected to IQ-TREE (Nguyen et al. 2015) under the LG+Γ+F model with 100 non-parametric bootstrap analyses.

### Horizontal gene transfers

Total *N. putrida* protein sequences were subjected to Orthofinder with default settings (Emms and Kelly 2015) together with whole protein sets of the photosynthetic diatom genomes *P. tricornutum*, *F. cylindrus*, and *T. pseudonana*, resulting in 4,920 proteins unique to *N. putrida*. The 4,920 proteins were subjected to similarity search using BLASTP (BLAST+ 2.6.0) against GenBank non-redundant protein sequence database and protein sequence database provided by the Marine Microbial Eukaryote Transcriptome Sequencing Project (https://github.com/dib-lab/dib-MMETSP; Johnson et al. 2019) with an e-value threshold of 1.0E-15. BLASTP hits from two databases were combined and sorted by their bit-scores. Top ten hits with higher bit-scores for each query sequence were then inspected their taxonomic composition with two criteria: 1) if those hits do not include any diatom sequences, namely, sequences from diatom species or dinoflagellate species bearing diatom-derived plastids, 2) if the ten hits do not include the diatom sequences except sequences from diatoms belonging to genus *Nitzschia*. Query sequences, whose top ten BLASTP hits match the two criteria, were considered as initial candidates for 1) genes derived from *N. putrida* specific horizontal gene transfer (HGT) event and 2) *Nitzchia*-lineage-specific HGT, respectively. Each of candidates with HGT origin was aligned together with all of the hit sequences in aforementioned BLASTP analyses by MAFFT (version 7.394; Katoh and Standley 2013). After ambiguously aligned positions were removed by BMGE (version 1.12; Criscuolo and Gribaldo 2010), each single-gene alignment was subjected to a maximum-likelihood (ML) phylogenetic analysis by IQ-TREE (Nguyen et al. 2015) under LG+Γ+F model. The statistical supports for the bipartitions in ML trees were assessed by a nonparametric ML bootstrap analysis (100 replicates) under the same substitution model used for the ML trees. All topologies of ML trees for both of *N. putrida*-specific HGT and *Nitzchia*-lineage-specific HGT candidates were manually assessed.

### Transcriptome analyses

The cells of *N. putrida* NIES-4239 cultivated under the 12 hours light and 12 hours dark conditions under the same condition as described above were then cultivated under the dark conditions for 48 hours. The cells possibly acclimated to the dark condition were transferred to the fresh liquid medium and incuvated under the 12 hours light and 12 hours dark condition for 25 hours in which the last one hour was of the second light term. RNA was extracted every 4 hours at the one hour, 5 hours, and 9 hours after initiation of the first light condition, one hour, 5 hours, and 9 hours after initiation of the dark condition, and one hour after initiation of the second light term. The cultivation and RNA extraction were performed twice, samples which were called Replicates A and B. The extracted total RNAs were subjected to 151 bp paired-end sequencing by NextSeq 500 according to the manufacturer’s instruction in Bioengineering Lab, Japan, resulting in 9.8-13 million paired-end reads for each sample. Adapter trimming and quality filtering were performed with fastX toolkit (http://hannonlab.cshl.edu/fastx_toolkit/). In quality filtering, reads with quality scores > 20 for at least 75% of their length were retained, resulting in 7.3-11.1 million paired-end reads. To obtain gene expression scores, one side of the paired-end reads was mapped to the reference by Bowtie2 ver. 2.3.4.1 (Langmead and Salzberg 2012). SAMtools ver. 1.8 (Li et al. 2009), BEDtools ver. 2.19.1 (Quinlan and Hall 2010), and R ver. 3.5.3 (Ihaka and Gentleman 1996) were used to calculate the reads per kilobase of exon per million mapped reads (RPKM). To extract the reproducible change in transcriptome of Replicate A and B, the genes with low value of Pearson’s correlation coefficient between two biological replicates (≦ 0.9) were omitted, resulting in 1,971 genes. To investigate expression patterns of 1,971 genes, the values of RPKM + 1 of each gene were transformed to log2 and centered by median values, and *k*-means clustering (*k* = 8) was performed by using Cluster 3.0 (de Hoon et al. 2004) based on Spearman correlation and complete linkage. The clustered expression patterns were visualized by Java TreeView (Saldanha 2004) and R.

The transcriptome data were deposited in DNA Data Bank of Japan under the accession number PRJDB11016.

### Divergence time estimation

All sequences of interest in diatoms were first clustered using cd-hit/4.6.8, removing sequences of 100% alignment and clustering at low identity. The largest clusters for each gene family was then aligned using PRANK 170427 (Löytynoja, 2014), poorly aligned sequences were removed with TrimAl 1.2 (Capella-Gutiérrez, Silla-Martínez and Gabaldón, 2009) and put into gene blocks to remove gapped columns with Gblocks v0.91b (Castresana 2000; Talavera and Castresana 2007). Aligned sequences containing all species with available data (N = 1 to 4) were then analysed using phylogenetic analyses. Divergence estimates were obtained using Bayesian Markov Chain Monte Carlo (MCMC) analyse is implemented in Beast 2.5.0 (Bouckaert et al. 2019). The analysis was carried out using a relaxed molecular clock approach with an uncorrected log-normal distribution model of rate of variation, the HKY substitution model, four gamma categories and a Yule model of speciation, for each gene family 5 runs were carried out with 20 million MCMC generations sampled every 1,000^th^ generation. The results from all runs were combined and summarised using LOGcombiner v1.10.4, these were checked for convergence using Tracer v1.7.1 (Rambaut, 2009). A maximum clade credibility tree was generated using Tree Annotator v1.10.4 and was graphically visualised using FigTreev 1.4.3 software (Rambaut, 2012). The Phylogenetic Maximum Clade Credibility (MCC) trees were used to determine the rate of “speciation” (or more accurately, the expansion) within a given gene family. The R package TESS (Höhna, May and Moore, 2016) was used for this analysis with the mass extinction turned off, measuring only for the speciation rate (expansion rate), using the phylogenetic trees produced in the divergence time estimations.

Likelihood ratio tests (2ΔL) were performed by PAML 4 (Yang 2007) between 3 Codeml site model pairs (M0/M3; M1a/M2a and M7/M8), with the first model representing a site model without positive selection and the second a model that allows a proportion of sites to be under positive selection. In all 3 cases the model with positive selection was a significantly better fit to the data. For the M2a and M8 models, several sites were predicted to be positively selected by Bayes Empirical Bayes analysis (Yang et al. 2005). The sites 457 and 460 were identified by both models with high confidence (> 95%) and 457 by M8.

### Analysis of lipids, fatty acids, and quinones/quinols

Crude lipids were extracted from the cells by the method of Bligh and Dyer (1959). The lipid fraction was evaporated, and then the residue was heated at 90°C for 2 h with 2 mL of 5% (w/w) HCI-methanol to obtain fatty acid methyl esters. The methanol solution was extracted with 2 mL of n-hexane twice. The layer of n-hexane was concentrated to a minimum volume for use in gas-chromatographic analysis. Gas chromatography was performed, according to Mitani et al. (2017), with a fused-silica capillary column (0.25 mm internal diameter x 50 m; ChrompackCp-Sil 88, Agilent Technologies Inc., USA) with the oven temperature was increased from 150 to 210°C at 5°C min^-1^. The fatty acid composition was calculated by a Chromatopac C-R8A data processor (Shimadzu Corp., Kyoto, Japan). Each fatty acid was identified by comparing the retention time with those of known standards. Thin layer chromatograph (silica gel 60) of the crude lipids developed by hexane/diethyl ether/acetate (80:20:1 v/v/v). Lipids were visualized under UV light at 365 nm after spraying the plate with 0.01% (w/v) primulin in 80% (v/v) acetone (Wright 1971). Two-dimensional thin layer chromatograph of the polar lipids was performed according to Sato (1991) with the following solvent systems: first dimension, acetone/benzene/methanol/water (8:3:2:1 v/v/v/v); second dimension, chloroform/methanol/28% ammonium (13:7:1 v/v/v). Each spot of polar lipids was detected as described above. Each lipid was identified by spraying the plate with a reagent specific to each lipid class (Allen and Good 1971). Each spot on a plate was scraped off, and then subjected to gas-chromatographic analysis and fatty acid identification as described above.

Quinone/quinol extraction and detection was performed as described in Kayama et al. (2020). In this analyses, we treated the quinone/quinol extract with ferric chloride (final concentration, 1.2 mM) before the analysis for oxidation of total quinones and quinols, to convert them to quinone forms for higher detection sensitivity according to Kayama et al. (2020). We could identified only ubiquinone but no plastoquinone from the cells of *N. putrida* NIES-4239 cultivated as described above for genome sequencing, supporting lack of the complete pathway of plastoquinol synthesis (Fig. 1).

## Acknowledgements

This work was in part supported by JSPS grant-in-aid for Scientific Research (B) awarded to RK (19H03274), TN (20H03305), and GT (17H03723) and by JSPS grantin-aid for Scientific Research (A) YK (18H03743). This work was also supported by JSPS grant-in-aid for for Scientific Research on Innovative Areas, Platform for Advanced Genome Science to RK, TM, MS, YT, and YN (16H06279). We thank Ms. F. Hayashi for the manual correction of the genome assembly and the gene annotation for the secretome. TM, CvO, AT, KS, and SS acknowledge funding for this work from NERC (NE/R000883/1) and the School of Environmental Sciences at the University of East Anglia, UK. S.K. and J.W. were supported by the Gatsby Charitable Foundation and Biotechnology and Biological Sciences Research Council (BBSRC). Computational work was in part performed at the SuperComputer System, Institute for Chemical Research, Kyoto University.

## Author Contributions

RK, GT, TM, and YN conceived this research. RK, YT, TM, MS, GT, and YN sequenced, assembled, and annotated the genome. RK identified plastid metabolisms. RK and TN detected HGT candidates. RO, RK, and SM performed comparative transcriptome analyses. UC, BH, and VL searched and annotated CAZymes. DM and RK identified peroxisome metabolisms. SS, KS, AT, CO, and TM performed the coalescence analyses. KO and MK detected and identified lipids and fatty acids. KK and RK identified mitochondrial metabolisms. RK, KI, and SM isolated and identified the diatom. MK, TA, KI, HM performed growth experiments. MK and YK performed quinone detection. RK, JW, SK, TM, and MS identified secretomes. RK, CVO, and TM wrote the manuscript, and all the authors commented and approved the final version.

