## Supplementary figures and tables for "Genome evolution of a non-parasitic secondary heterotroph, the diatom *Nitzschia putrida*"

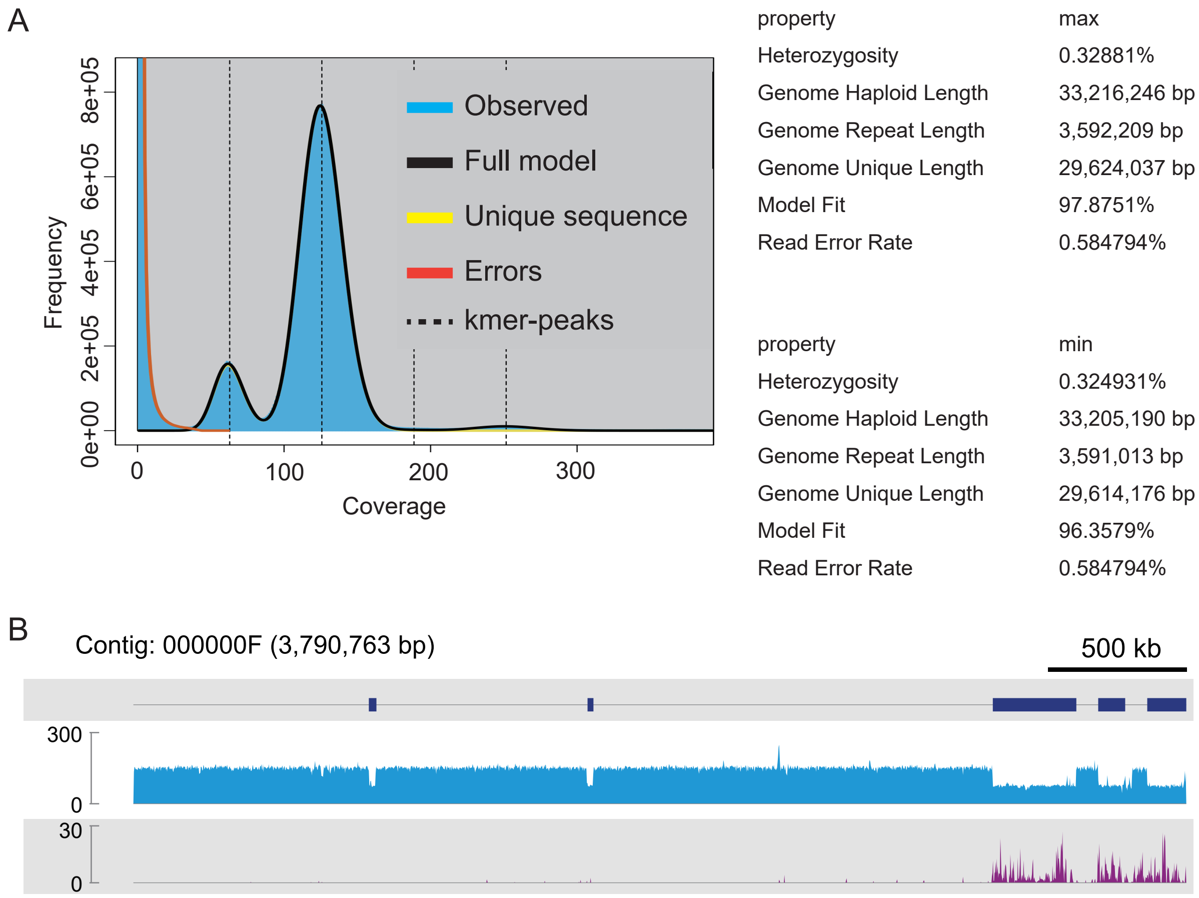

| property | max |
|---|---|
| Heterozygosity | 0.32881% |
| Genome Haploid Length | 33,216,246 bp |
| Genome Repeat Length | 3,592,209 bp |
| Genome Unique Length | 29,624,037 bp |
| Model Fit | 97.8751% |
| Read Error Rate | 0.584794% |

| property | min |
|---|---|
| Heterozygosity | 0.324931% |
| Genome Haploid Length | 33,205,190 bp |
| Genome Repeat Length | 3,591,013 bp |
| Genome Unique Length | 29,614,176 bp |
| Model Fit | 96.3579% |
| Read Error Rate | 0.584794% |

**Supplementary Fig. S1 Genomic features showing diploid genome structure with a portion of heterozygosity.** A. GenomeScope analysis. B. Read mapping of Illumina short reads onto the primary contig 000000F. Dark blue boxes show primary contig regions corresponding to haplotigs. Light blue shows the read depth, while purple shows the number of variants, such as SNPs and Indels, per 1,000 bp.

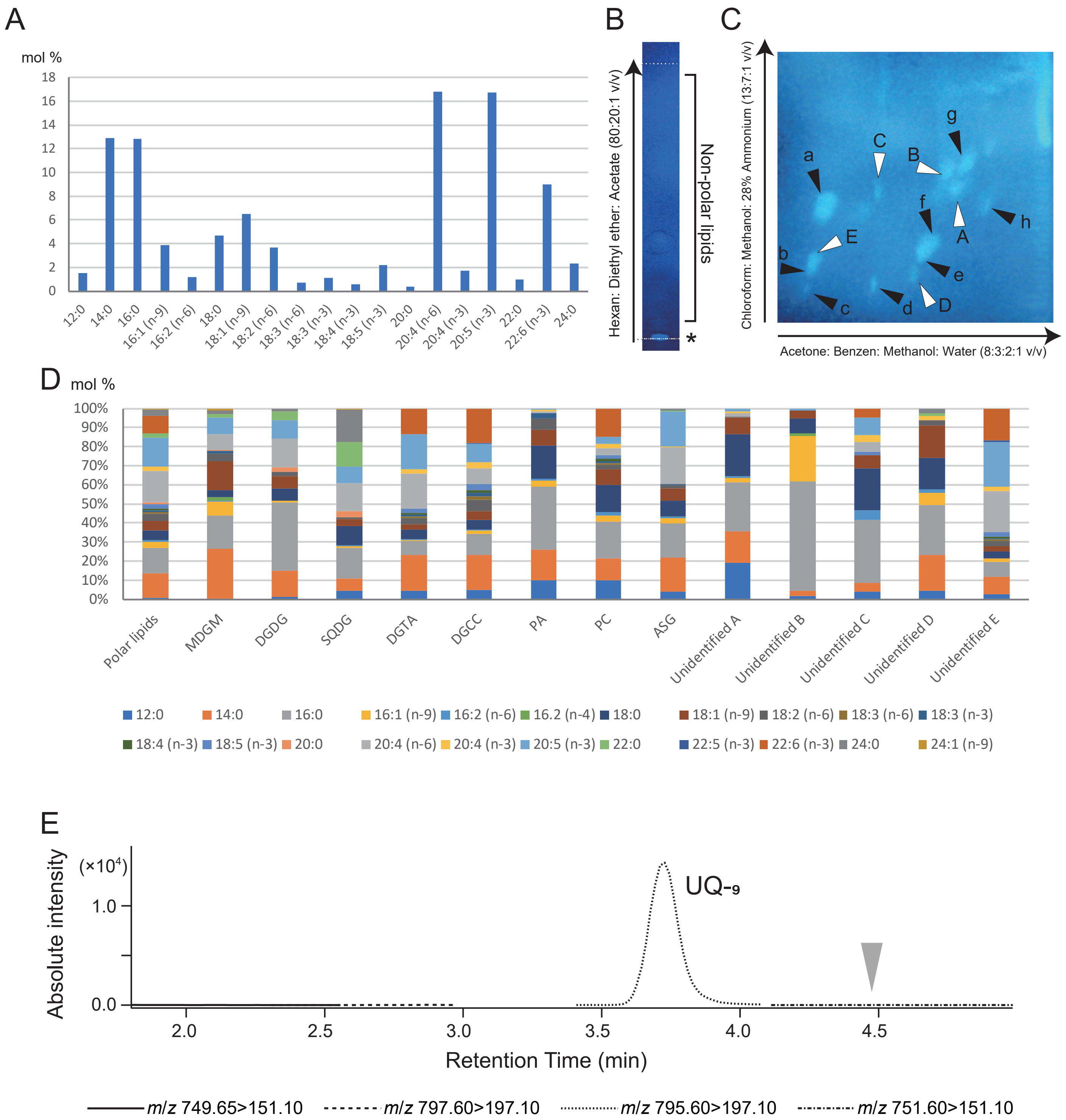

**Supplementary Fig. S2 Fatty acids, lipids, and quinones in Nitzschia putrida NIES-4239.** A. Fatty acid composition of the crude lipid extract. Crude lipids were methylated after extraction and subjected to gas chromatography. B. Thin layer chromatograph (silica gel 60) of the crude lipids developed by hexane/diethyl ether/acetate (80:20:1 v/v/v). Lipids were visualized under UV light at 365 nm after spraying the plate with 0.01% (w/v) primulin in 80% (v/v) acetone. Although lipids of 19 mg dry weight cells were applied, explicit spots of neutral lipids were not appeared. Polar lipids were indicated by an asterisk. C. Two-dimensional thin layer chromatograph of the polar lipids. Solvent systems: first dimension,acetone/benzene/methanol/water (8:3:2:1 v/v/v/v); second dimension, chloroform/methanol/28% ammonium (13:7:1 v/v/v). Each spot of polar lipids was detected as described above. Each spot of lipids was detected as polar lipids by primuline under UV. Each lipid was identified by the retardation factors of standard materials and the color reaction. a: Diacylglycerylhydroxymethyltrimethyl-β-alanine, b: Diacylglyceryl carboxyhydroxymethylcholine, c: Phosphatidylcholine, d: Phosphatidic acid, e: Digalactosyl diacylglycerol, f: Sulfoquivonosyl diacylglycerol, g: Monogalactosyl diacylglycerol, h: Acylated steryl glycoside, Open arrowheads A - E: unidentified lipids A – E. We could not detect phosphatidyl glycerol (PG), supporting absence of PG biosynthesis. D. Fatty acid compositions in each spot appeared in C. E. LC-MS/MS chromatograms (multiple reaction monitoring mode) of the acetone extract after oxidative treatment with ferric chloride. The grey arrow indicates the retention time for plastoquinone peak in this system (Kayama et al. 2020).

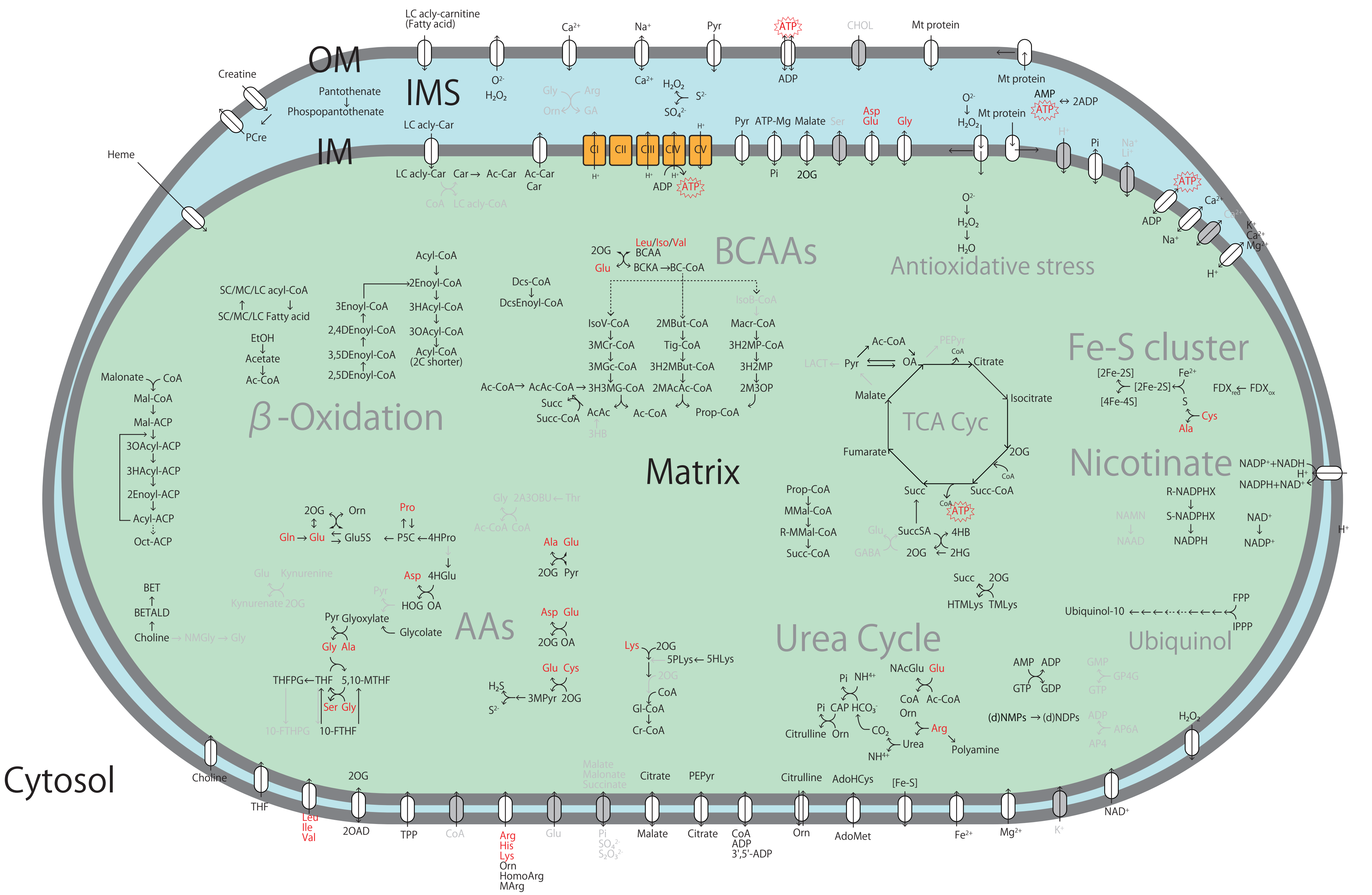

OM
IMS
IM
Matrix
Cytosol
LC acly-carnitine
(Fatty acid)
Ca2+
Na+
Pyr
ATP
CHOL
Mt protein
Creatine
PCre
Pantothenate
Phospopantothenate
O2-
H2O2
Gly
Arg
Orn
GA
H2O2
S2-
SO42-
ADP
AMP
2ADP
LC acly-Car
Car
Ac-Car
CoA
LC acly-CoA
Heme
CI
CII
CIII
CIV
CV
H+
Pyr ATP-Mg Malate Ser
Asp
Glu
Pi
2OG
Li+
K+
Mg2+
BCAAs
Leu/Iso/Val
BCAA
BCKA
BC-CoA
IsoV-CoA
3MCr-CoA
3MGc-CoA
3H3MG-CoA
2MBut-CoA
Tig-CoA
3H2MBut-CoA
2MAcAc-CoA
IsoB-CoA
Macr-CoA
3H2MP-CoA
3H2MP
2M3OP
Prop-CoA
Ac-CoA
AcAc-CoA
AcAc
Succ
Succ-CoA
3HB
Antioxidative stress
H2O
SC/MC/LC acyl-CoA
SC/MC/LC Fatty acid
EtOH
Acetate
Acyl-CoA
2Enoyl-CoA
3Enoyl-CoA
3HAcyl-CoA
2,4DEnoyl-CoA
3OAcyl-CoA
3,5DEnoyl-CoA
Acyl-CoA (2C shorter)
2,5DEnoyl-CoA
Dcs-CoA
DcsEnoyl-CoA
β-Oxidation
Malonate
Mal-CoA
Mal-ACP
3OAcyl-ACP
3HAcyl-ACP
2Enoyl-ACP
Acyl-ACP
Oct-ACP
TCA Cyc
LACT
OA
PEPyr
Citrate
Isocitrate
Malate
Fumarate
Succ-CoA
Fe-S cluster
[2Fe-2S]
[4Fe-4S]
Fe2+
S
Cys
Ala
FDXred
FDXox
Nicotinate
NADP++NADH
NADPH+NAD+
R-NADPHX
S-NADPHX
NADPH
NAMN
NAAD
NAD+
NADP+
Prop-CoA
MMal-CoA
R-MMal-CoA
SuccSA
4HB
2HG
GABA
HTMLys
TMLys
FPP
IPPP
Ubiquinol-10
Ubiquinol
Urea Cycle
AAs
Gln
Glu5S
Pro
P5C
4HPro
Thr
2A3OBU
Ala
Pyr
Kynurenine
Kynurenate
4HGlu
HOG
Glyoxylate
Glycolate
BET
BETALD
Choline
NMGly
THFPG
THF
5,10-MTHF
10-FTHPG
10-FTHF
H2S
3MPyr
Lys
5PLys
5HLys
Gl-CoA
Cr-CoA
NACGlu
CAP
HCO3-
CO2
Citrulline
Urea
Polyamine
NH4+
AMP ADP
GTP GDP
GMP
GP4G
(d)NMPs
(d)NDPs
AP6A
AP4
2OAD
TPP
His
HomoArg
MArg
Leu
Ile
Val
SO42-
S2O32-
Malonate
Succinate
ADP
3',5'-ADP
AdoHcys
AdoMet
[Fe-S]

**Supplementary Fig. S3 Mitochondrial metabolisms predicted from the genome data of *Nitzschia putrida.***

Overview of the metabolic pathway of the mitochondria based on in silico analyses. OM, IMS, and IM: outer membrane, intermembrane space, and inner membrane, respectively. Arrows indicates metabolic reactions. Essential amino acids and ATP are highlighted in red. Ovals indicates membrane translocators. Undetectable reactions and their involving substrates are shown in gray. Abbreviations:

10-FTHF: 10-Formyltetrahydrofolate, 10-FTHPG: 10-formyltetrahydrofolate polyglutamate, 2:4DEnoyl-CoA: 2:4-dienoyl-CoA, 2:5DEnoyl-CoA: 2:5-dienoyl-CoA, 2A3OBU: 2-Amino-3-oxobutanoic acid, 2Enoyl-ACP: 2-enoyl-ACP, 2Enoyl-CoA: 2-enoyl-CoA, 2HG: 2-Hydroxyglutarate, 2M3OP: 2-Methyl-3-oxopropanoate, 2MAcAc-CoA: 2-Methylacetoacetyl-CoA, 2MBut-CoA: 2-Methylbutanoyl-CoA, 2OAD: 2-oxoadipate, 2OG: 2-oxoglutarate, 3:5DEnoyl-CoA: 3:5-dienoyl-CoA, 3Enoyl-CoA: 3-enoyl-CoA, 3H2MBut-CoA: 3-Hydroxy-2-methylbutanoyl-CoA, 3H2MP: 3-Hydroxy-2-methylpropanoate, 3H2MP-CoA: 3-hydroxy-2-methylpropanoyl-CoA, 3H3MGl-CoA: 3-Hydroxy-3-methylglutaryl-CoA, 3HAcyl-ACP: 3-hydroxyacyl-ACP, 3HAcyl-CoA: 3-hydroxyacyl-CoA, 3HB: 3-Hydroxybutanate, 3MCr-CoA: 3-methylcrotonoyl-CoA, 3MGc-CoA: 3-Methylglutaconyl-CoA, 3MPyr: 3-Mercaptopyruvate, 3OAcyl-ACP: 3-oxoacyl-ACP, 3OAcyl-CoA: 3-oxoacyl-CoA, 4HB: 4-Hydroxybutanoate, 4HGlu: 4-Hydroxy-L-glutamate, 4HPro: 4-Hydroxyproline, 5:10-MTHF: 5:10-methylenetetrahydrofolate, 5HLys: 5-Hydroxy-L-lysine, 5PLys: 5-Phosphooxy-L-lysine, Ac-Car: Acetylcarnitine, Ac-CoA: Acetyl-CoA, AcAc: Acetoacetate, AcAc-CoA: Acetoacetyl-CoA, ACP: Acyl-carrier protein, AdoHCys: S-Adenosyl-L-homocysteine, AdoMet: S-Adenosyl-L-methionine, AP4: Adenosine tetraphosphate, AP6A: Diadenosine hexaphosphate, BC-CoA: Branched-chain acyl-CoA, BCAA: Branched-chain amino acid, BCKA: Branched-chain ketoacid (2-oxoacid), CAP: Carbamoyl phosphate, Car: L-Carnitine, CHOL: Cholesterol, CI - CV: Complex I - V, CoA: Coenzyme A, Cr-CoA: Crotonoyl-CoA, Dcs-CoA: Docosanoyl-CoA, DcsEnoyl-CoA: Docos-2-enoyl-CoA, FDX: Ferredoxin, FPP: Farnesyl diphosphate, GA: Guanidinoacetate, Gl-CoA: Glutaryl-CoA, Glu5S: Glutamate 5-semialdehyde, GP4G: Diguanosine tetraphosphate, HOG: 4-Hydroxy-2-oxoglutarate, HomoArg: Homoarginine, HTMLys: Hydroxy-trimethyllysine, IPPP: Isopentenyl diphosphate, IsoB-CoA: Isobutyryl-CoA, IsoV-CoA: Isovaleryl-CoA, LACT: Lactate, LC acly-Car: Long-chain acyl-Carnitine, LC acyl-CoA: Long-chain acyl-CoA, Macr-CoA: Methacrylyl-CoA, Mal-ACP: Malonyl-ACP, Mal-CoA: Malonyl-CoA, MArg: Methylarginine, MC acyl-CoA: Medium-chain acyl-CoA, MMal-CoA: Methylmalonyl-CoA, NAAD: Nicotinamide adenine dinucleotide, NAcGlu: N-Acetyl-L-glutamate, NAD: Nicotinamide adenine dinucleotide, NAMN: Nicotinamide mononucleotide, NMGly: N-Methacryloylglycinamide, OA: Oxaloacetate, Oct-ACP: Octanoyl-ACP, Orn: Ornithine, P5C: Pyrroline-5-carboxylate, PCre: Phosphocreatine, PEPyr: Phosphoenolpyruvate, Pi: Phosphate, Prop-CoA: Propanoyl-CoA, Pyr: Pyruvate, R-MMal-CoA: R-Methylmalonyl-CoA, R-NADPHX: R-NADPH-hydrate, S-NADPHX: S-NADPH-hydrate, SC acyl-CoA: Short-chain acyl-CoA, Succ: Succinate, Succ-CoA: Succinyl-CoA, SuccSA: Succinate semialdehyde, THF: Tetrahydrofolate, THFPG: Tetrahydrofolate polyglutamate, Tig-CoA: Tiglyl-CoA, TMLys: Trimethyllysine, TPP: Thiamine diphosphate.

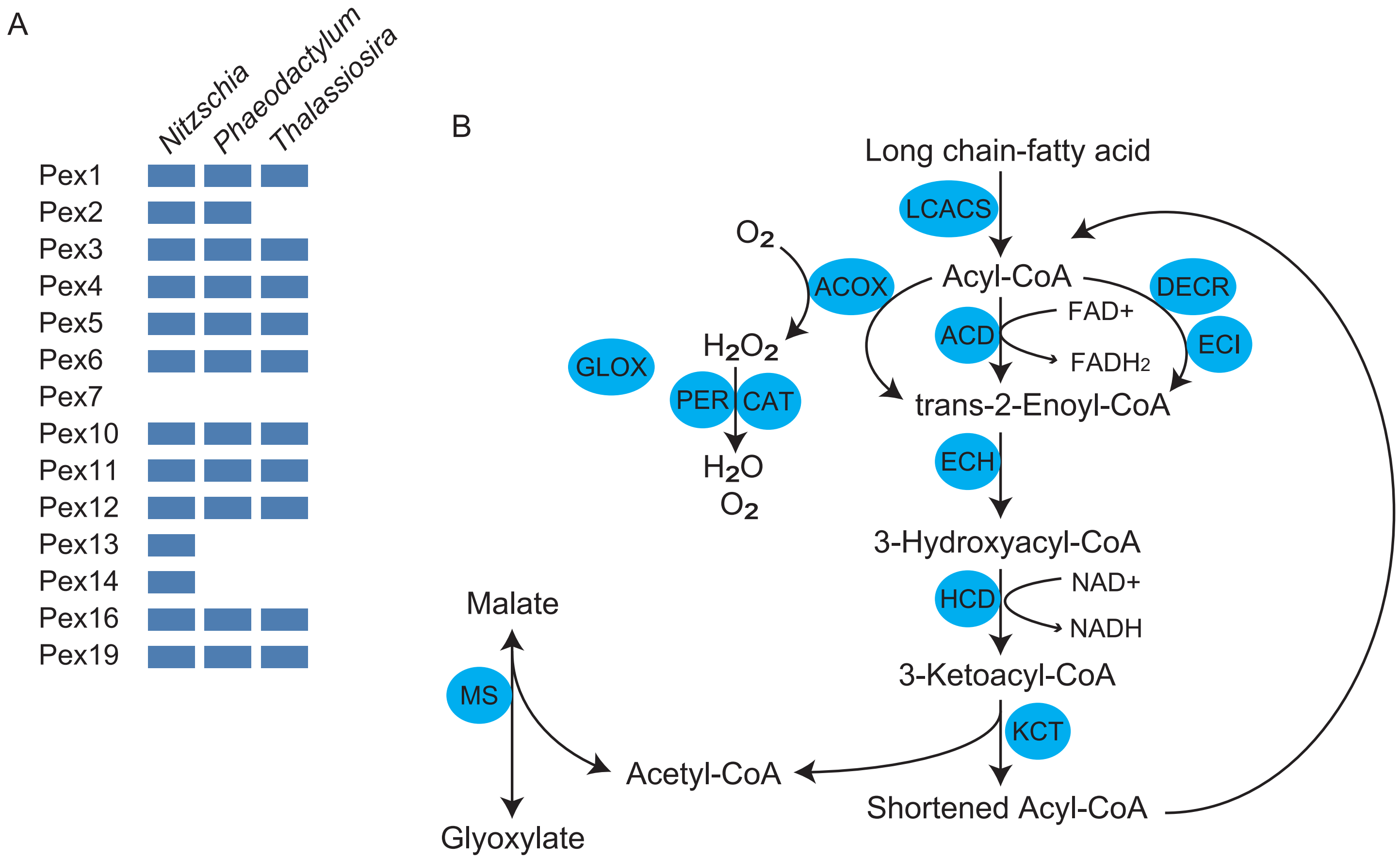

**Supplementary Fig. S4 Peroxisomal metabolisms and peroxin compositions predicted from the genome data of *Nitzschia putrida.*** A. Distribution of Pex factors. Genes identified in the diatom genomes are shown by blue boxes. *Nitzschia*: *N. putrida*, *Phaeodactylum*: *P. tricornutum*, *Thalassiosira*: *T. pseudonana*. B. Predicted peroxisomal metabolism in *N. putrida*. Arrows show metabolic reactions while grey circles show responsible enzymes detected in the genome. ACD: Acyl-CoA dehydrogenase, ACOX: acyl-CoA oxidase, CAT: catalase, DECR: peroxisomal 2,4-dienoyl-CoA reductase, ECH: 3-alpha,7-alpha,12-alpha-trihydroxy-5-beta-cholest-24-enoyl-CoA hydratase, ECI: peroxisomal 3,2-trans-enoyl-CoA isomerase, GLOX: glutathione-dependent disulfide-bond oxidoreductase, HCD: 3-hydroxyacyl-CoA dehydrogenase, KCT: beta-ketoacyl-coa thiolase, LCACS: long-chain acyl-CoA synthetase, MS: malate synthase, PER: peroxiredoxin.

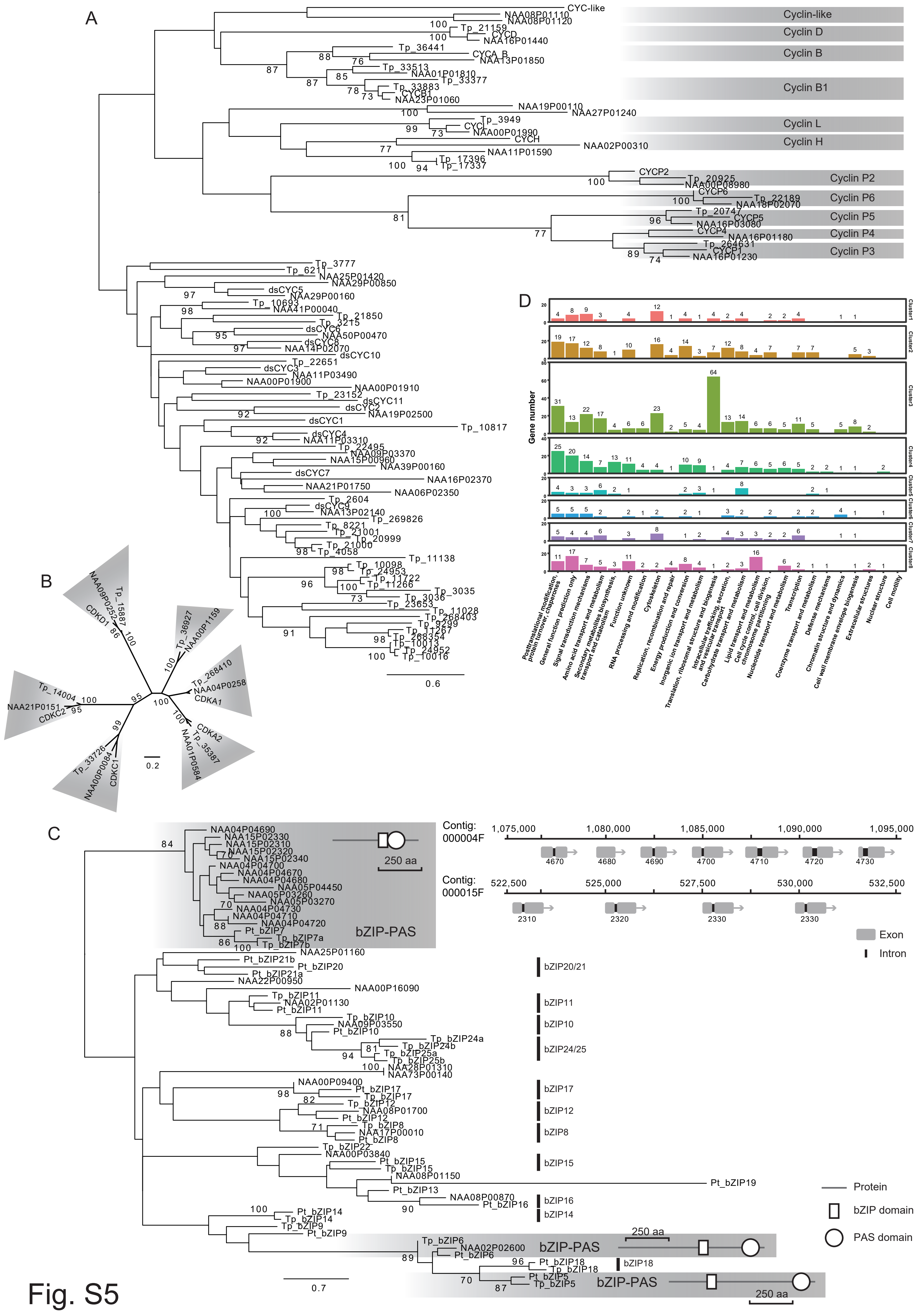

A
Cyclin-like
Cyclin D
Cyclin B
Cyclin B1
Cyclin L
Cyclin H
Cyclin P2
Cyclin P6
Cyclin P5
Cyclin P4
Cyclin P3
0.6
B
0.2
C
bZIP-PAS
250 aa
Contig: 000004F
1,075,000
1,080,000
1,085,000
1,090,000
1,095,000
Contig: 000015F
522,500
525,000
527,500
530,000
532,500
Exon
Intron
bZIP20/21
bZIP11
bZIP10
bZIP24/25
bZIP17
bZIP12
bZIP8
bZIP15
bZIP16
bZIP14
bZIP18
Protein
bZIP domain
PAS domain
0.7
D
Gene number
Cluster1
Cluster2
Cluster3
Cluster4
Cluster5
Cluster6
Cluster7
Cluster8
Posttranslational modification, protein turnover, chaperones
General function prediction only
Signal transduction mechanisms
Amino acid transport and metabolism
Secondary metabolites biosynthesis, transport and catabolism
Function unknown
RNA processing and modification
Cytoskeleton
Replication, recombination and repair
Energy production and conversion
Inorganic ion transport and metabolism
Translation, ribosomal structure and biogenesis
Intracellular trafficking, secretion, and vesicular transport
Carbohydrate transport and metabolism
Lipid transport and metabolism
Cell cycle control, cell division, chromosome partitioning
Nucleotide transport and metabolism
Coenzyme transport and metabolism
Transcription
Defense mechanisms
Chromatin structure and dynamics
Cell wall/membrane/envelope biogenesis
Extracellular structures
Nuclear structure
Cell motility

Fig. S5

**Supplementary Fig. S5 Phylogenetic diversity of genes for cell cycle regulation and transcription and functional annotation of differentially expressed genes.** A. Cyclin gene phylogeny of diatoms. CYC and dsCYC are cyclin and diatom-specific cyclins of *Phaeodactylum tricornutum*, respectively (Huysman et al. 2010). Tp: *Thalassiosira pseudonana*. Numbers of *Thalassiosira* homologues are of protein ID in *T. pseudonana* genome v3 in JGI. B. Cyclin-dependent kinase phylogeny of diatoms. CDKs are *Phaeodactylum* homologues (Huysman et al. 2010). C. bZIP phylogeny in diatoms. Tp_bZIP and Pt_bZIP are of bZIP homologues in *T. pseudonana* and *P. tricornutum* according to Rayko et al. (2010). Domain structures are depicted for bZIP homologues with PAS domain (bZIP5-bZIP7). Positions of genes for duplicated and diversified homologues of bZIP7 with PAS domain in *N. putrida* are also depicted. Many of them are of tandem duplication. D. KOG-based functional annotation of differentially expressed genes clustered in Fig. 2D.

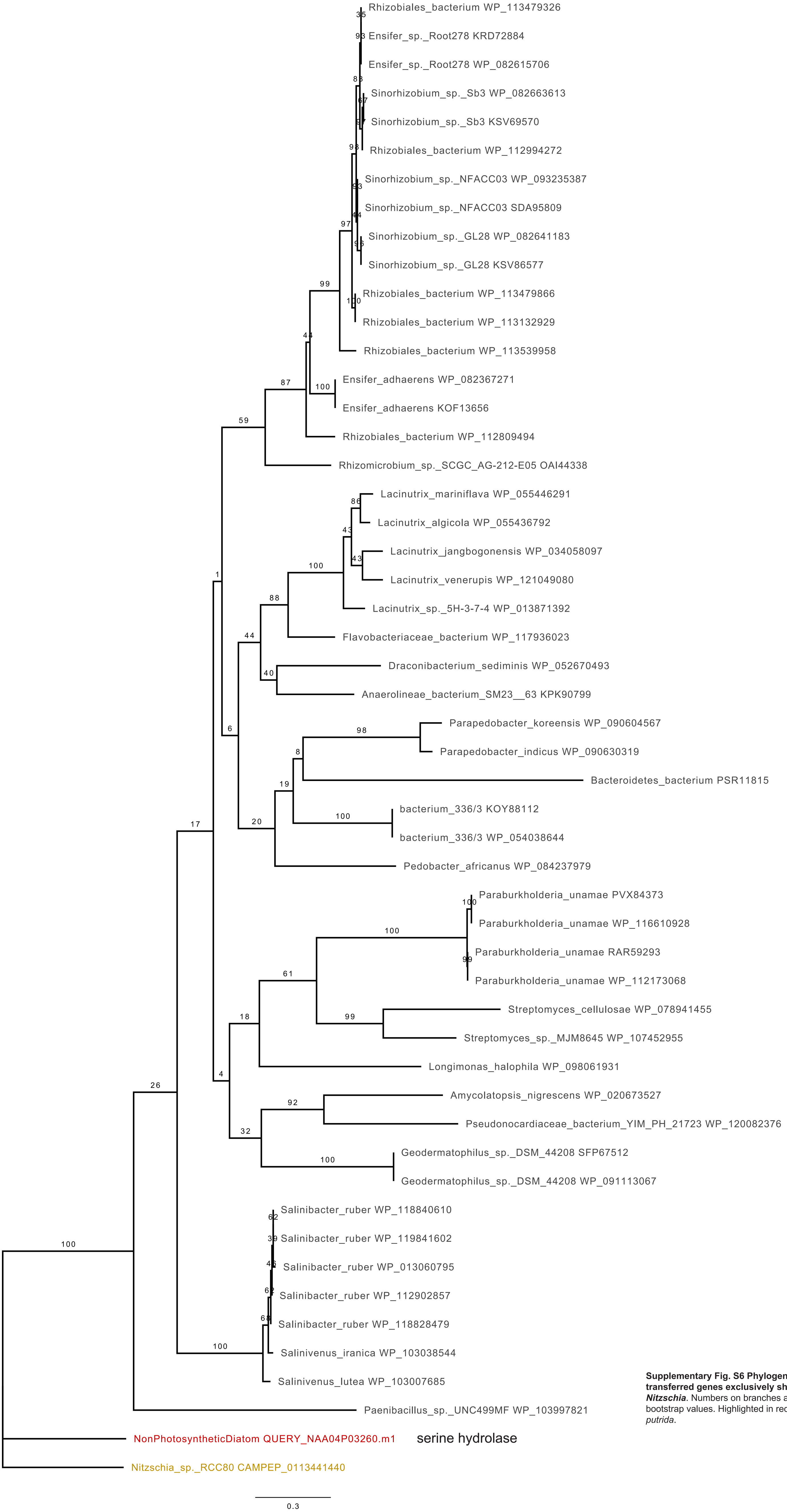

**Supplementary Fig. S6 Phylogeny of three laterally transferred genes exclusively shared in the genus *Nitzschia*.** Numbers on branches are maximum likelihood bootstrap values. Highlighted in red are sequences of *N. putrida*.

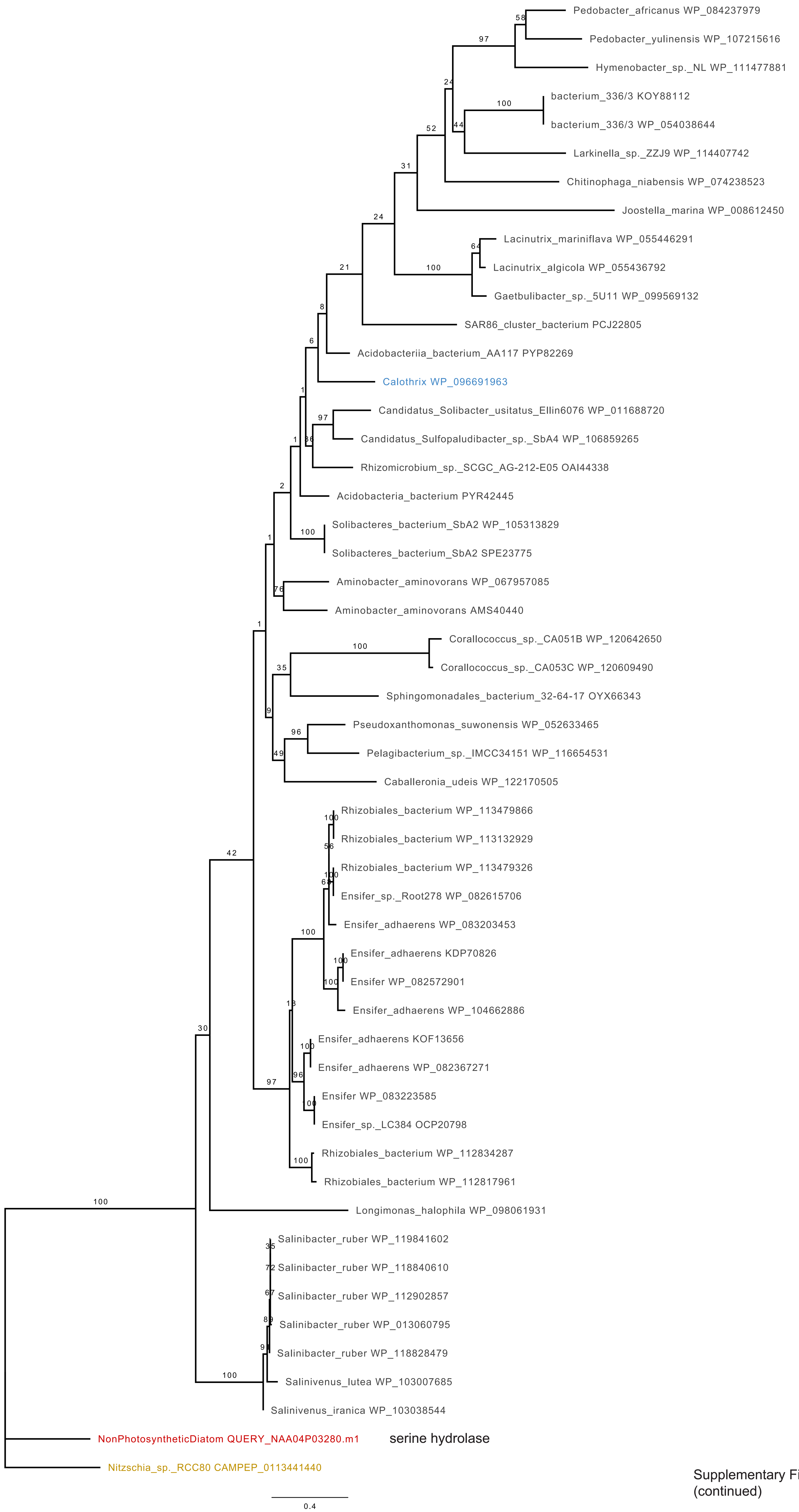

Supplementary Fig. S6
(continued)

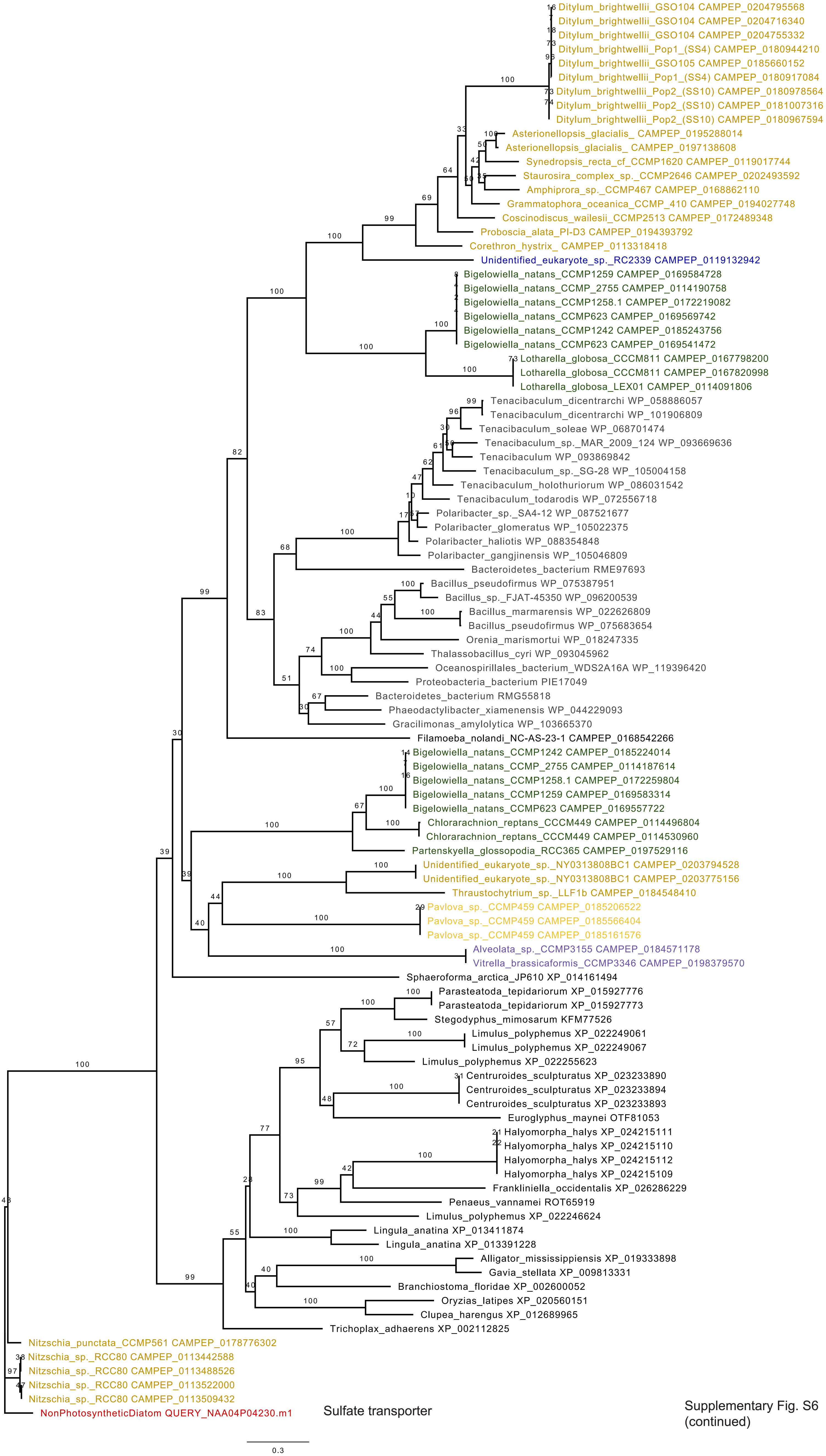

Supplementary Fig. S6 (continued)

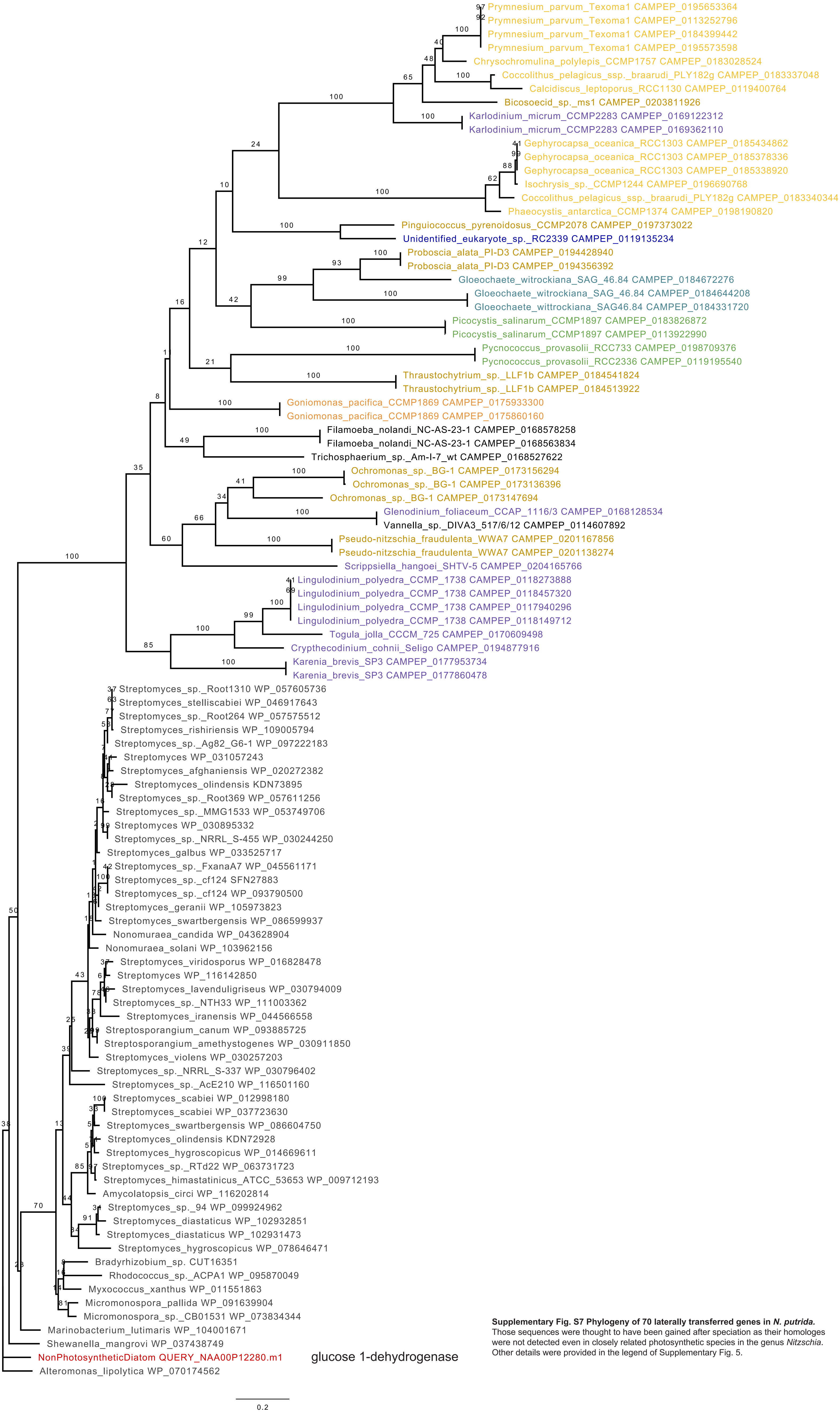

**Supplementary Fig. S7 Phylogeny of 70 laterally transferred genes in *N. putrida*.** Those sequences were thought to have been gained after speciation as their homologes were not detected even in closely related photosynthetic species in the genus *Nitzschia*. Other details were provided in the legend of Supplementary Fig. 5.

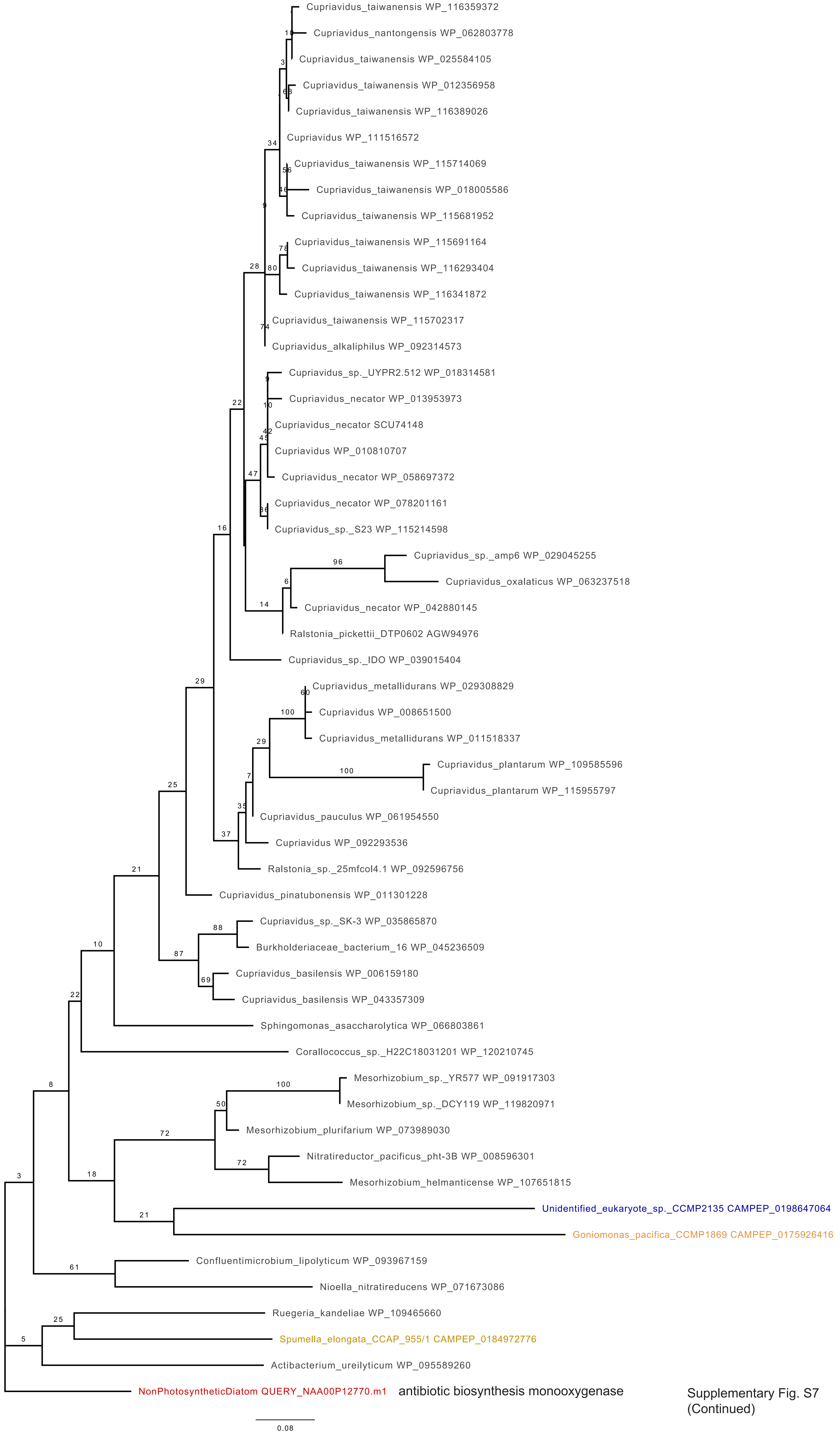

Supplementary Fig. S7
(Continued)

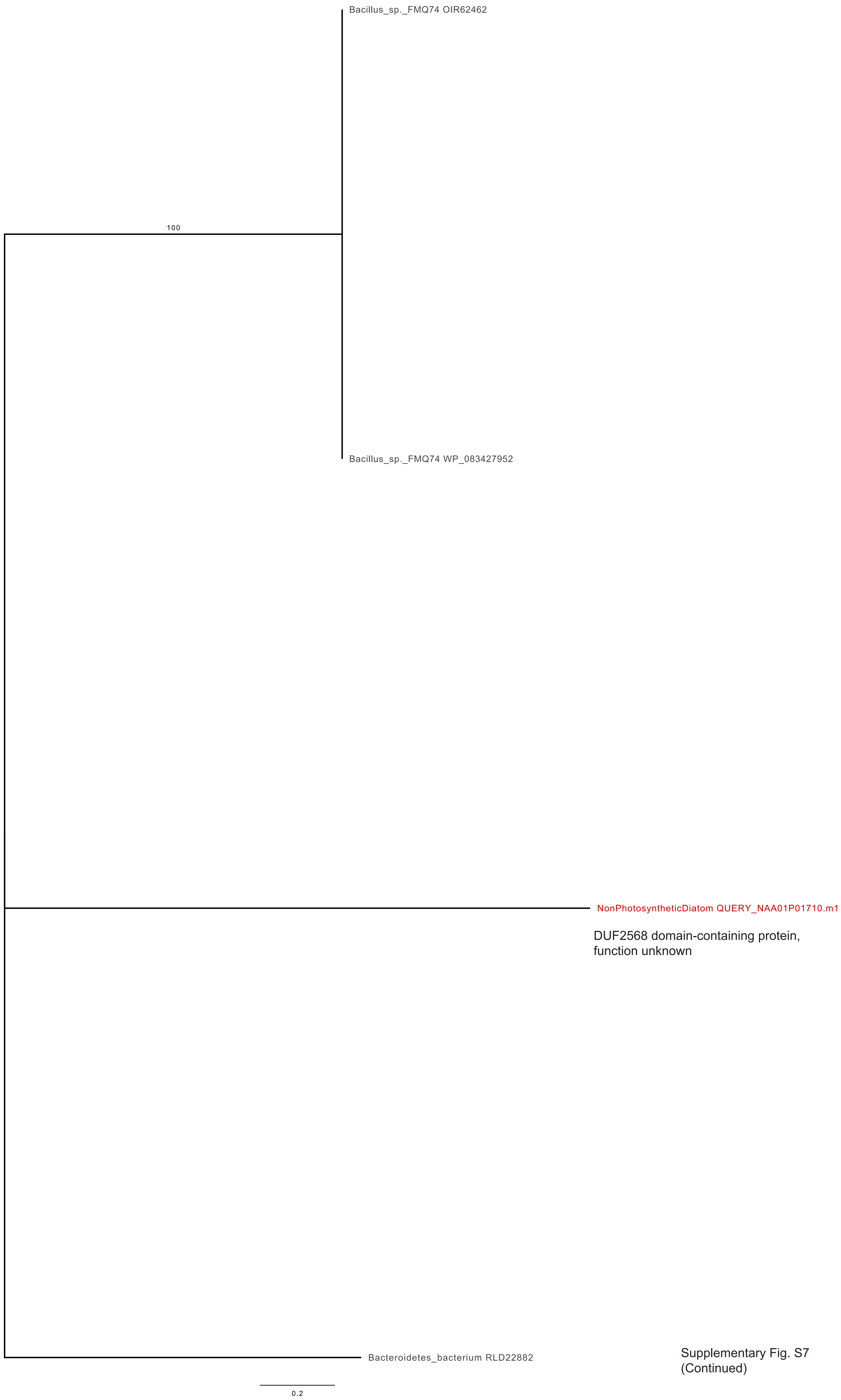

Supplementary Fig. S7
(Continued)

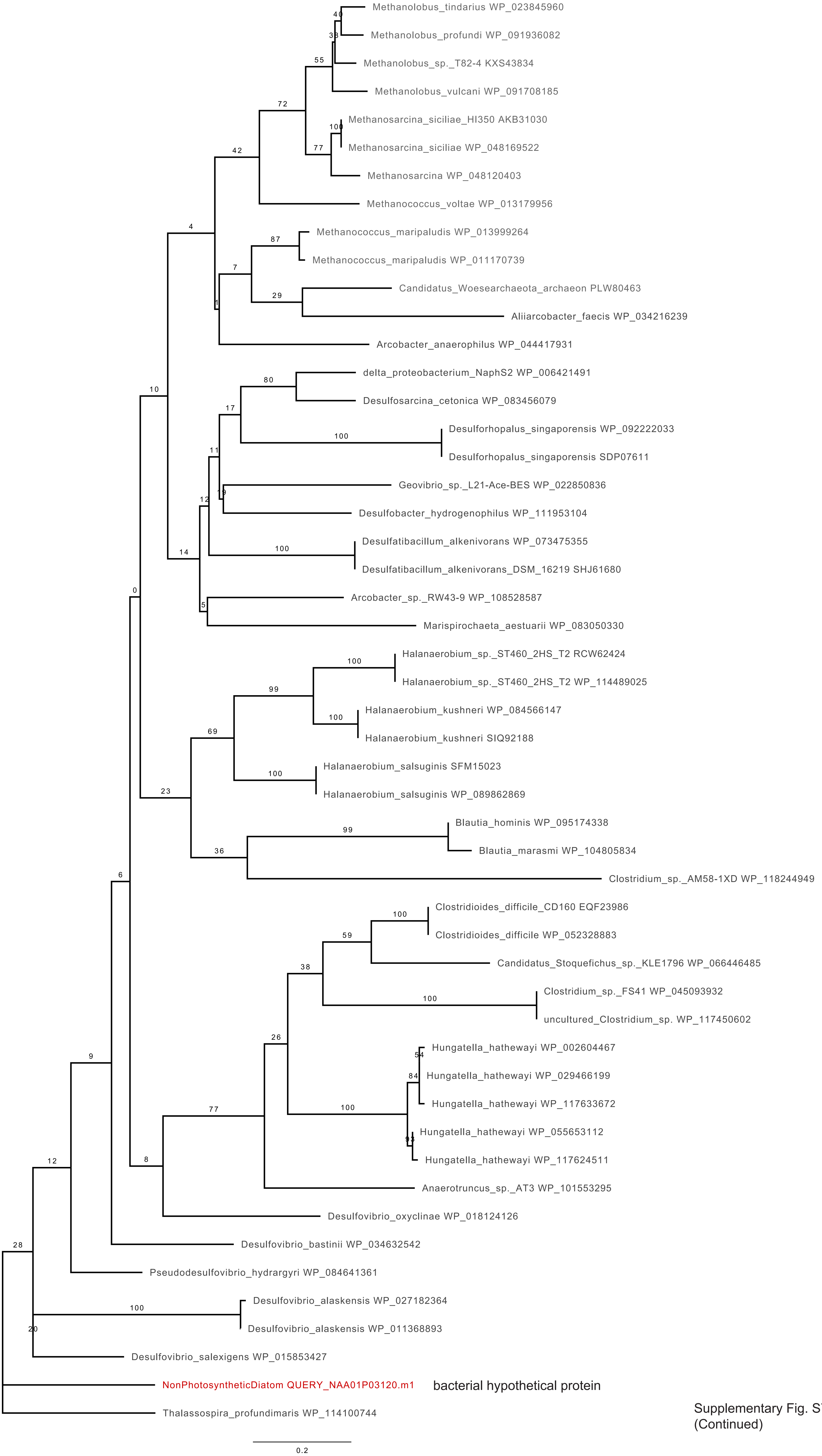

Supplementary Fig. S7
(Continued)

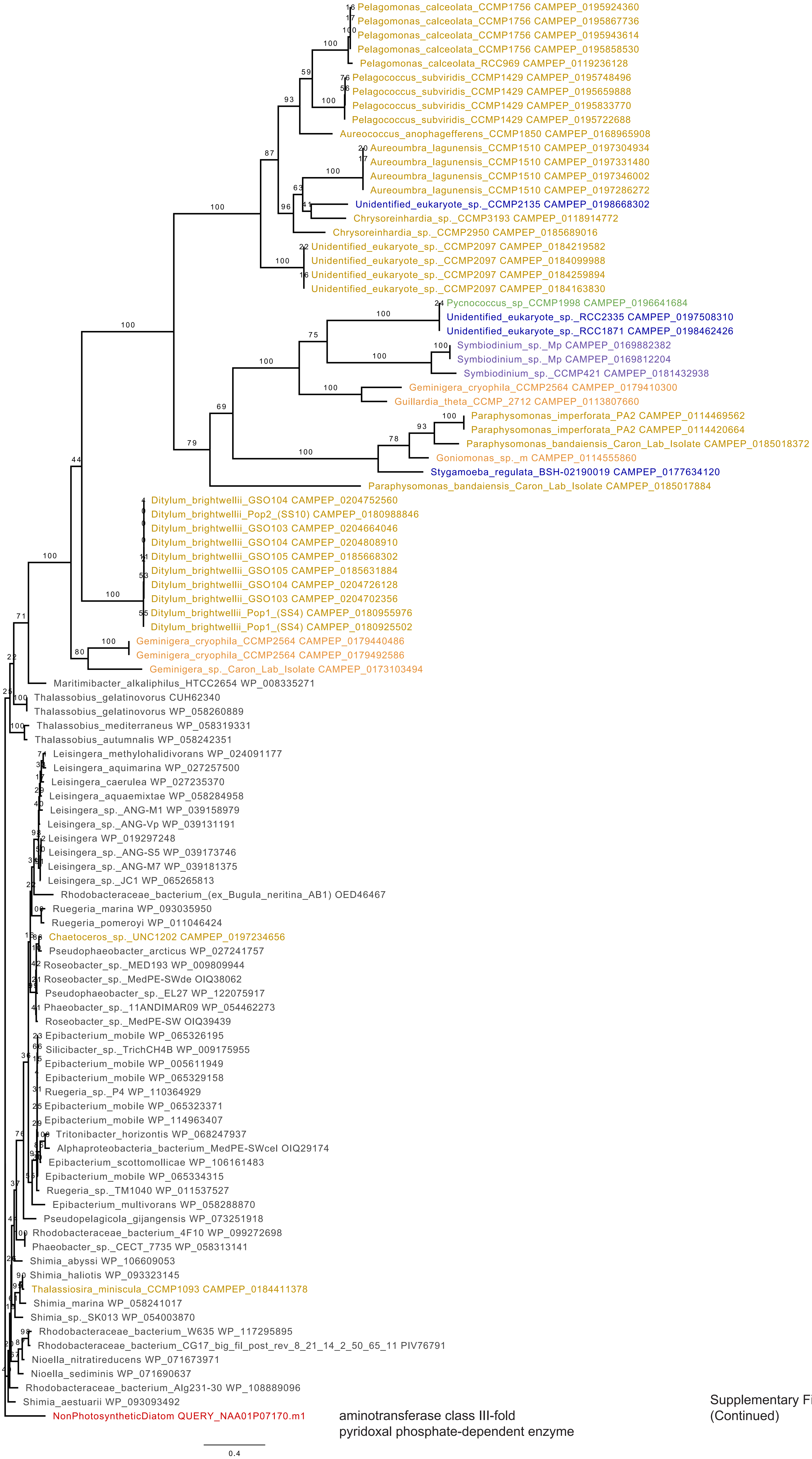

Supplementary Fig. S7
(Continued)

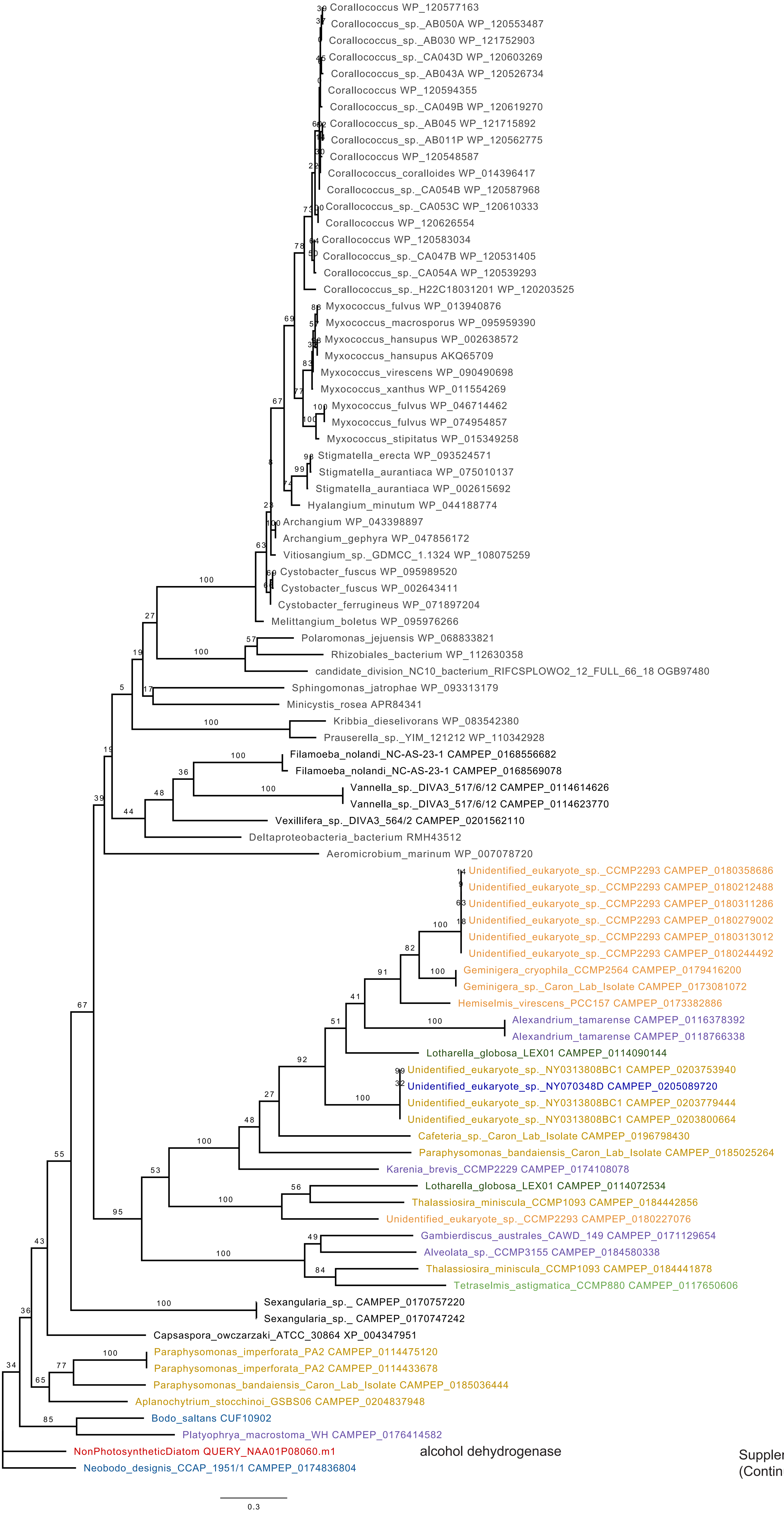

Supplementary Fig. S7
(Continued)

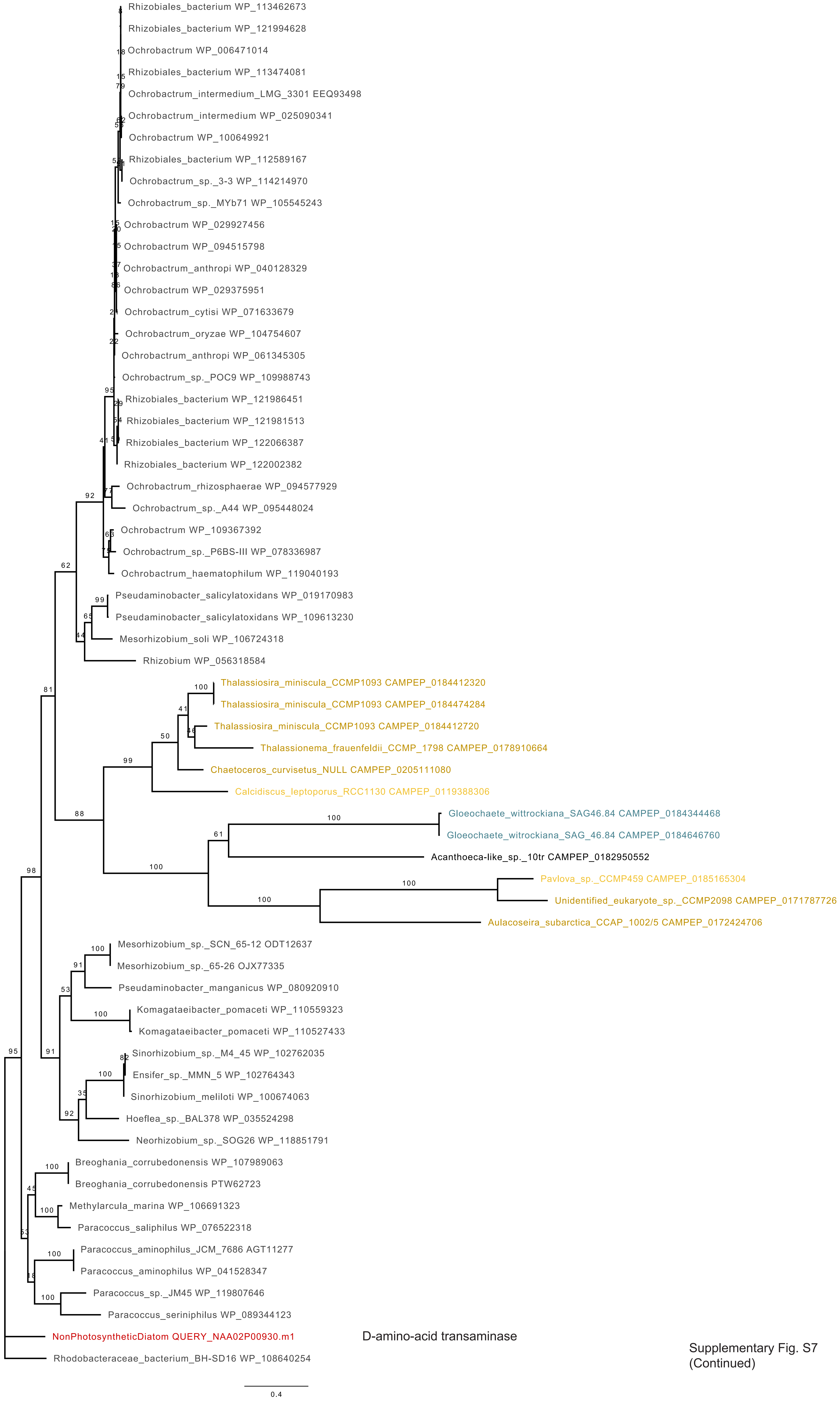

Supplementary Fig. S7
(Continued)

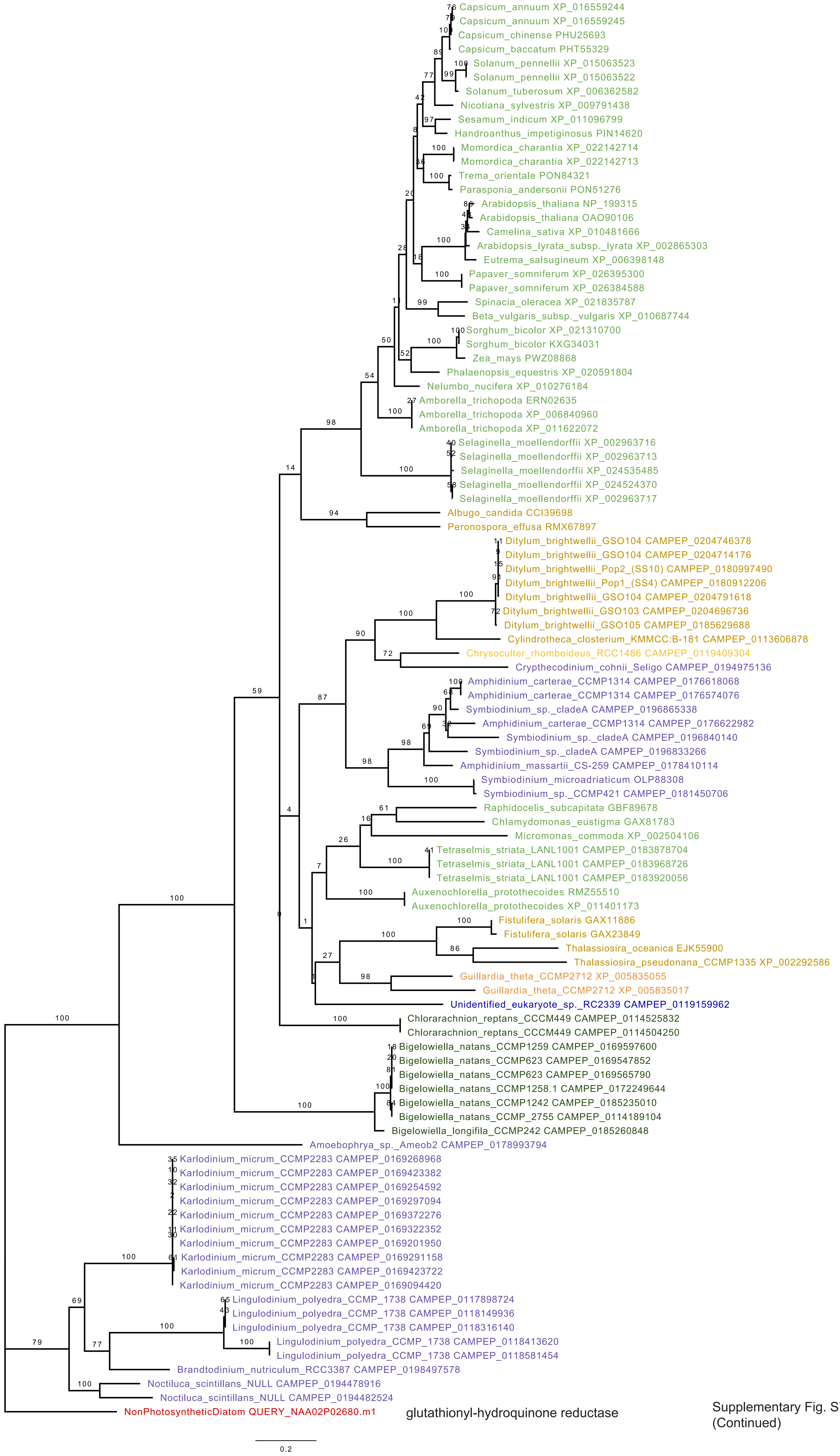

glutathionyl-hydroquinone reductase

Supplementary Fig. S7 (Continued)

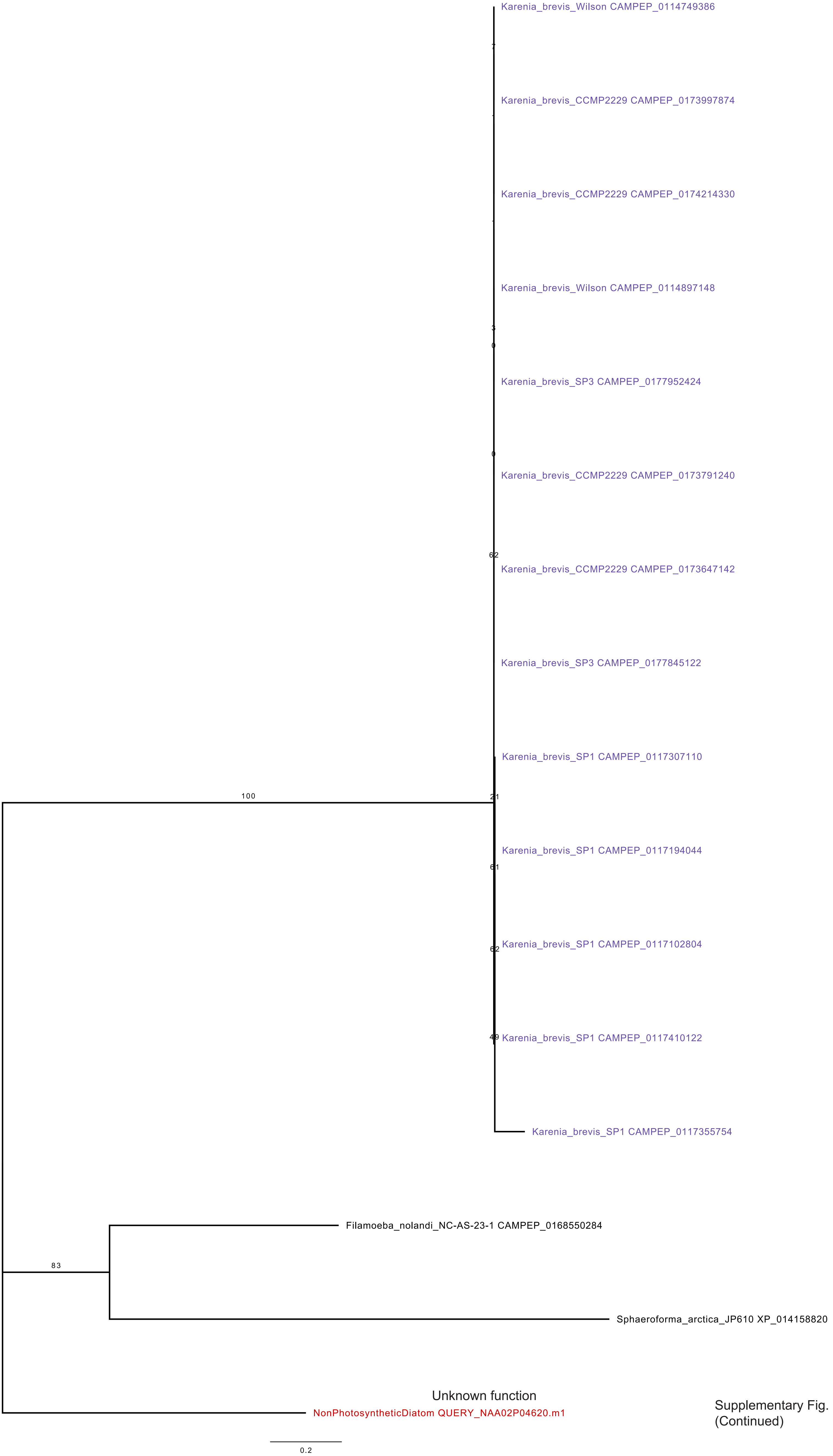

Supplementary Fig. S7
(Continued)

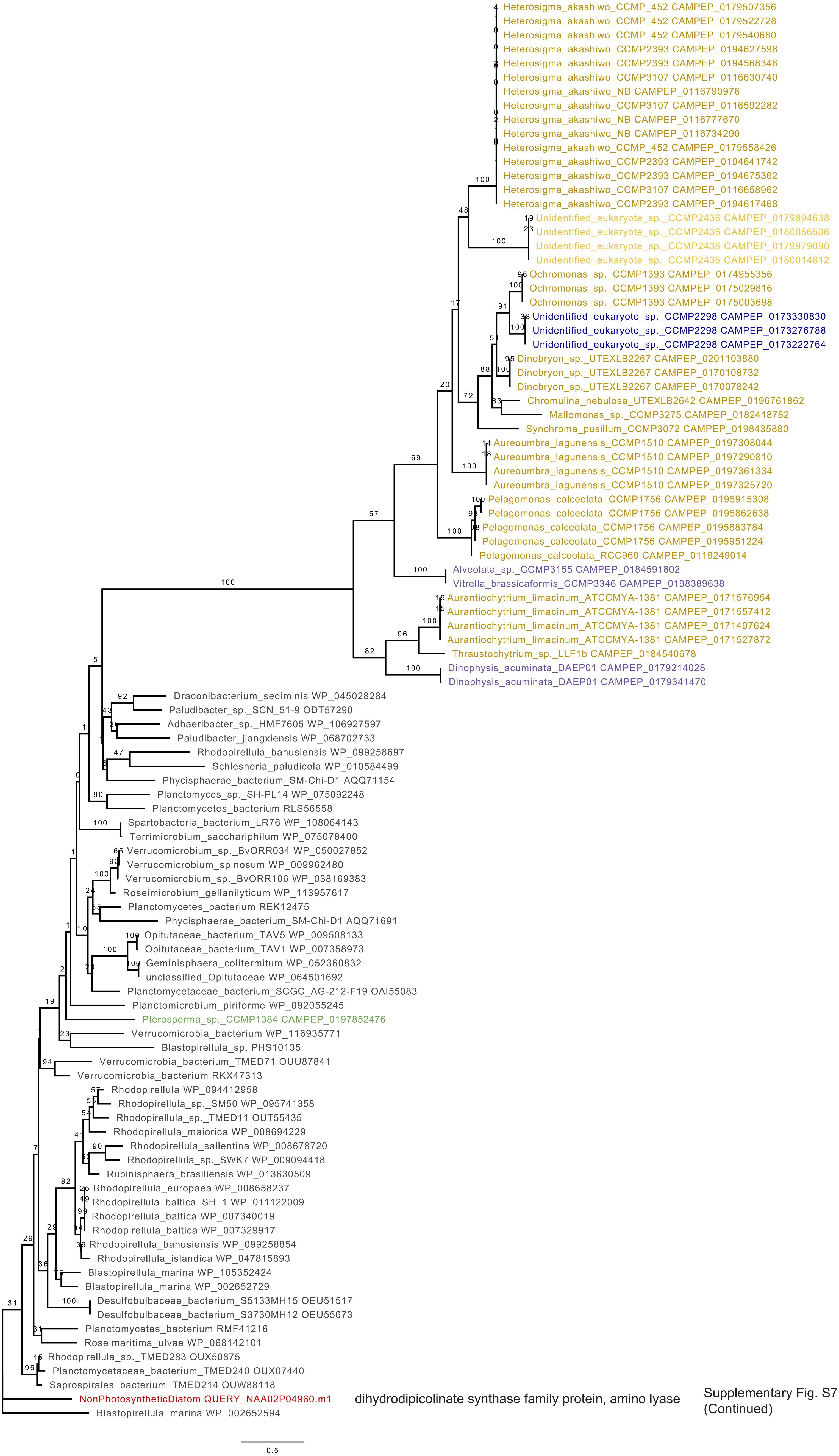

Supplementary Fig. S7 (Continued)

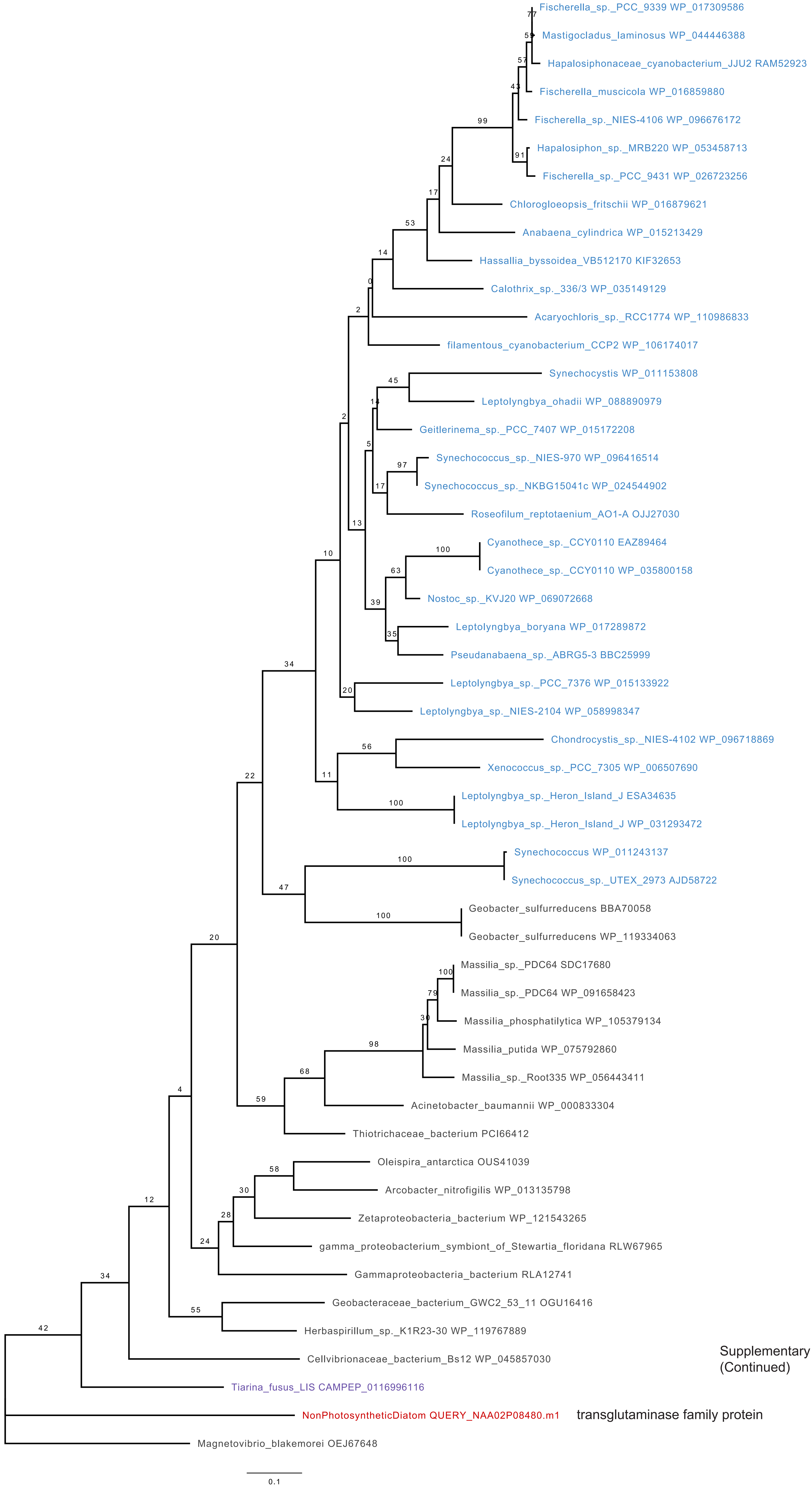

Supplementary Fig. S7 (Continued)

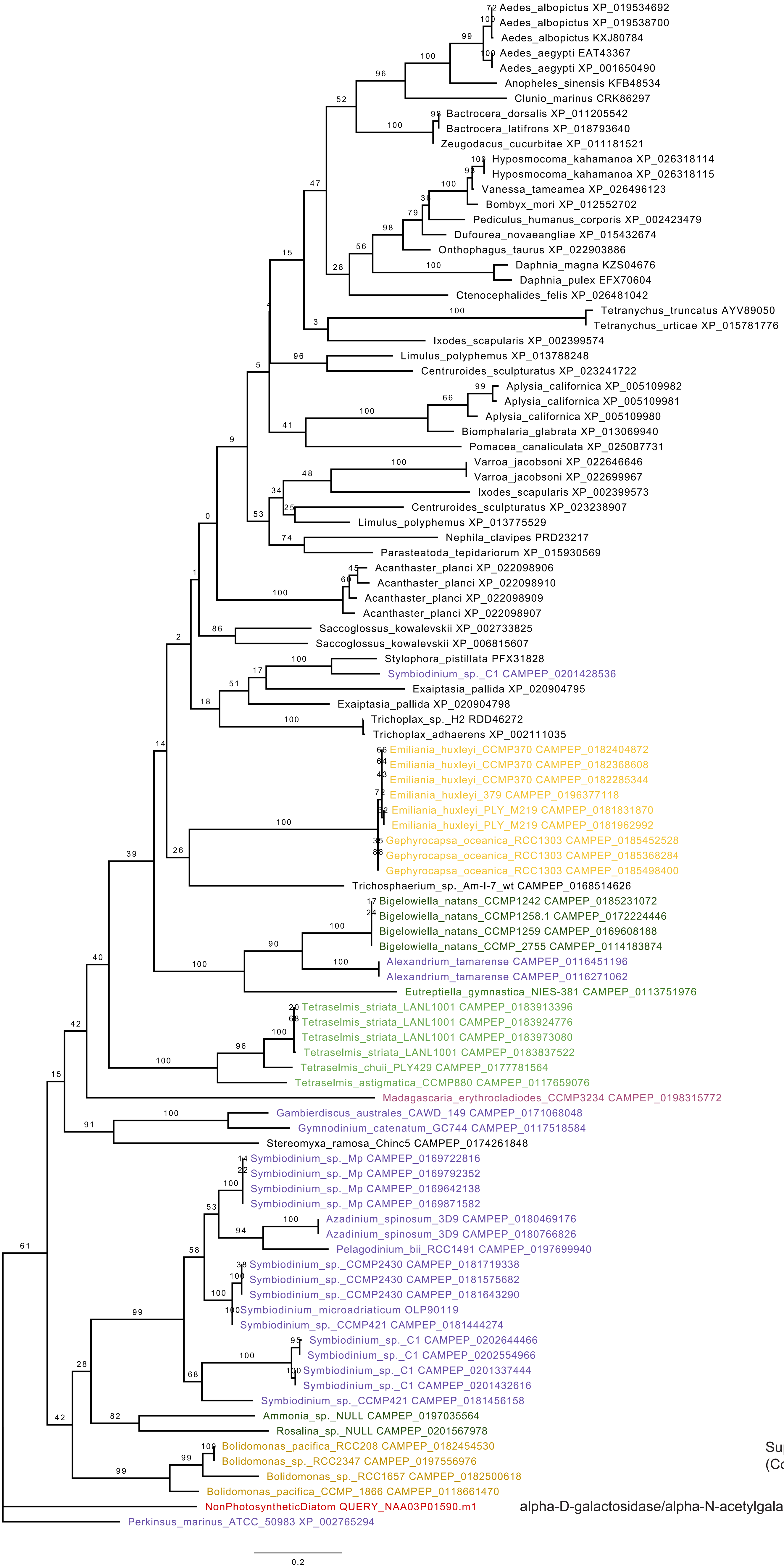

Supplementary Fig. S7 (Continued)

alpha-D-galactosidase/alpha-N-acetylgalactosaminidase GH27

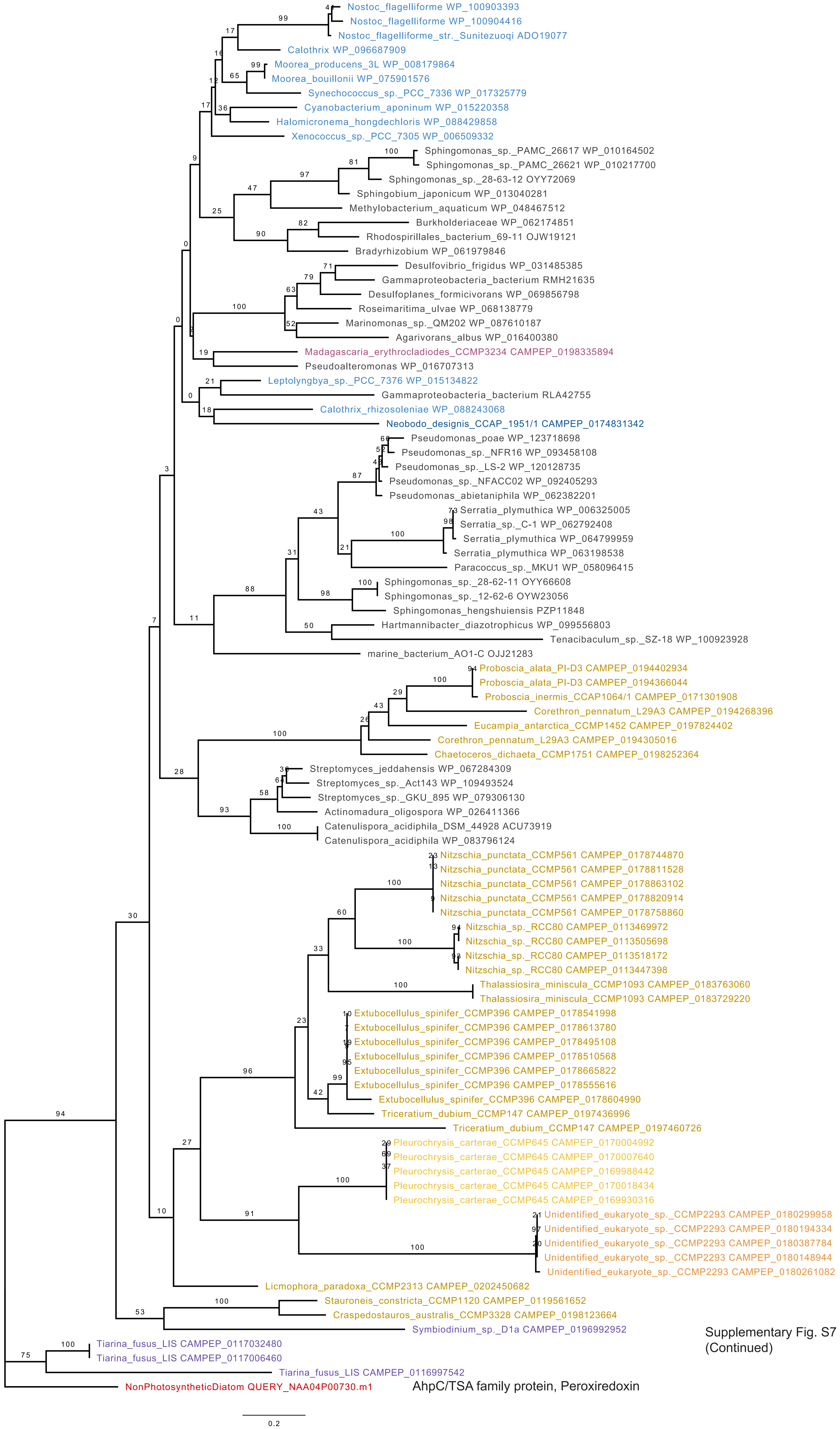

Supplementary Fig. S7
(Continued)

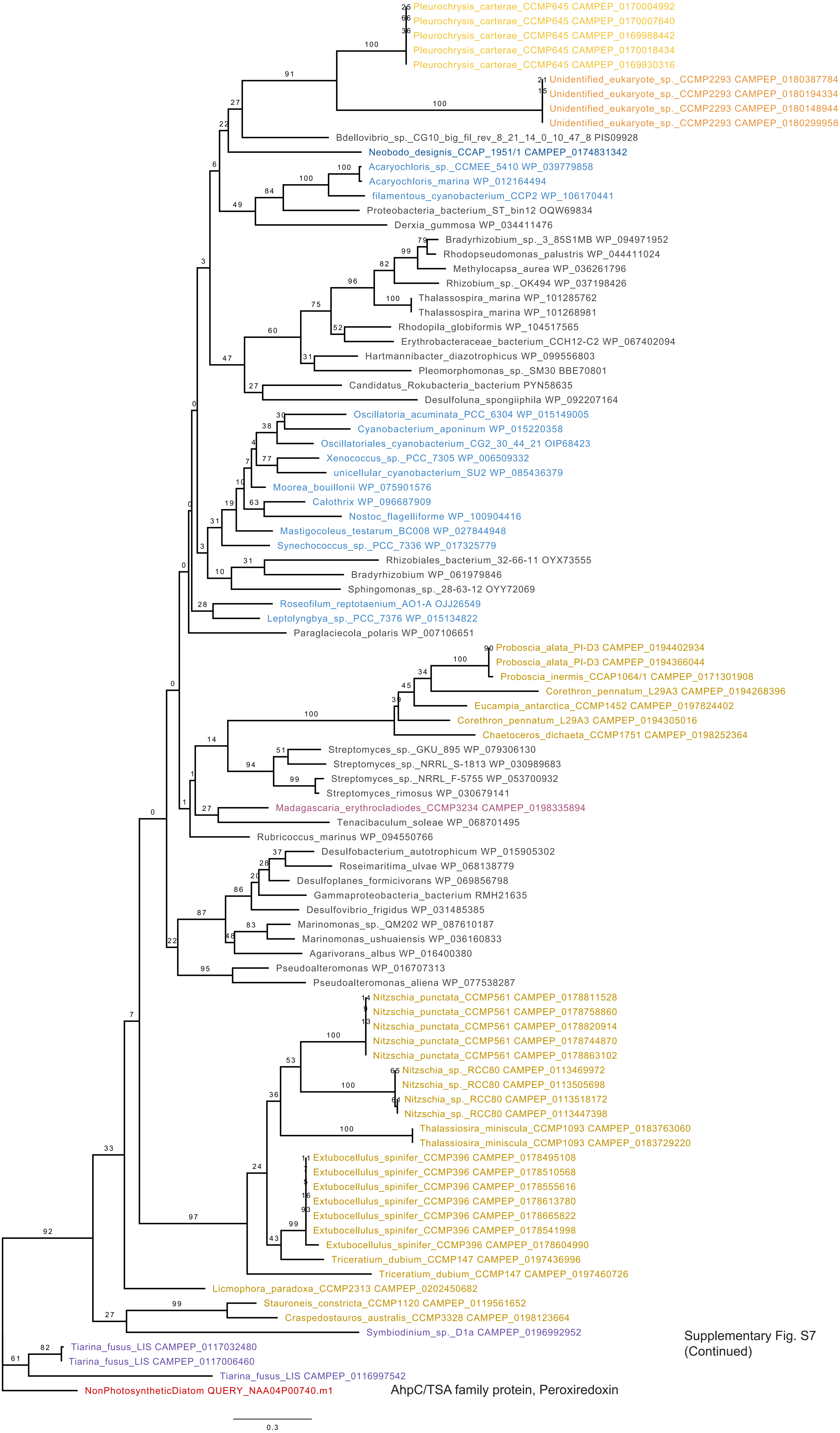

Supplementary Fig. S7 (Continued)

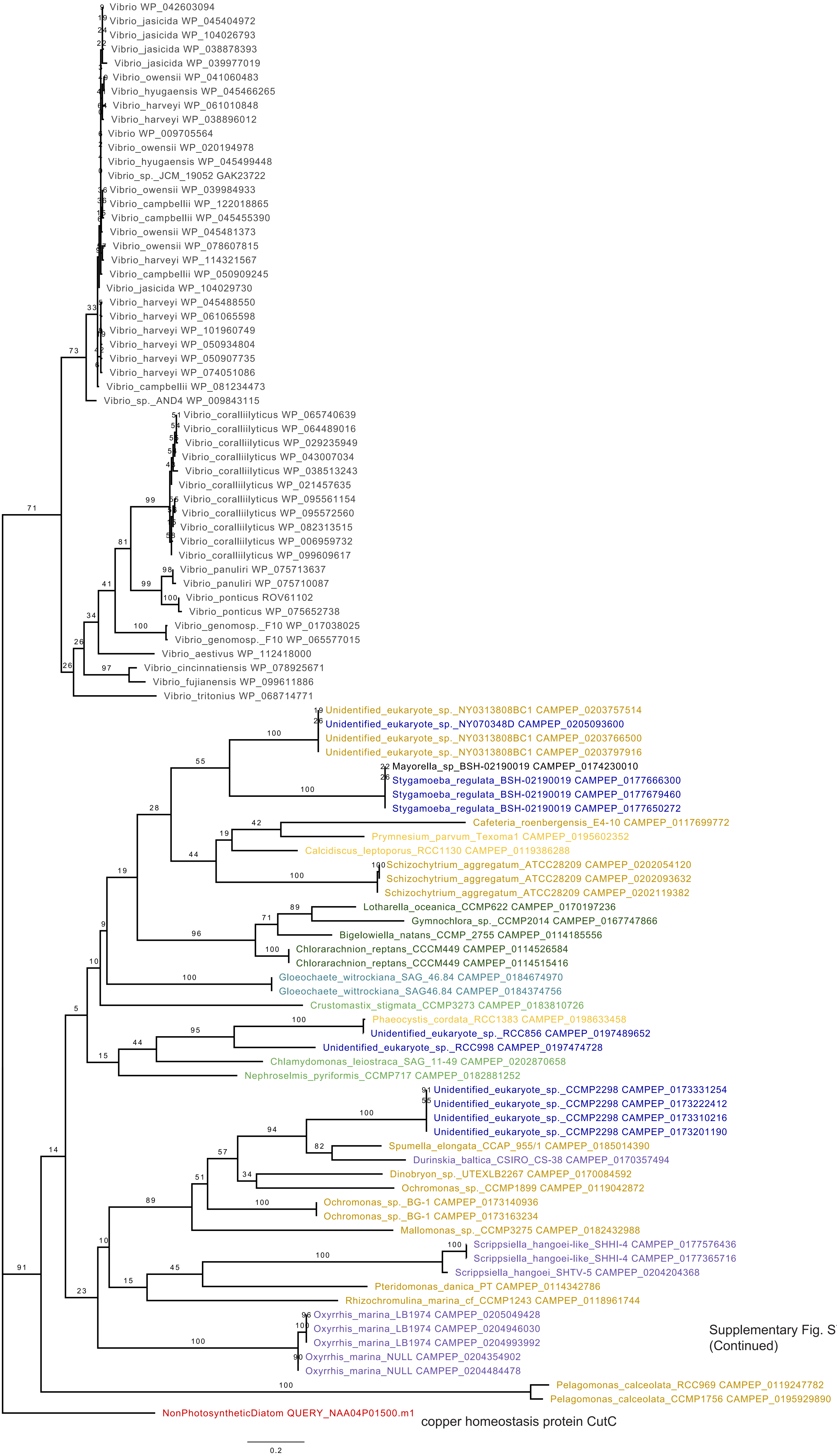

Supplementary Fig. S7
(Continued)

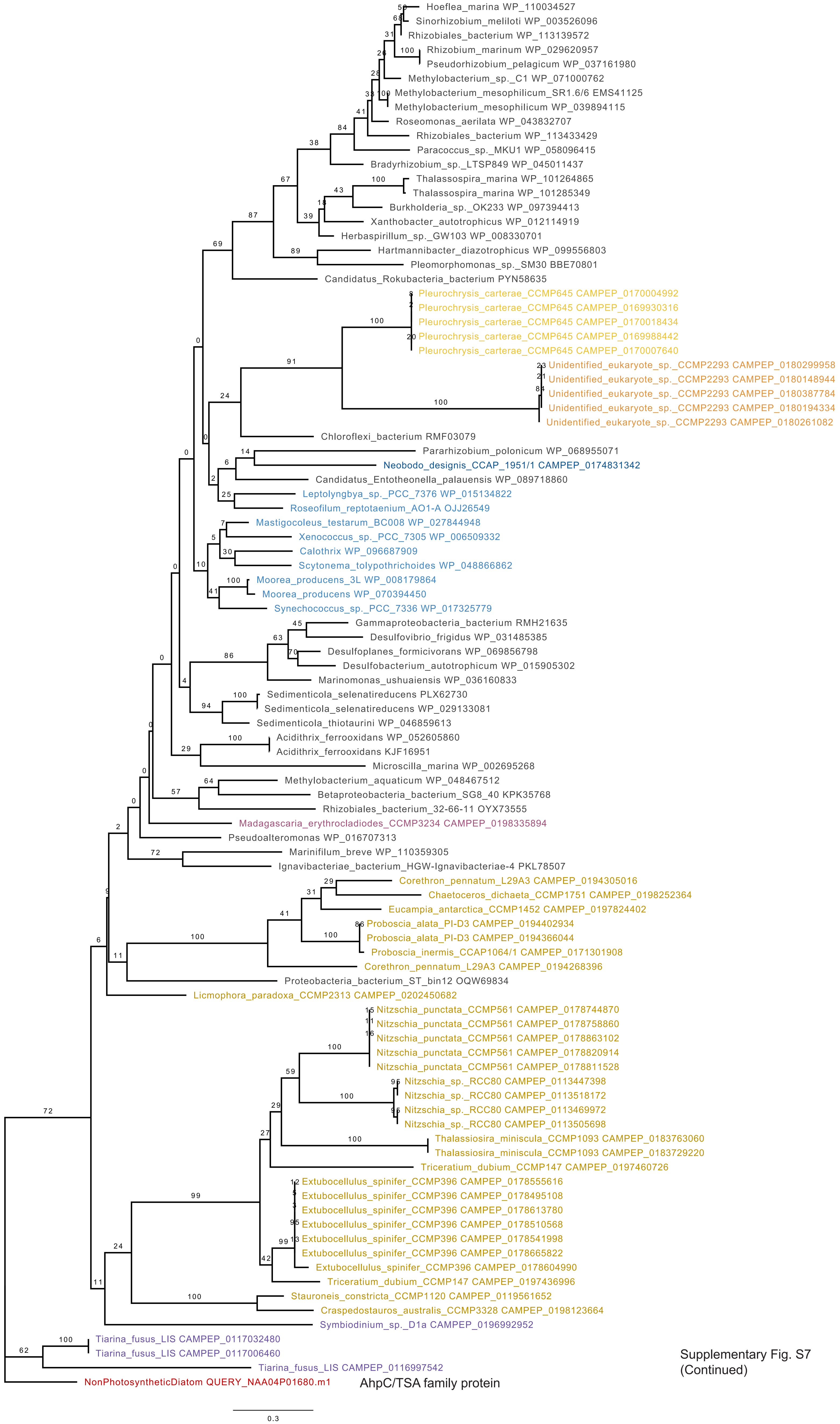

Supplementary Fig. S7
(Continued)

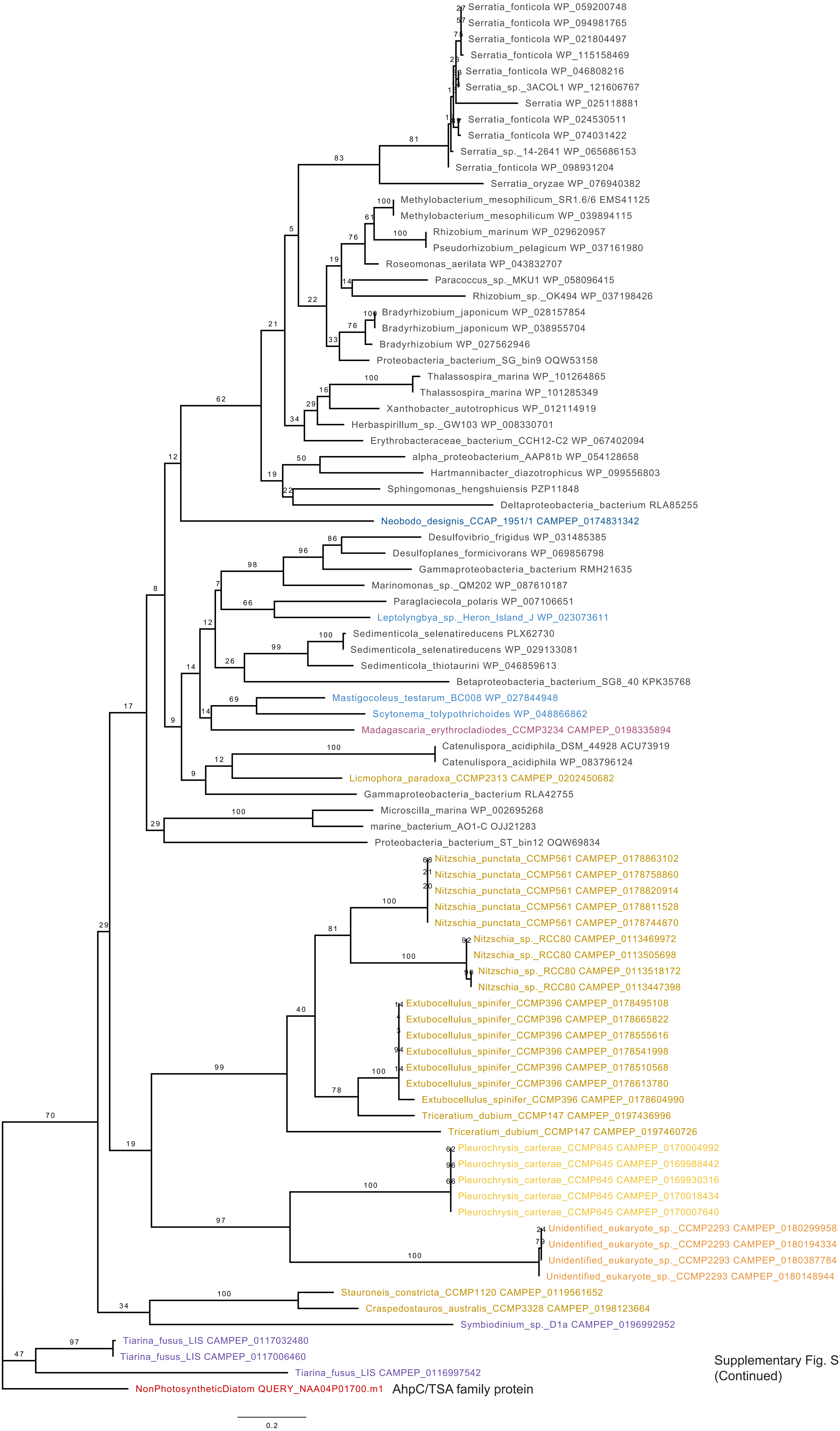

Supplementary Fig. S7
(Continued)

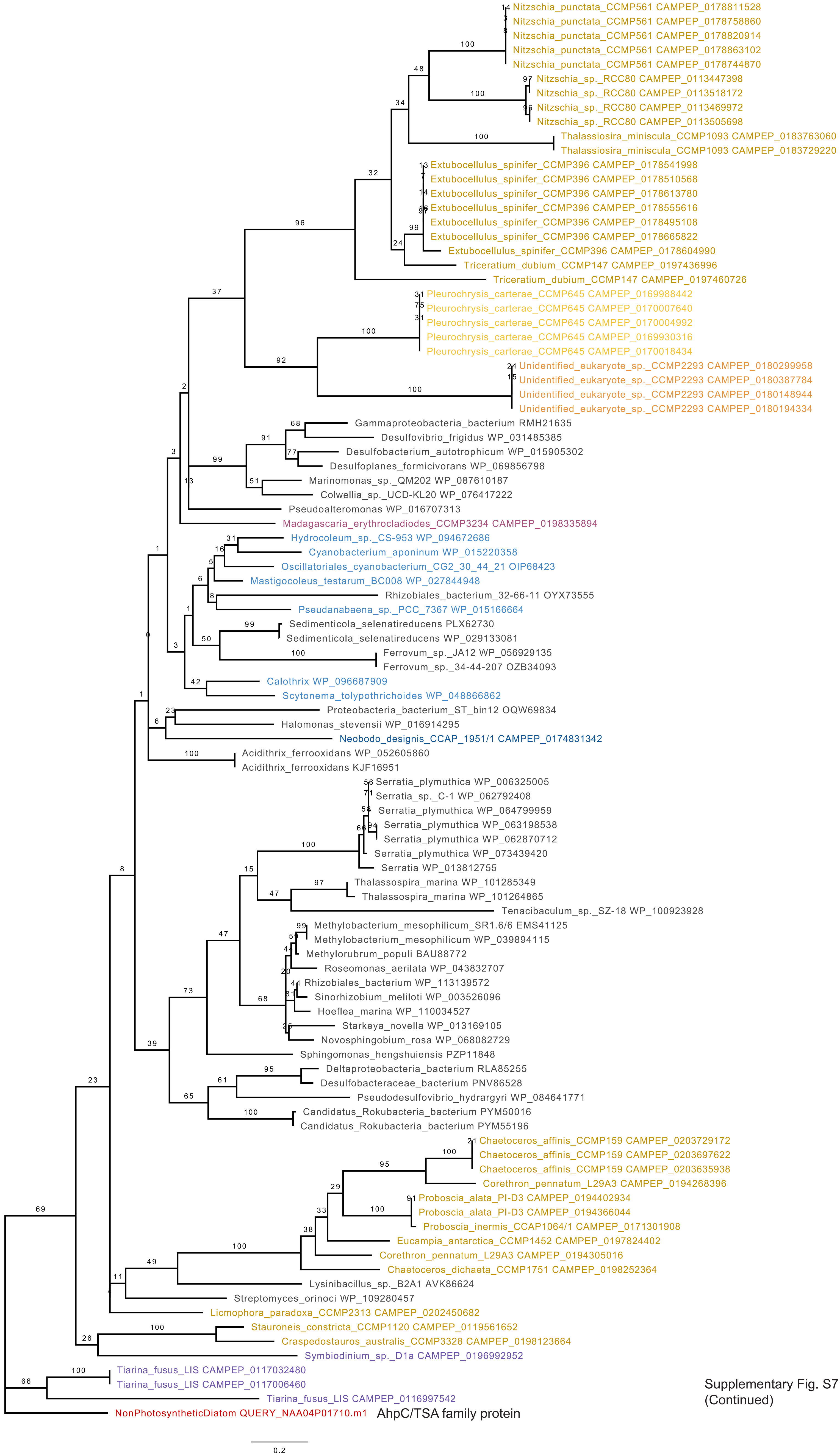

Supplementary Fig. S7
(Continued)

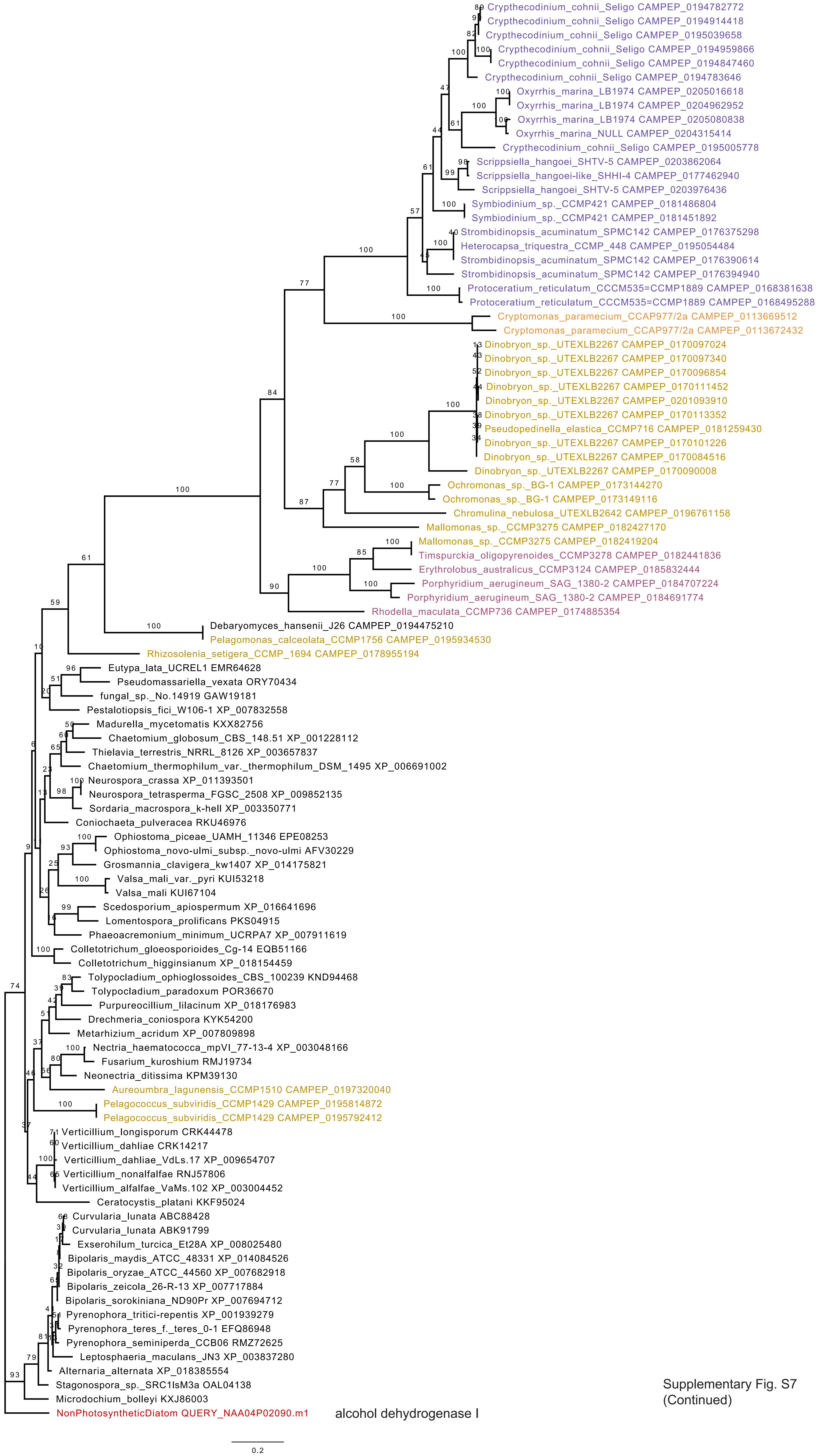

Supplementary Fig. S7
(Continued)

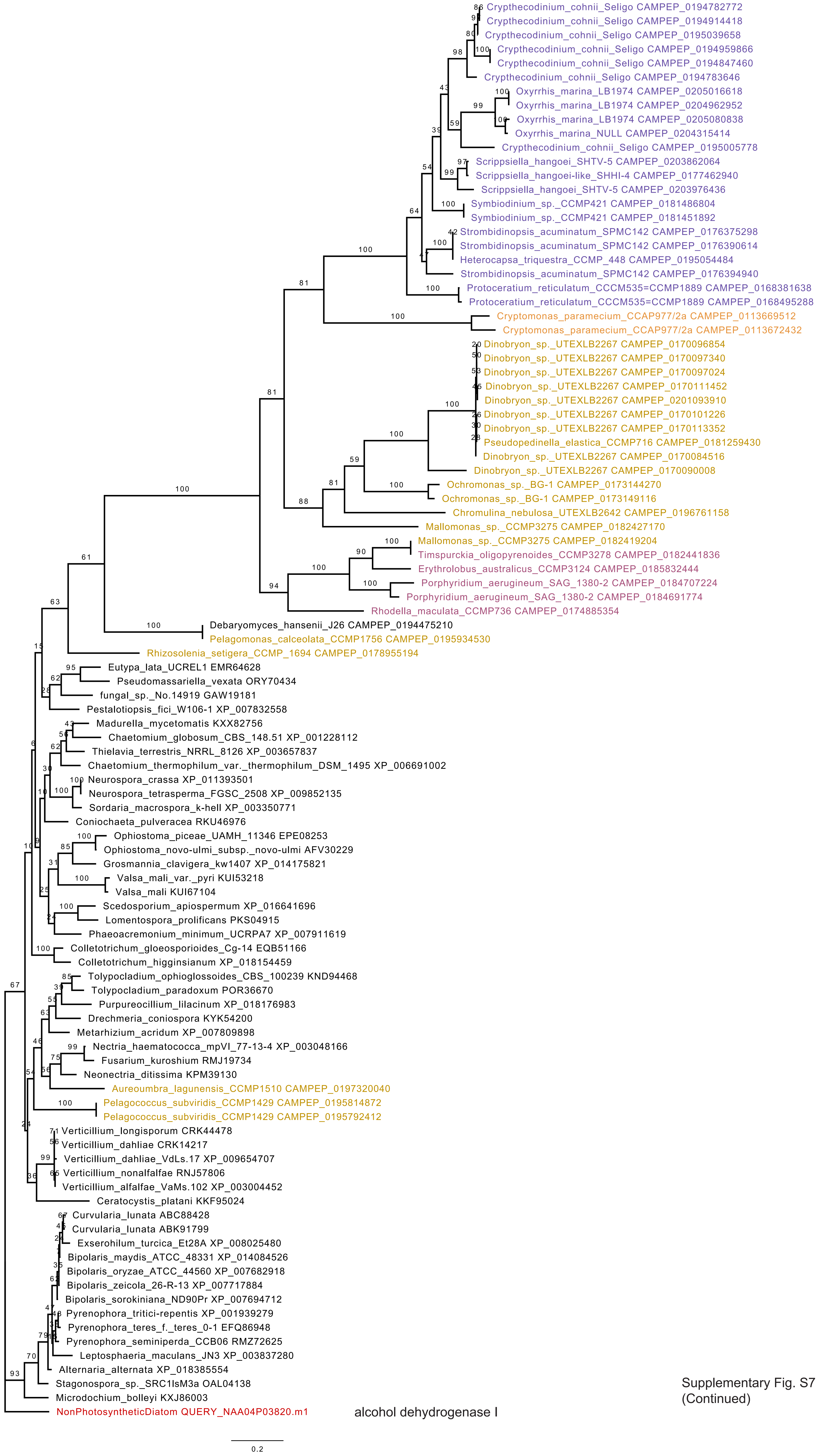

Supplementary Fig. S7
(Continued)

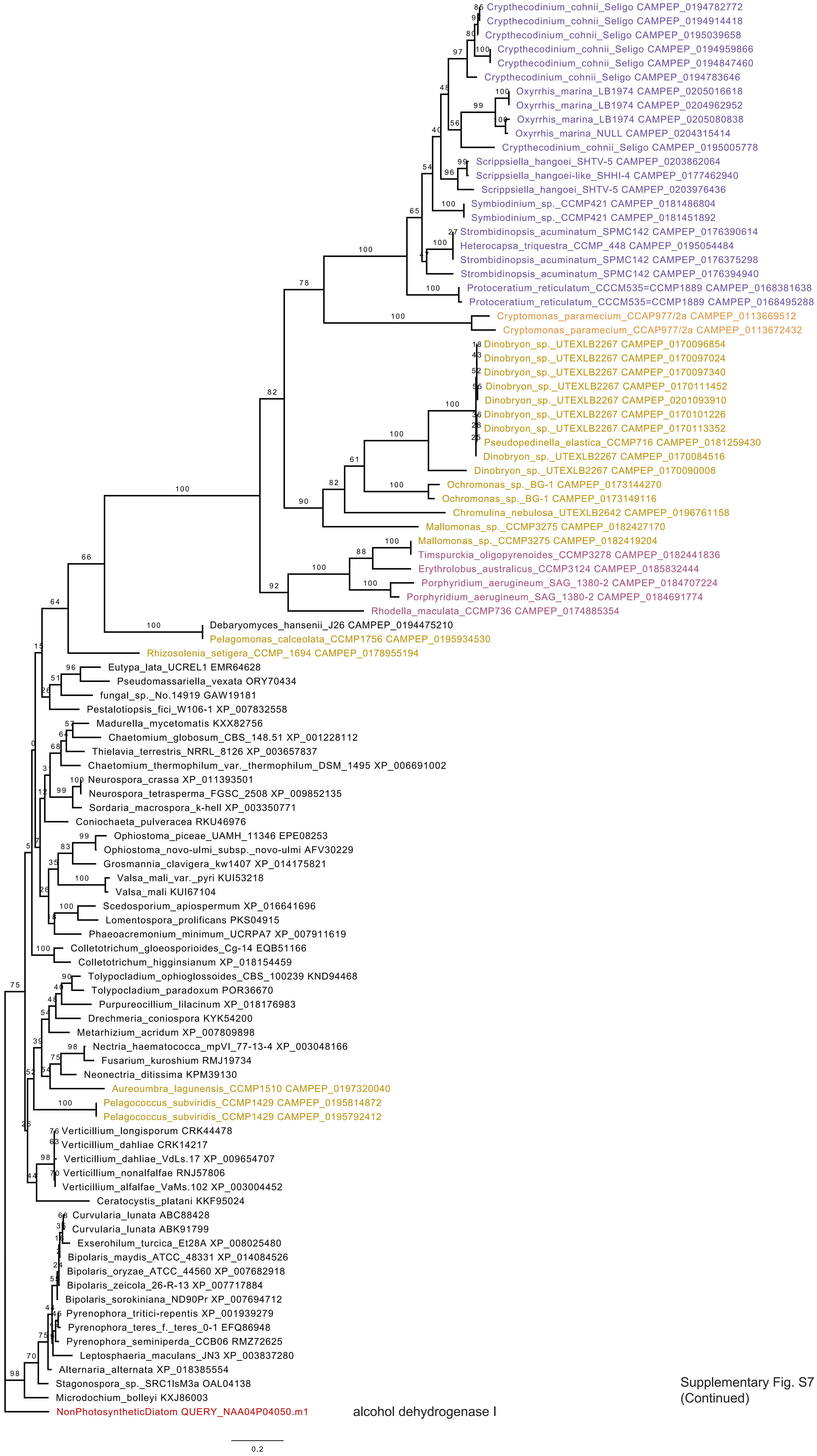

alcohol dehydrogenase I

Supplementary Fig. S7
(Continued)

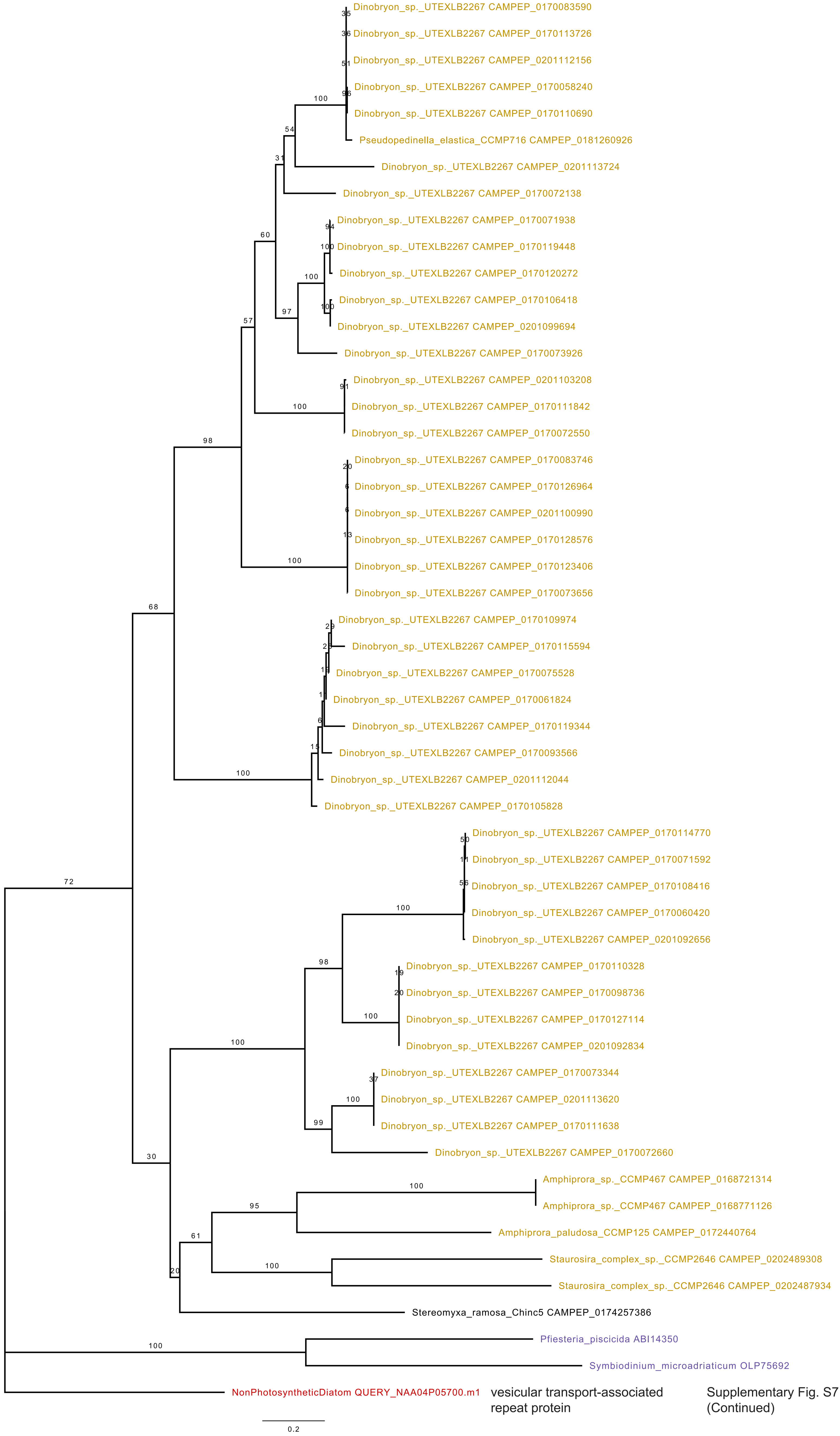

vesicular transport-associated repeat protein

Supplementary Fig. S7 (Continued)

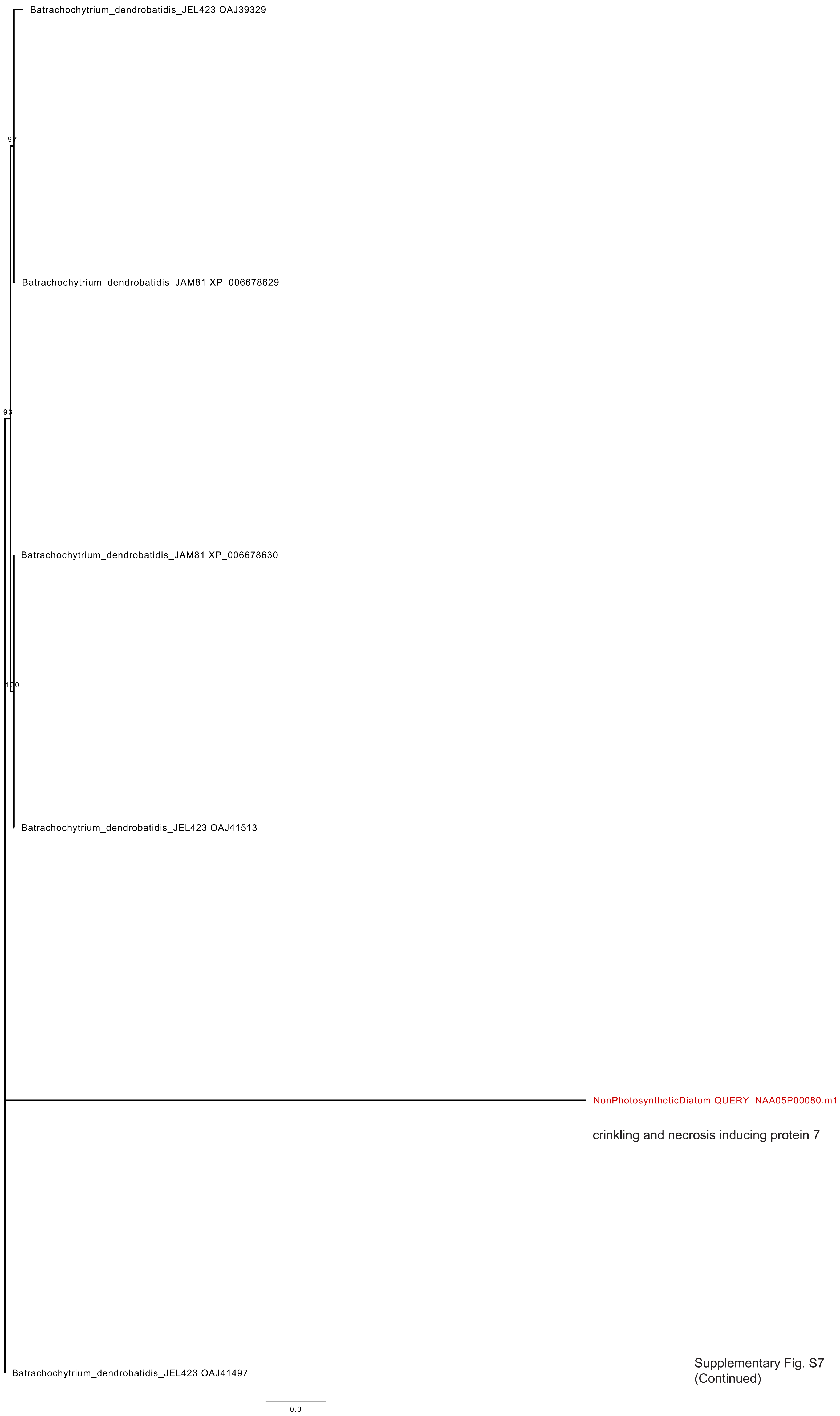

Supplementary Fig. S7
(Continued)

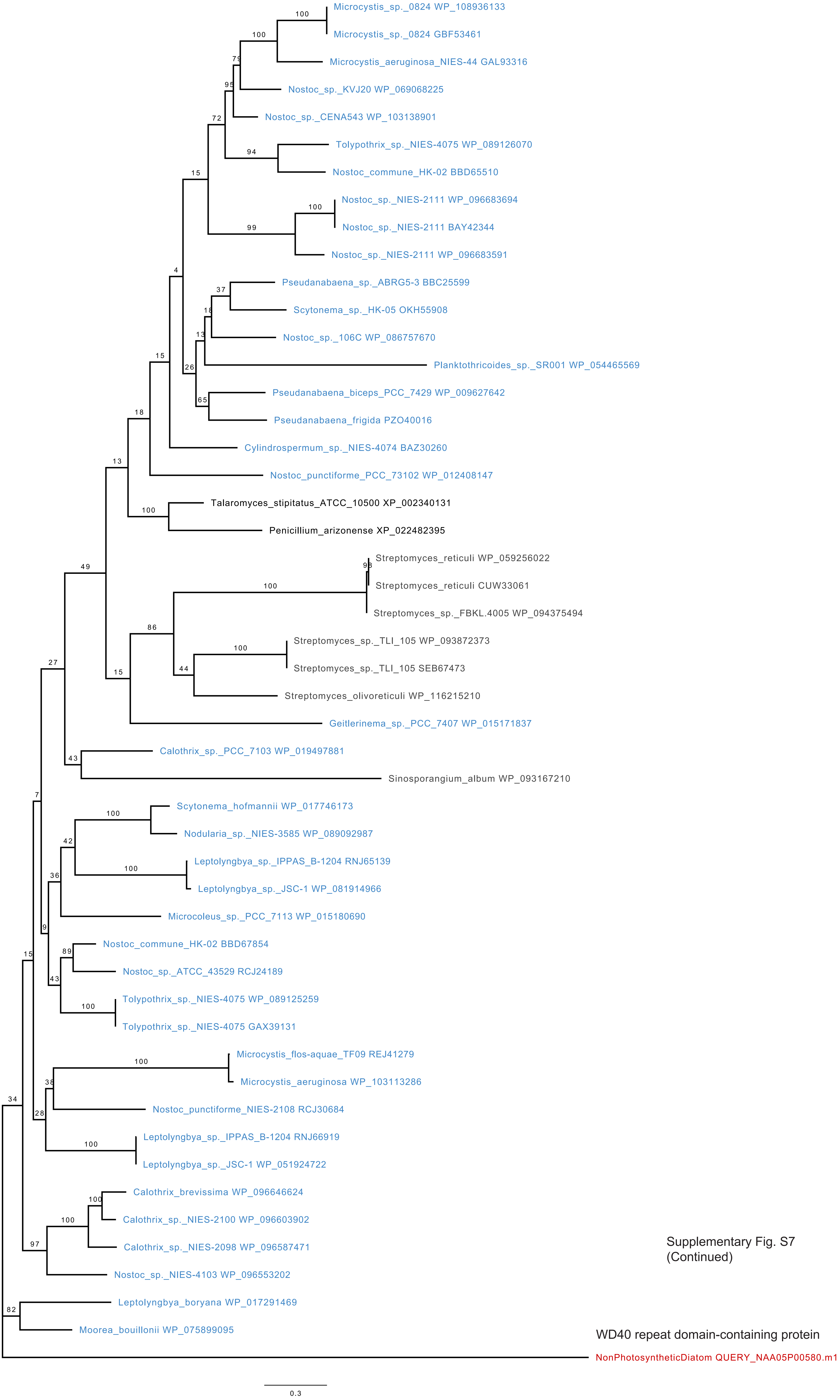

Supplementary Fig. S7 (Continued)

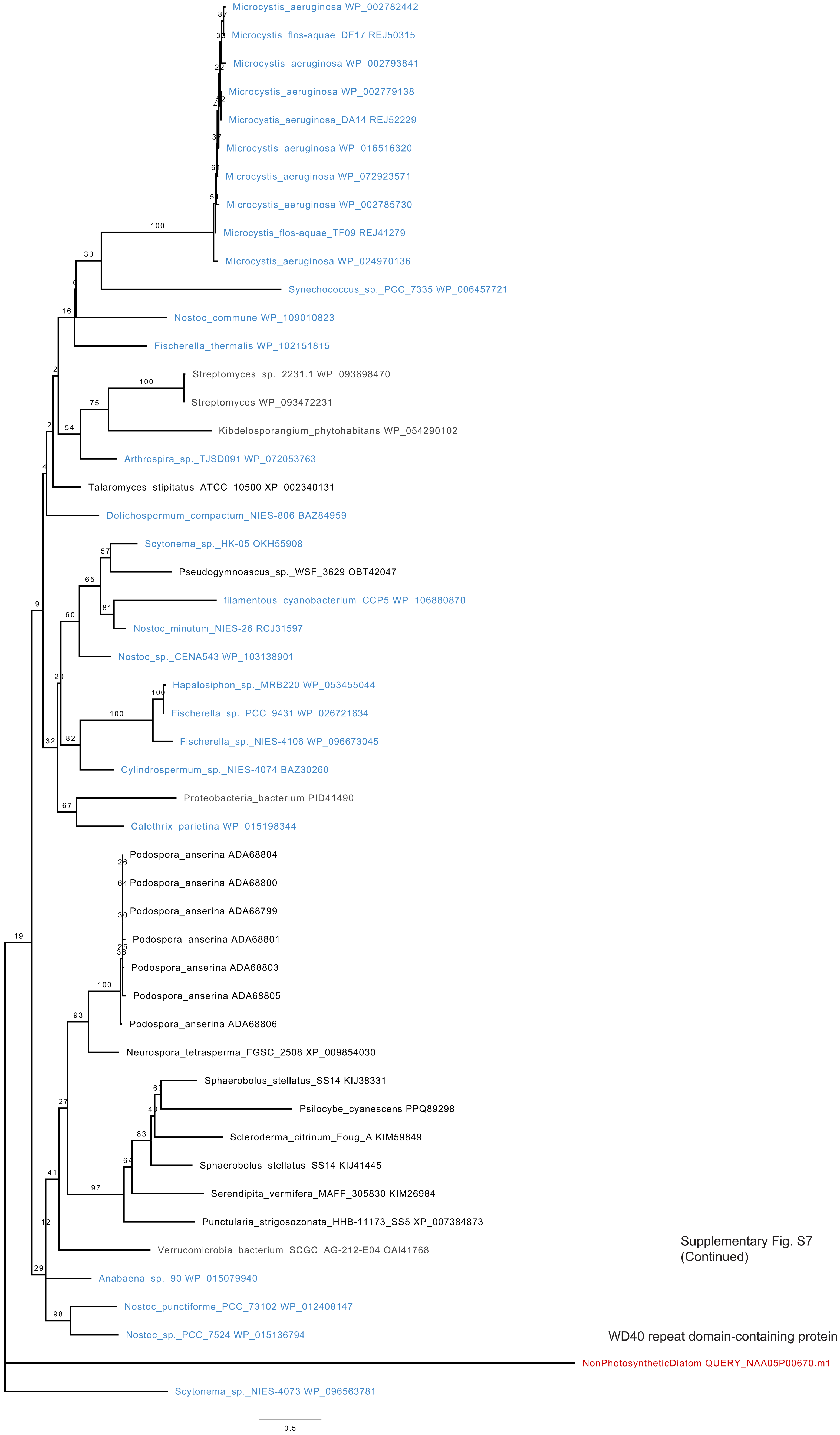

Supplementary Fig. S7
(Continued)

WD40 repeat domain-containing protein

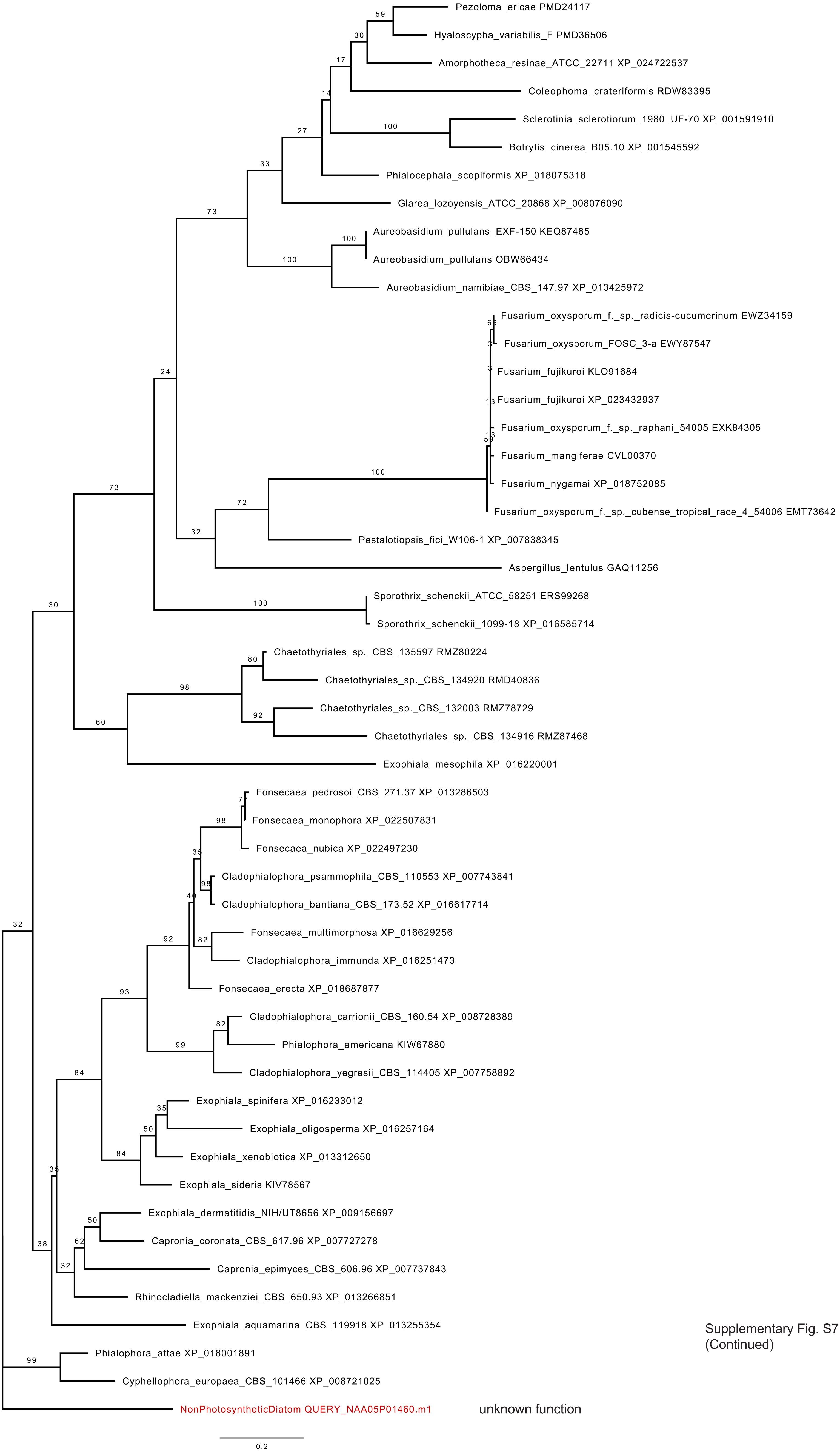

Supplementary Fig. S7
(Continued)

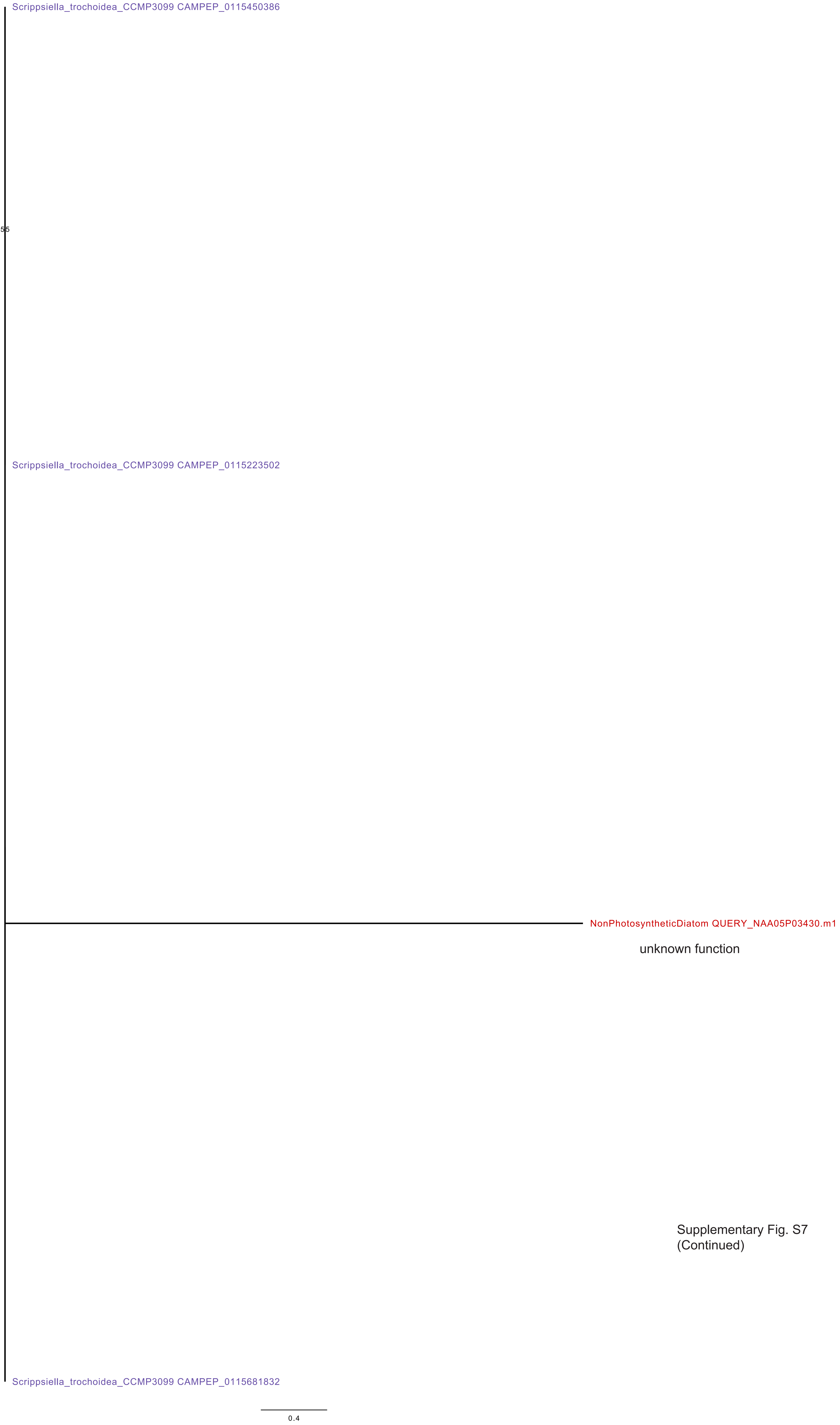

Supplementary Fig. S7
(Continued)

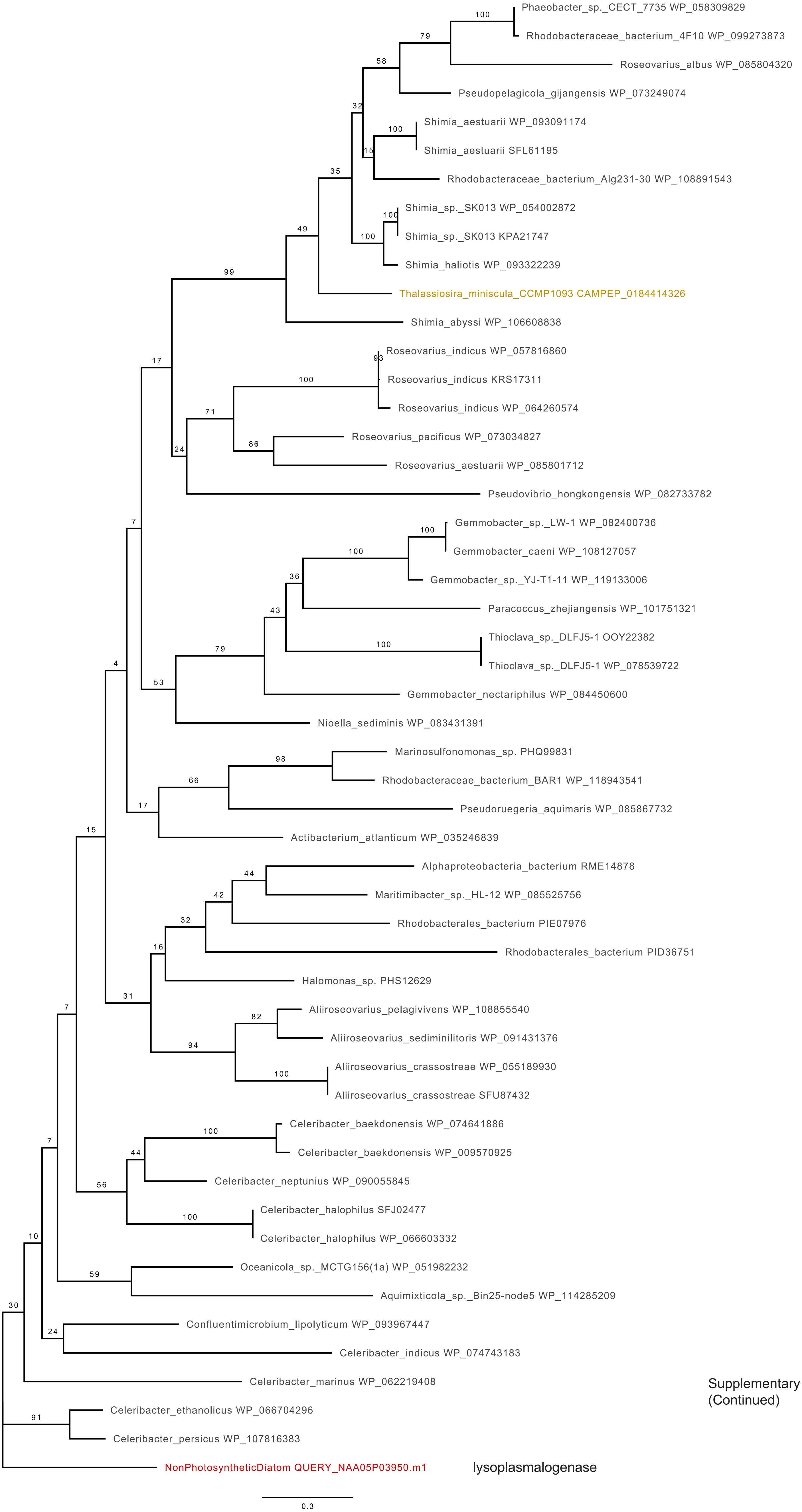

Supplementary Fig. S7
(Continued)

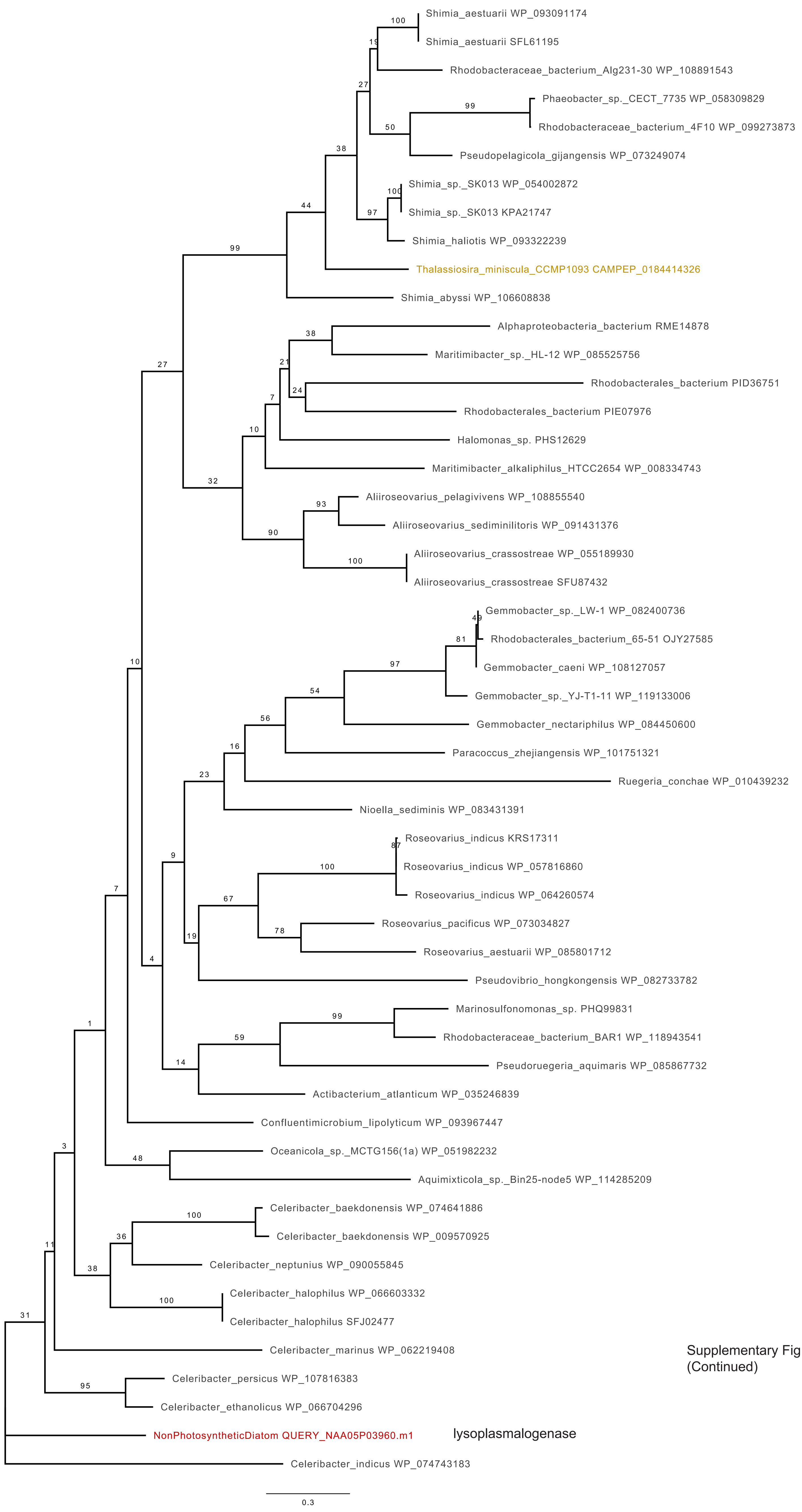

Supplementary Fig. S7
(Continued)

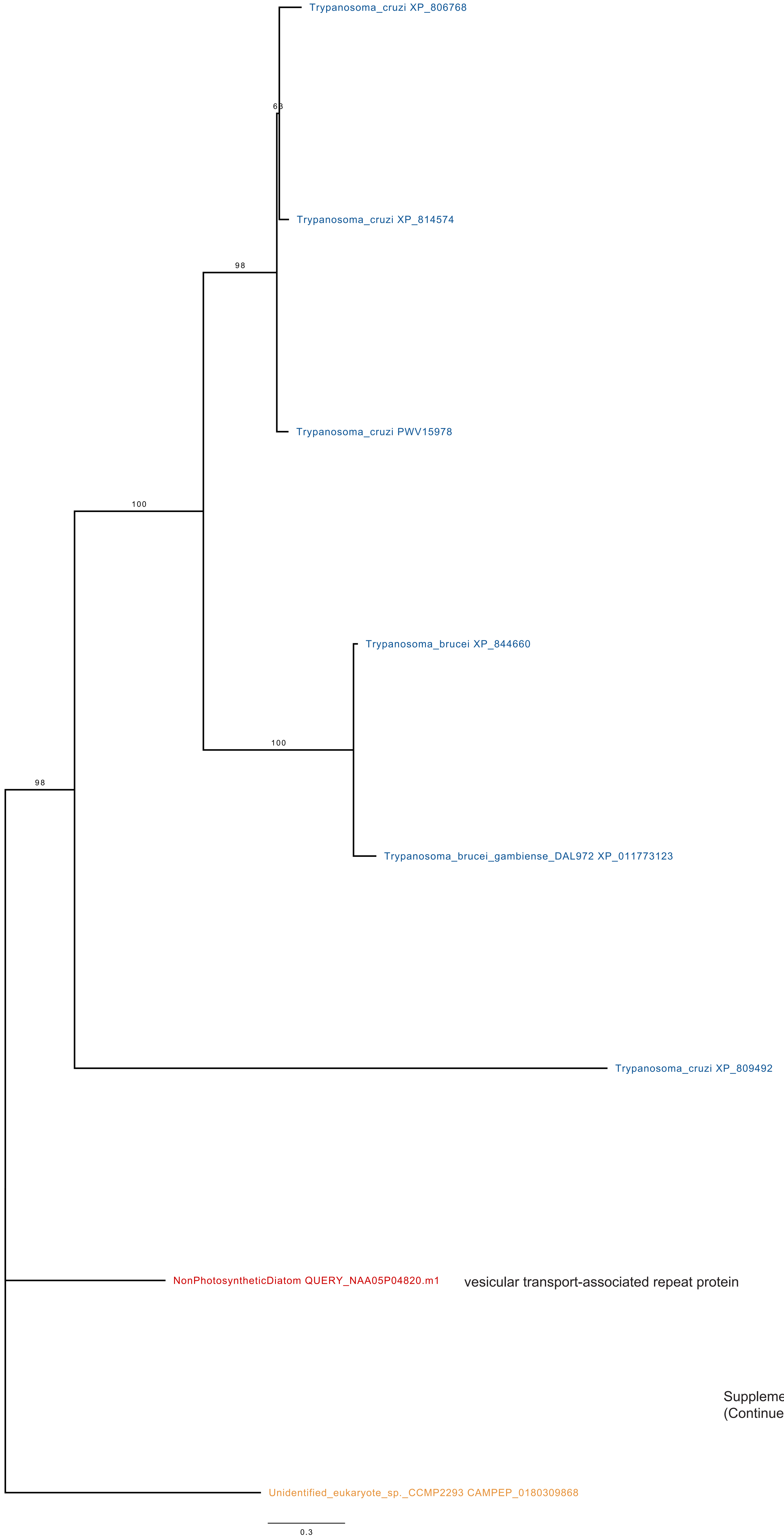

Supplementary Fig. S7
(Continued)

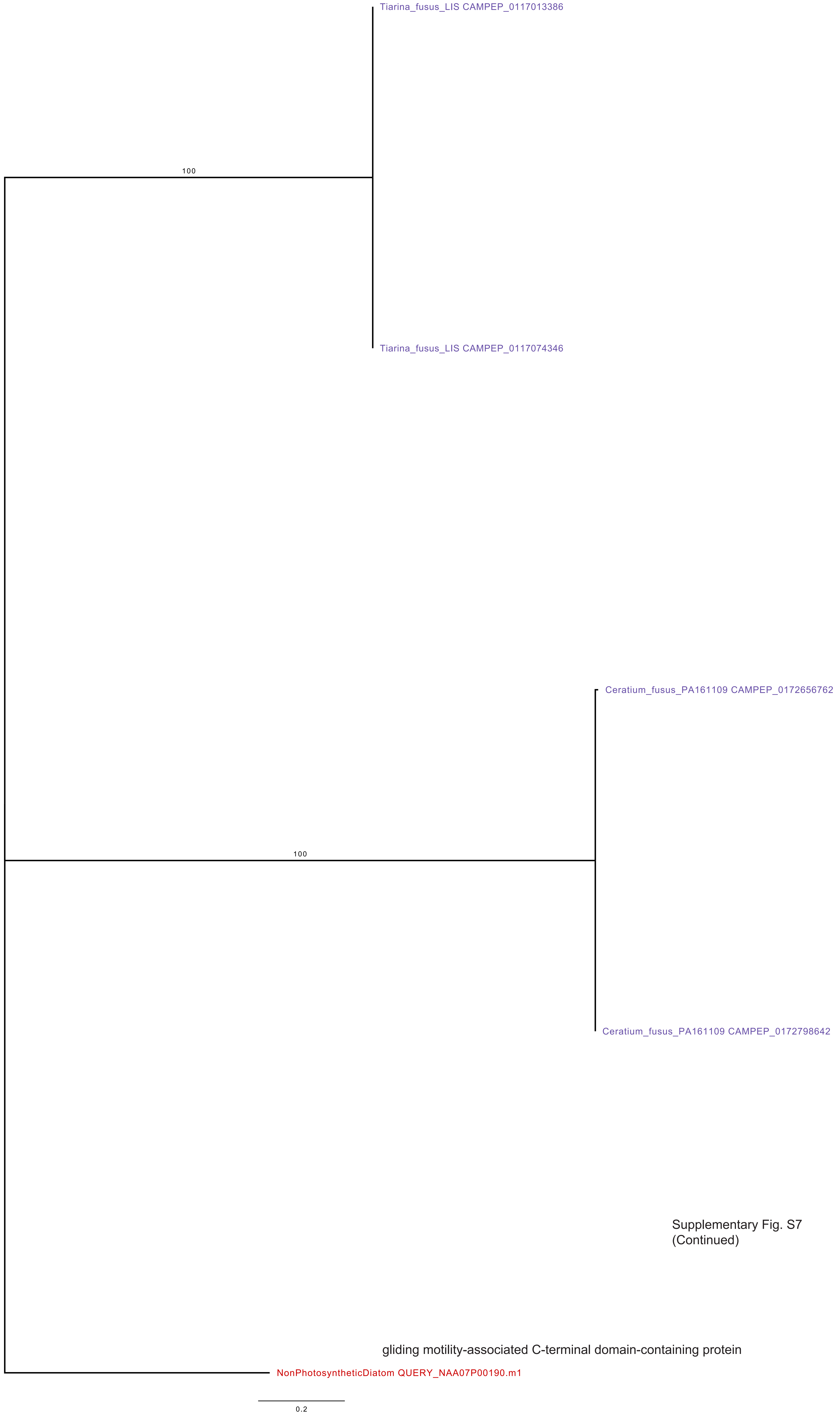

Supplementary Fig. S7
(Continued)

gliding motility-associated C-terminal domain-containing protein

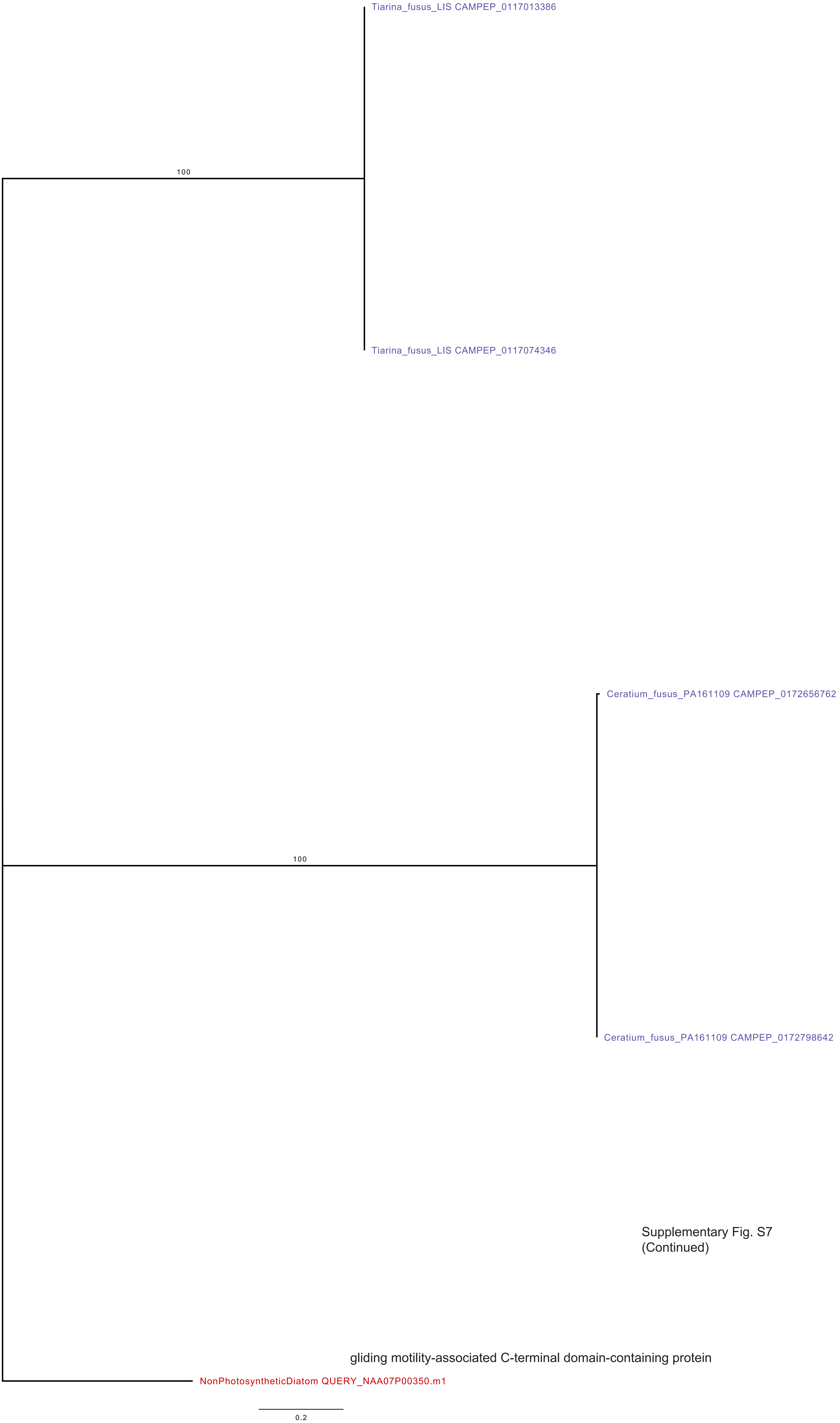

Supplementary Fig. S7
(Continued)

gliding motility-associated C-terminal domain-containing protein

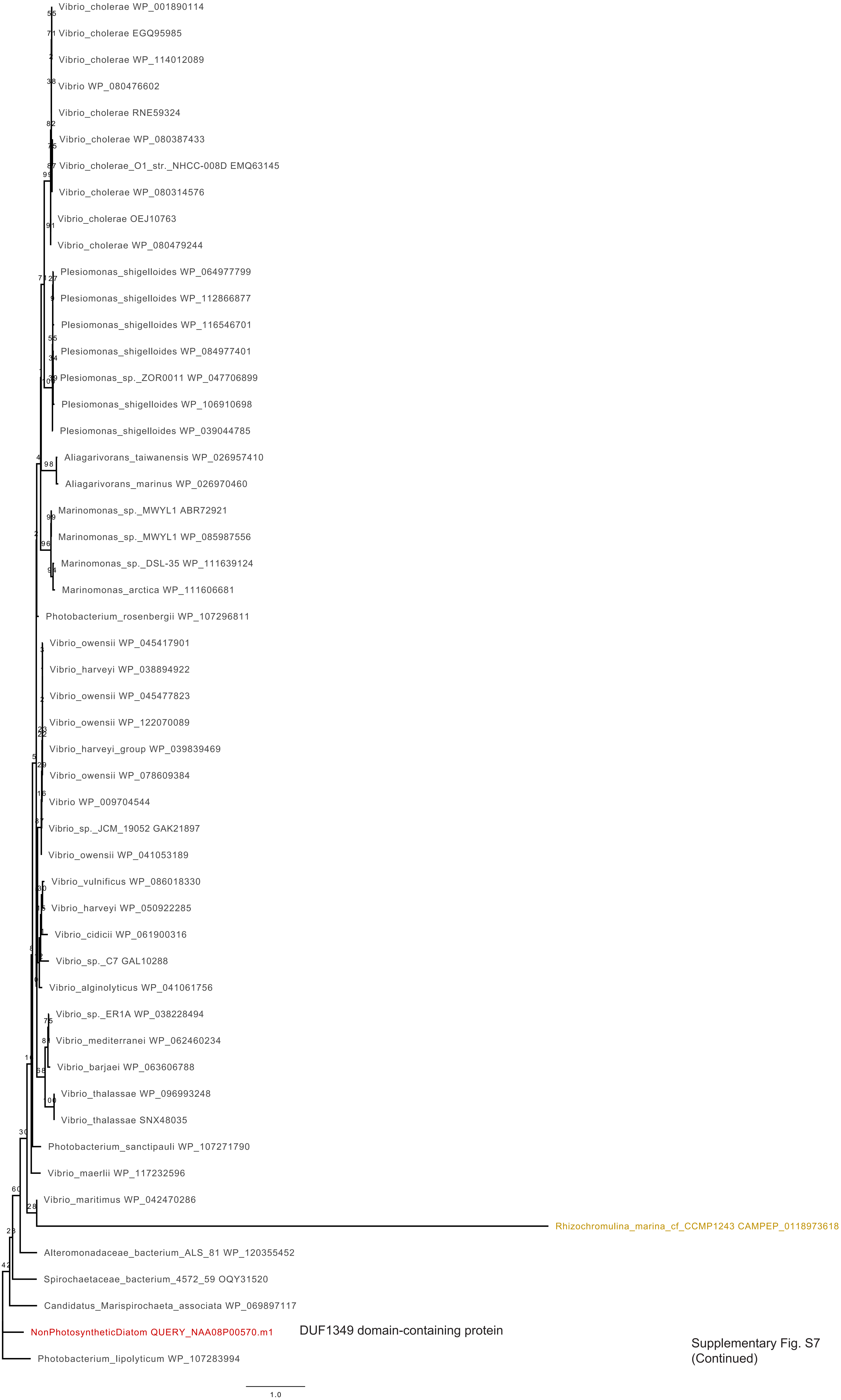

Supplementary Fig. S7
(Continued)

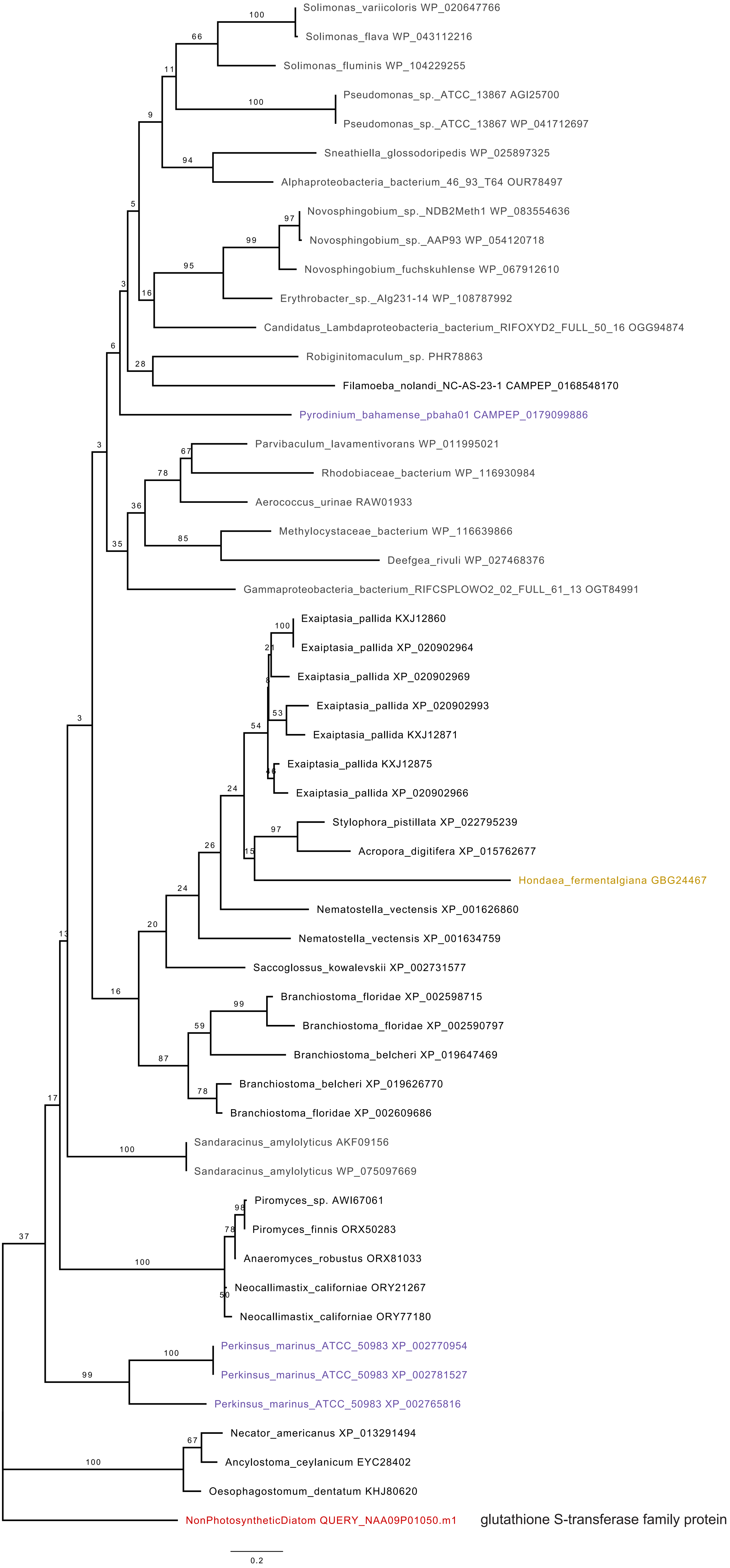

Supplementary Fig. S7
(Continued)

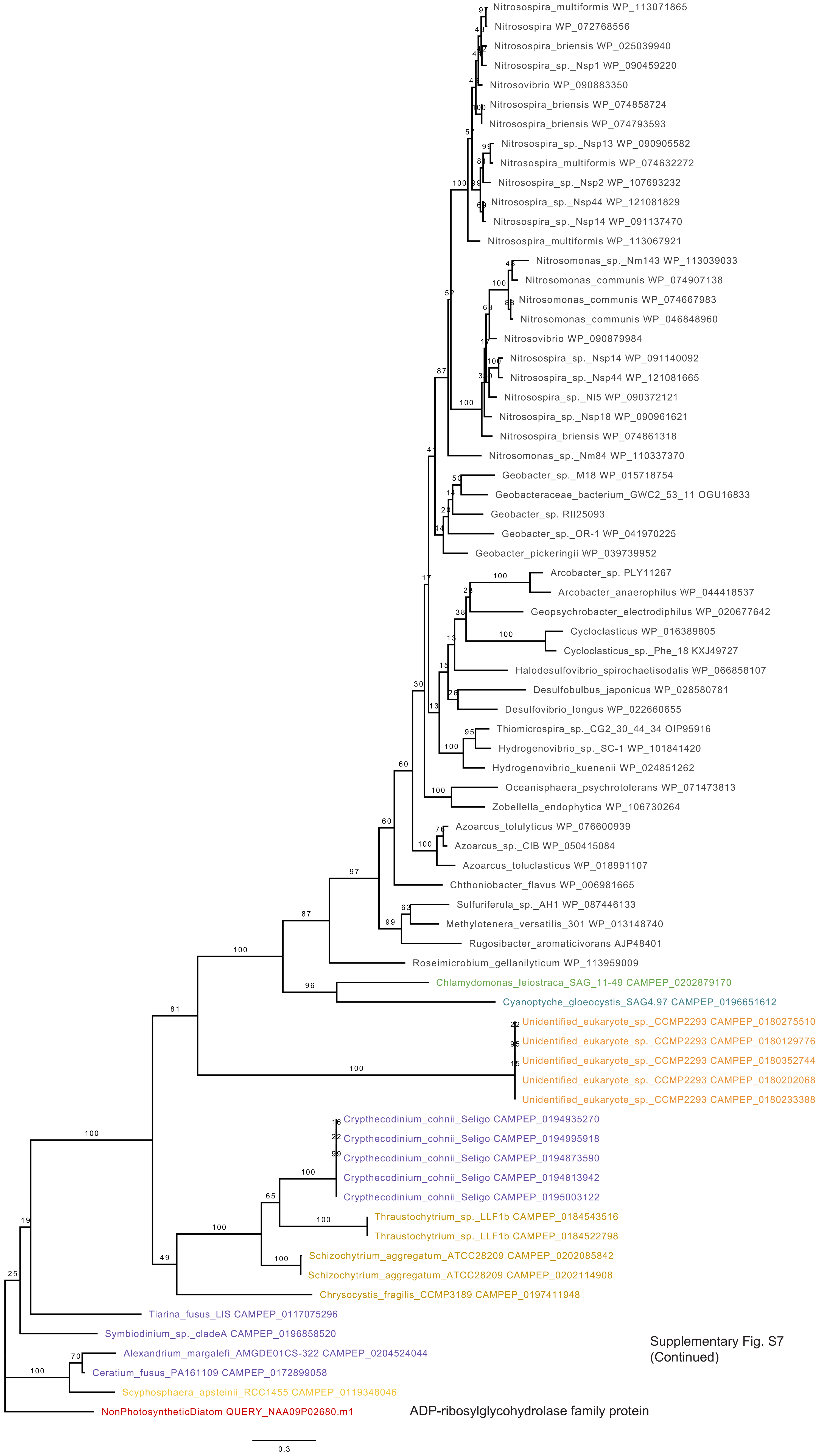

Supplementary Fig. S7
(Continued)

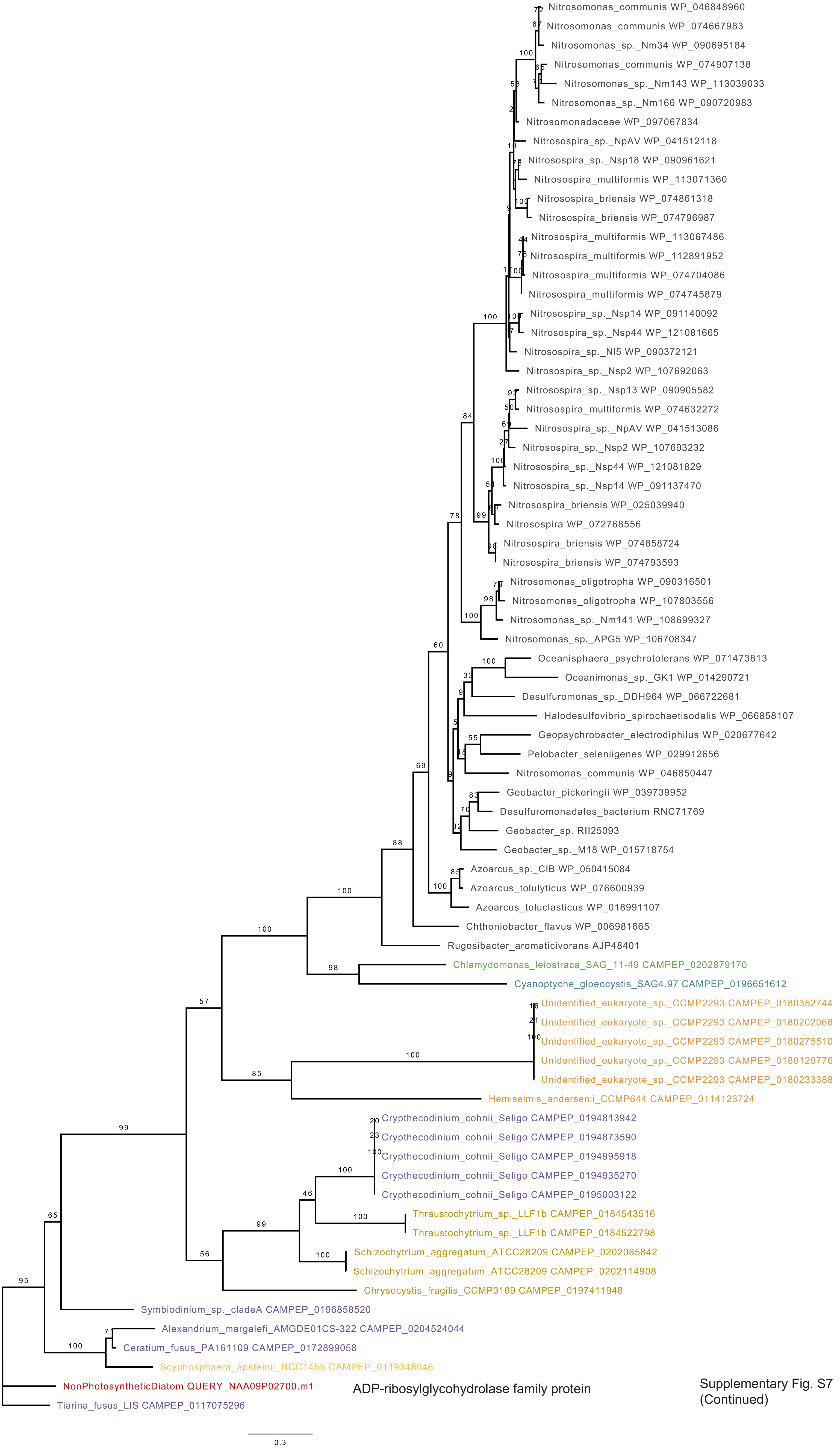

Supplementary Fig. S7 (Continued)

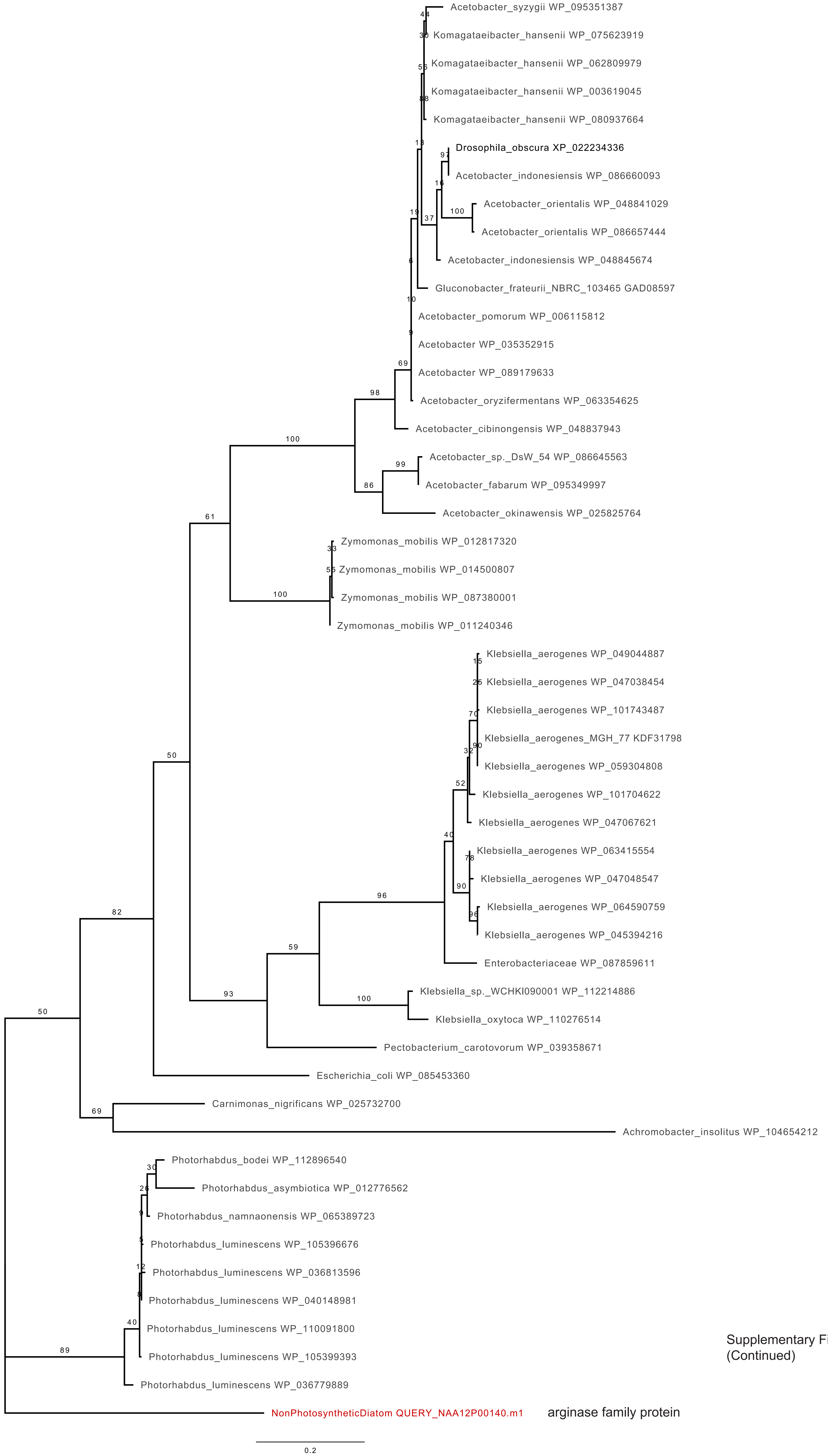

Supplementary Fig. S7
(Continued)

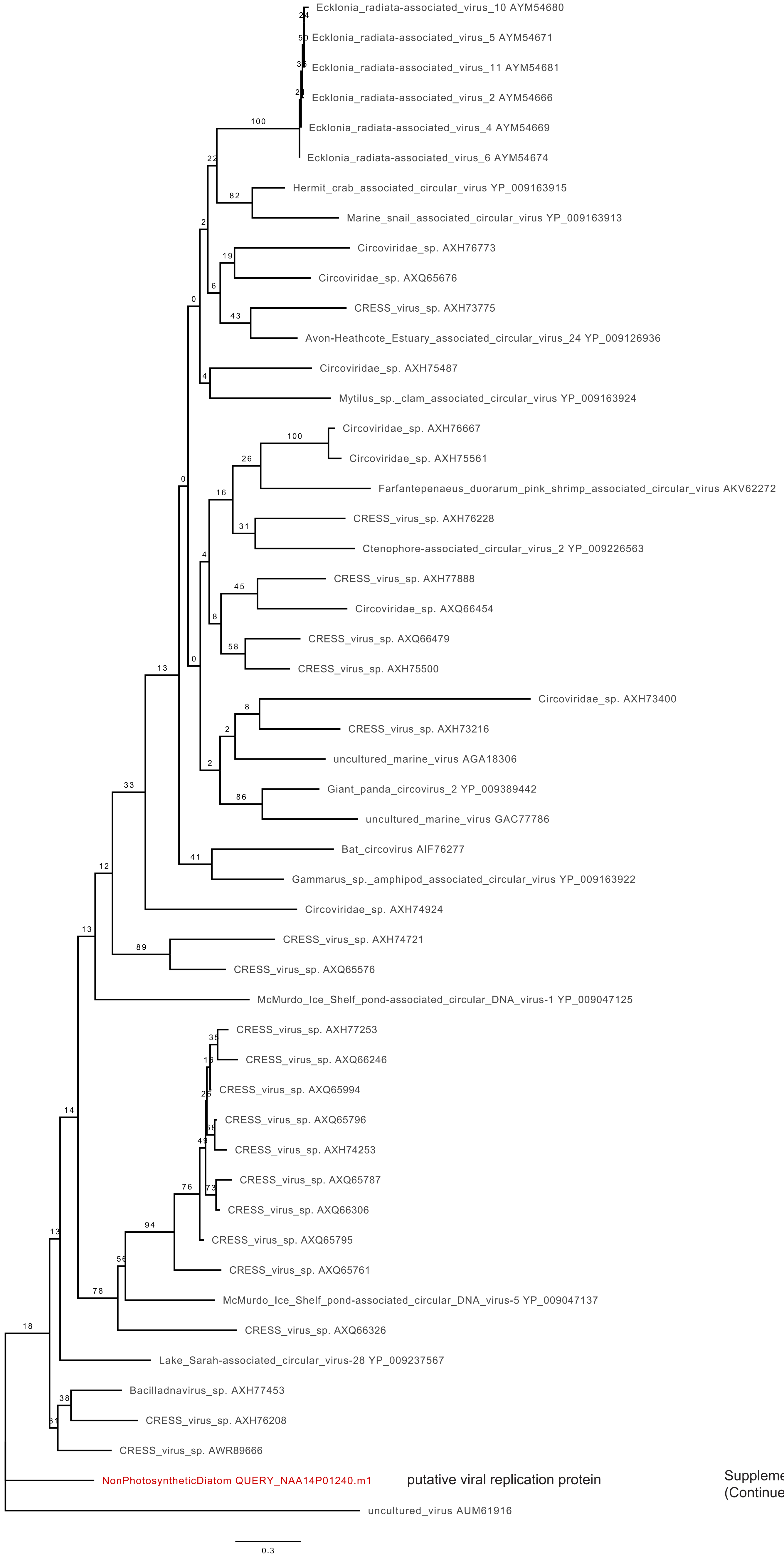

Supplementary Fig. S7 (Continued)

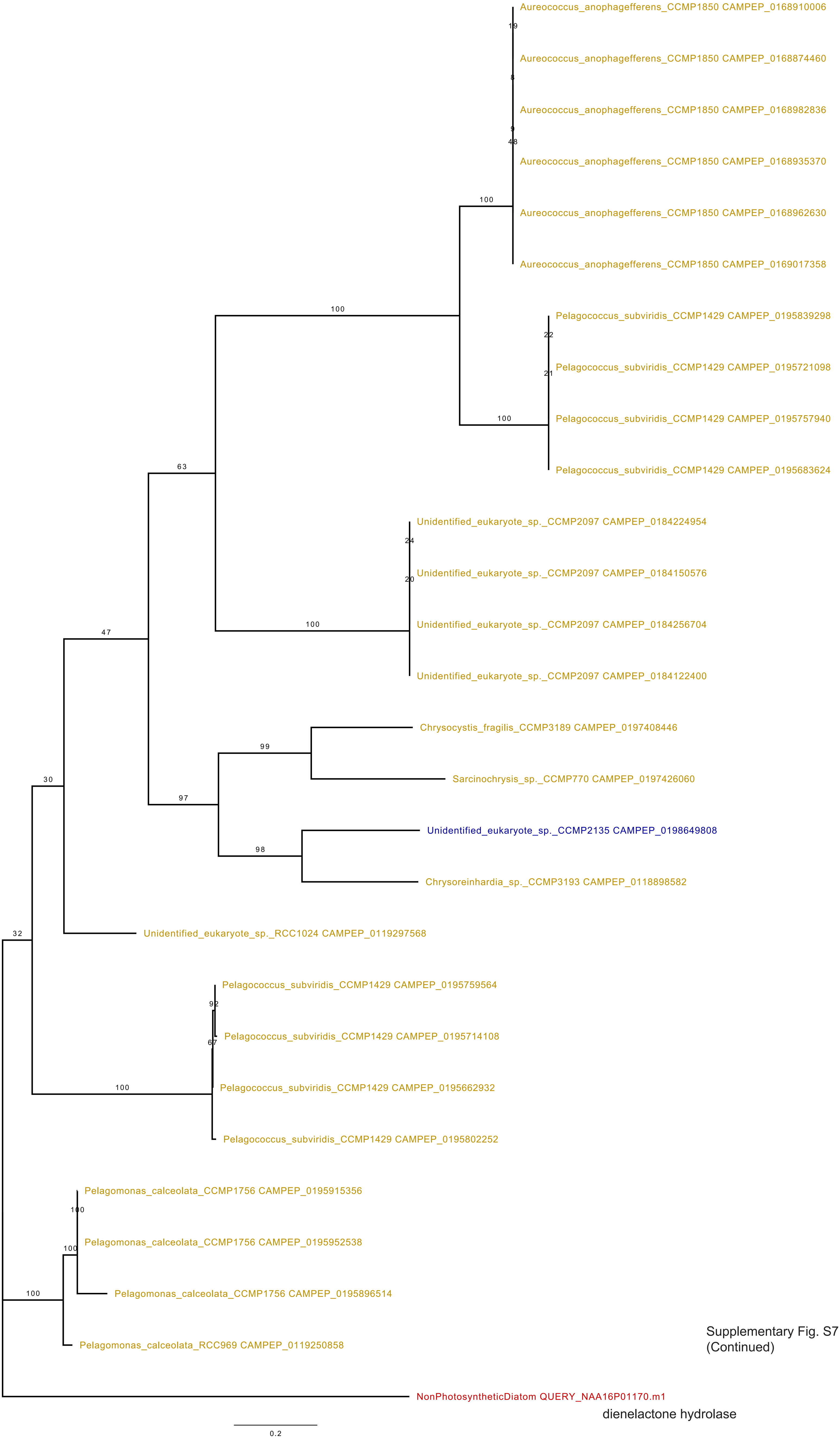

Supplementary Fig. S7
(Continued)

dienelactone hydrolase

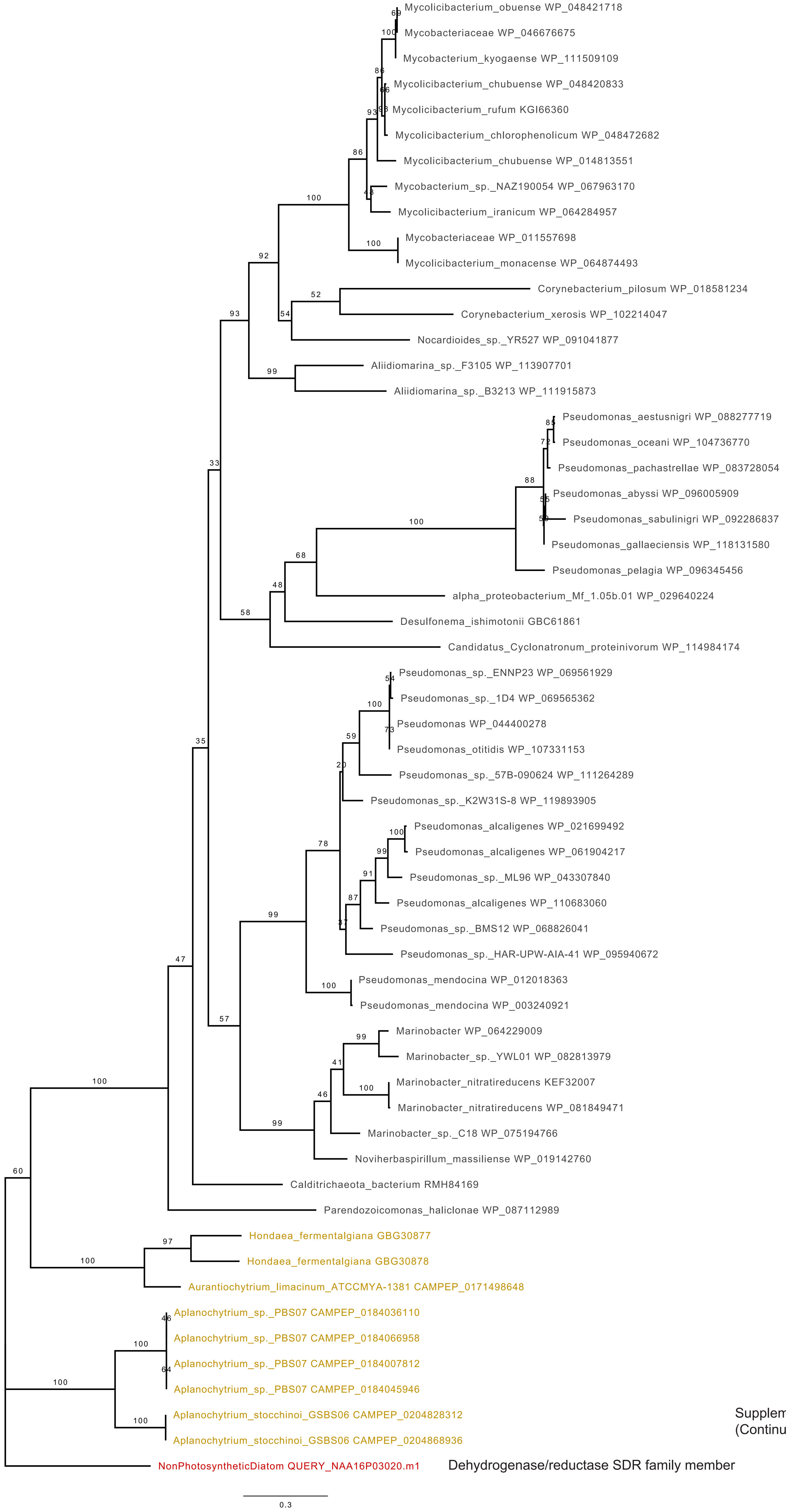

Supplementary Fig. S7
(Continued)

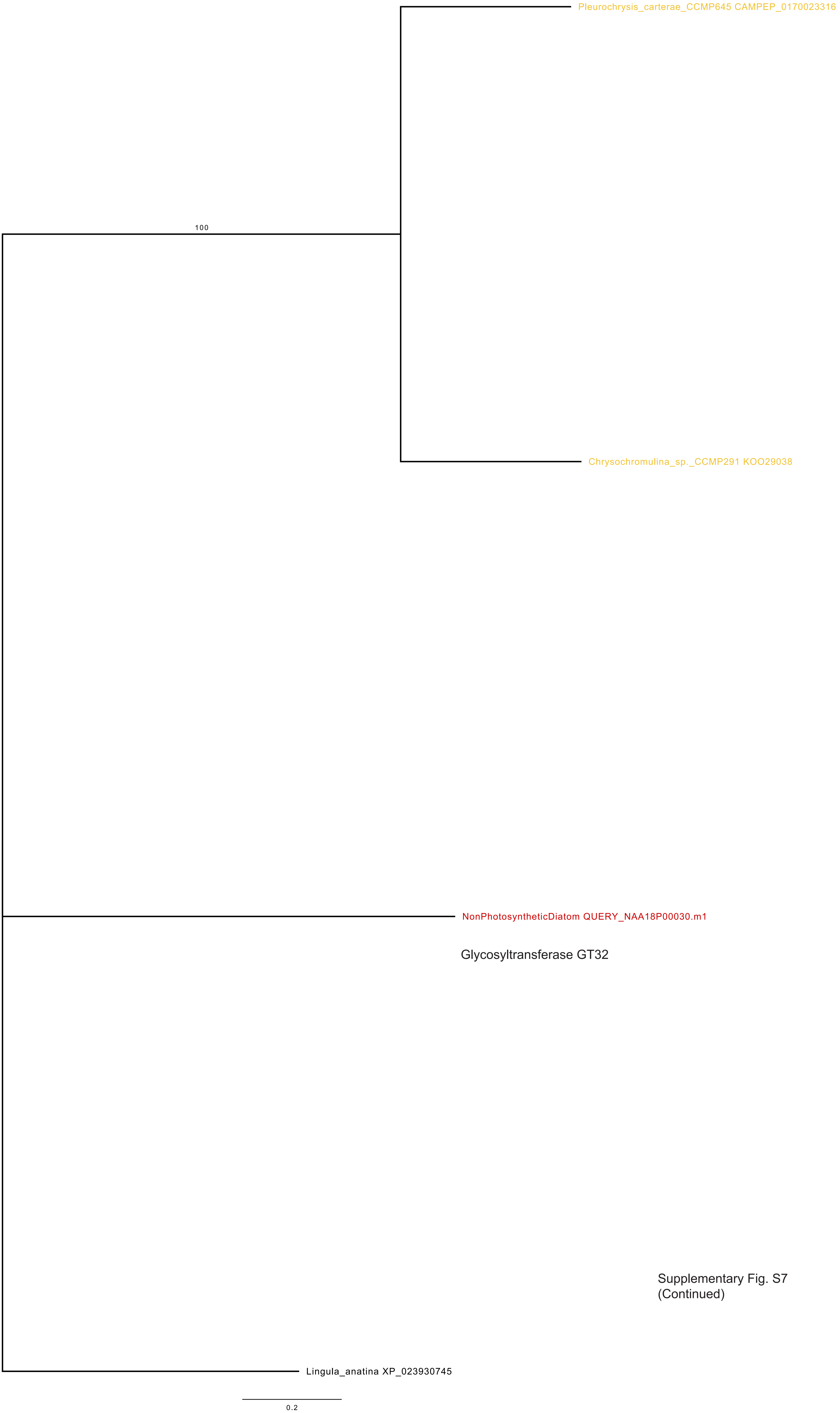

Supplementary Fig. S7
(Continued)

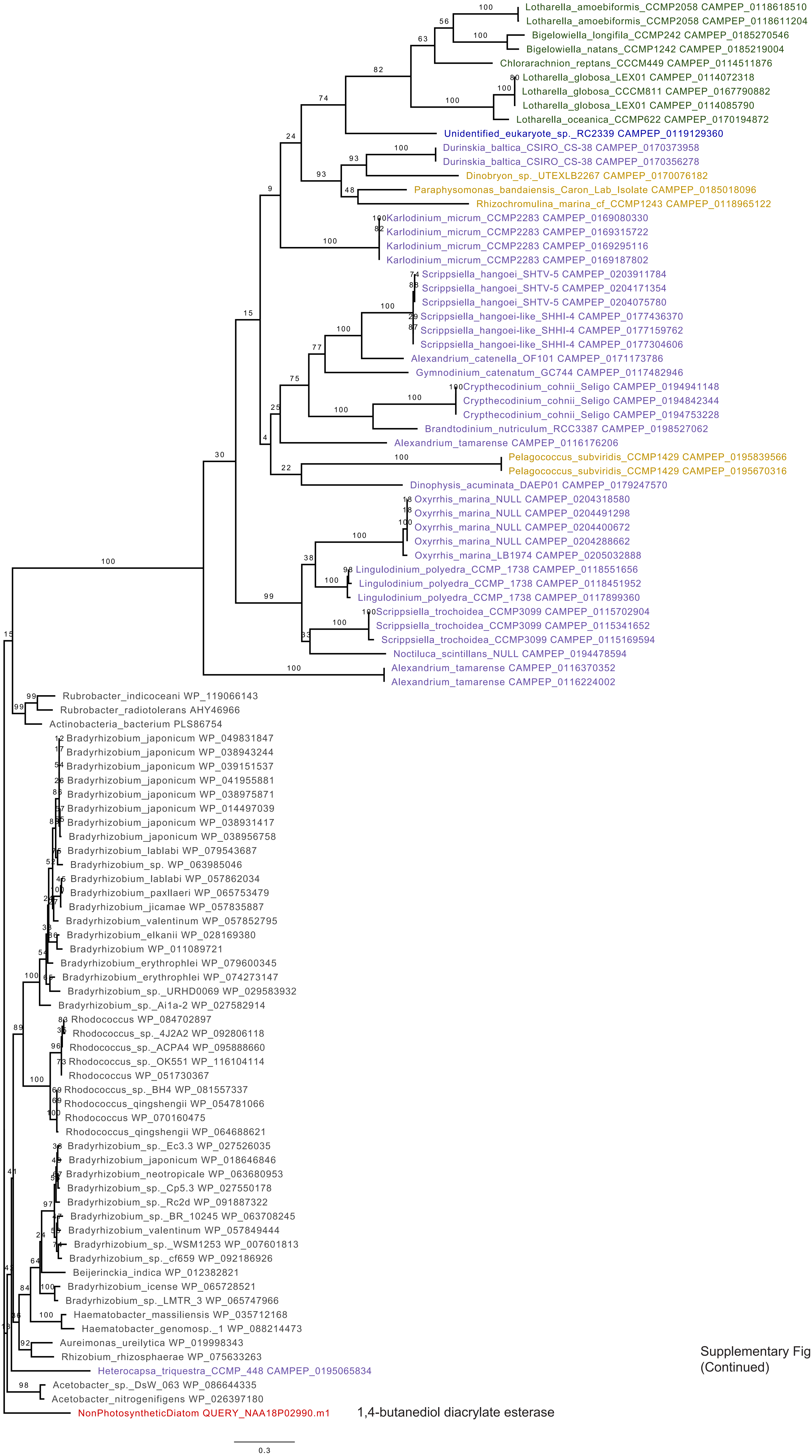

Supplementary Fig. S7
(Continued)

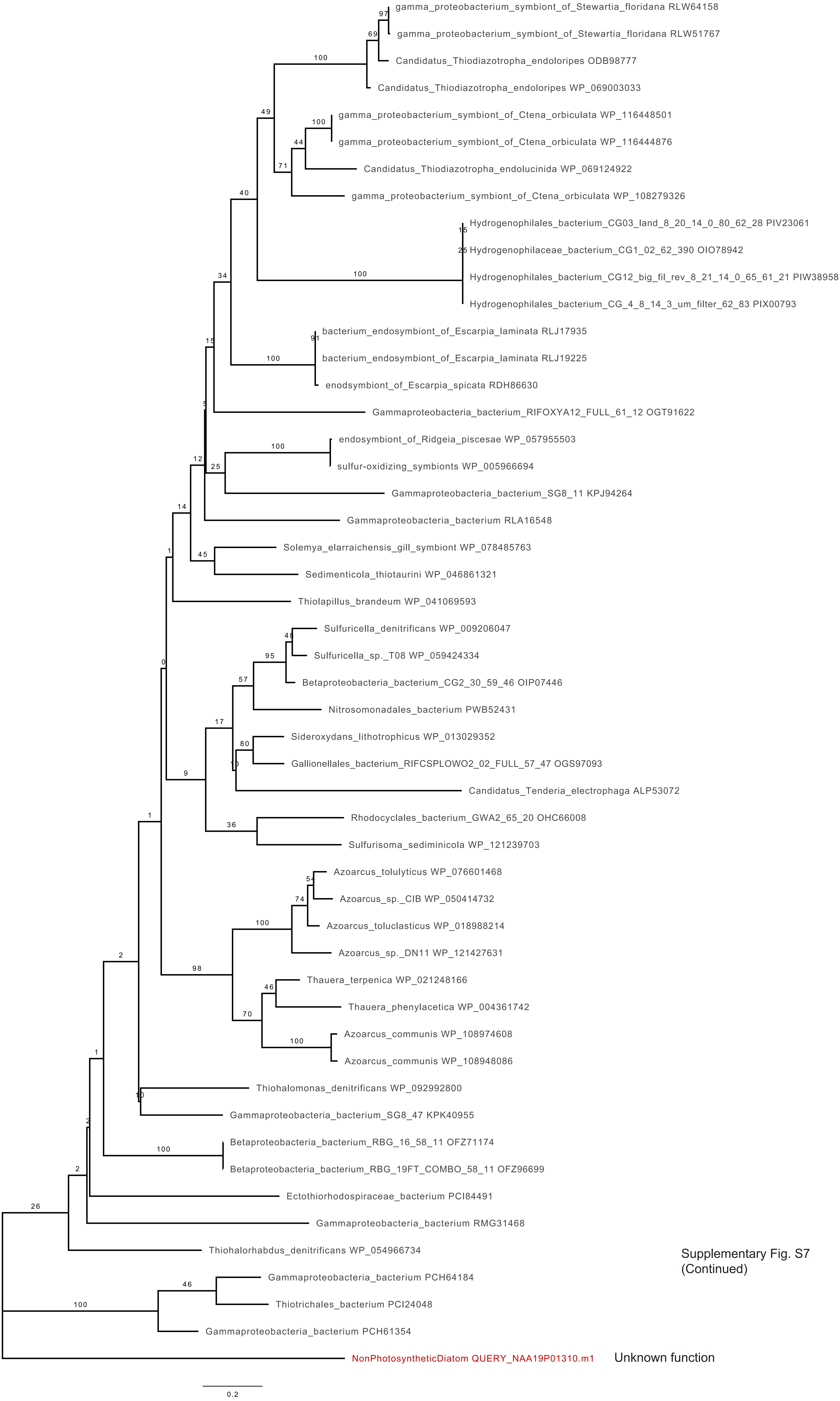

Supplementary Fig. S7
(Continued)

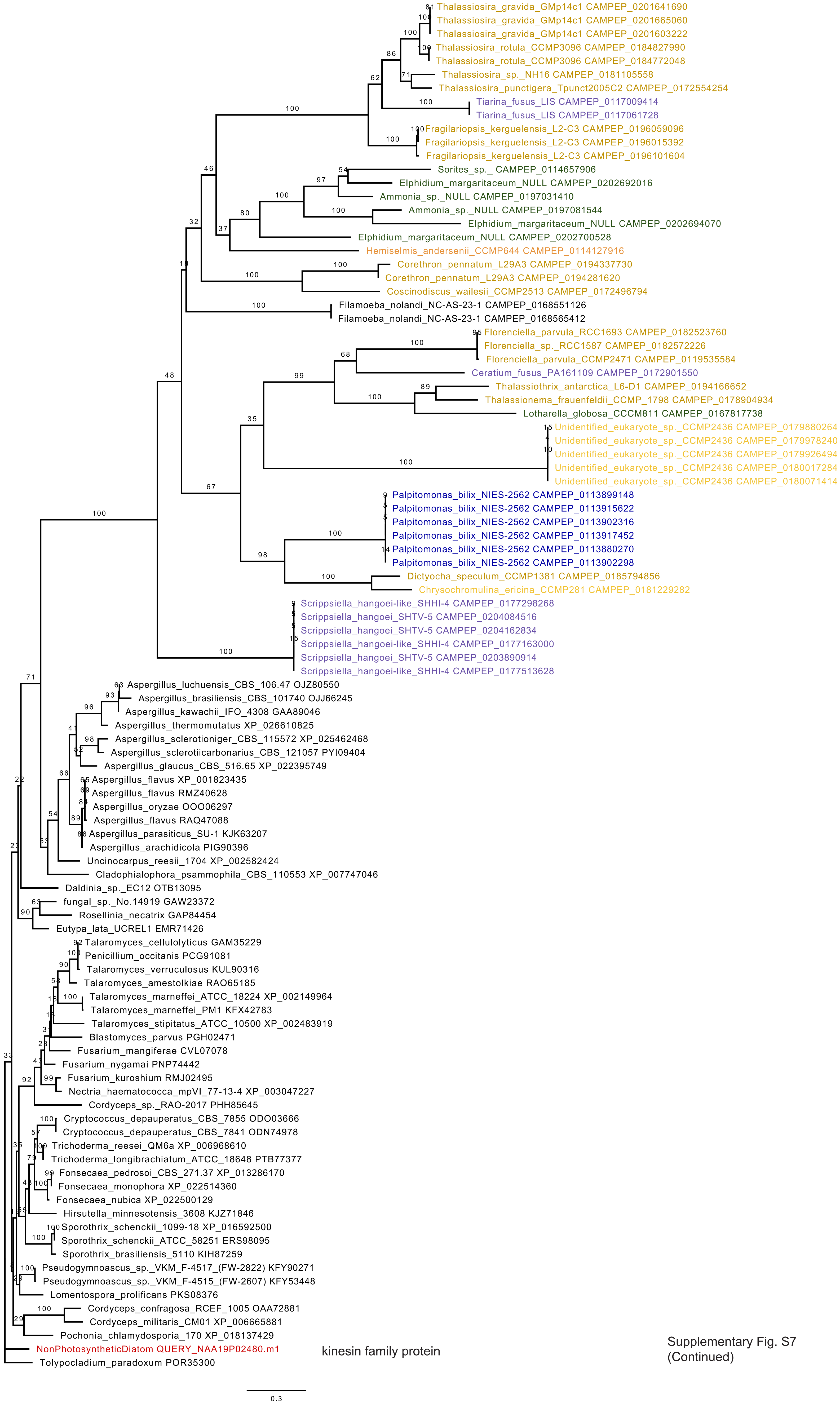

Supplementary Fig. S7
(Continued)

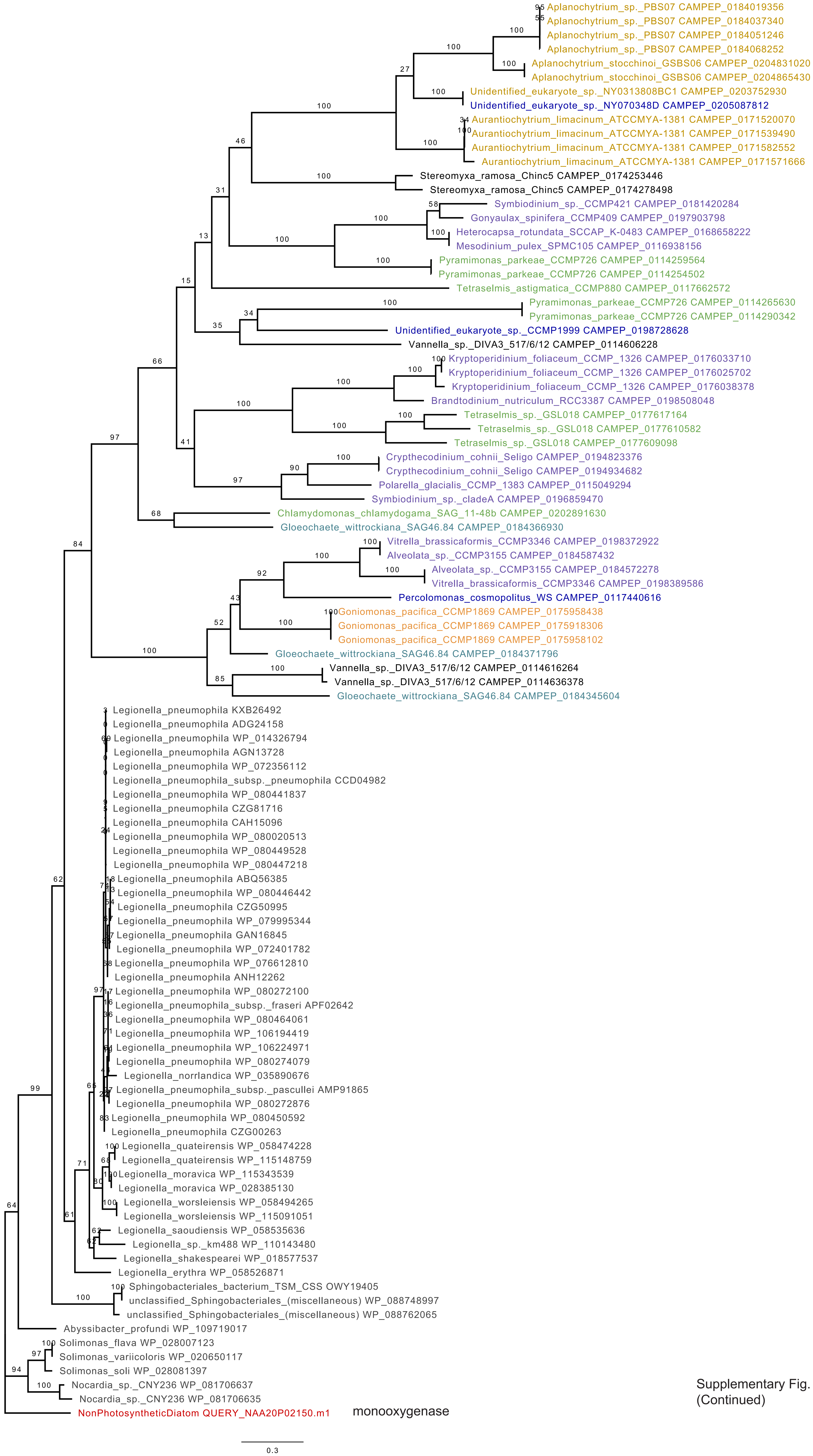

Supplementary Fig. S7
(Continued)

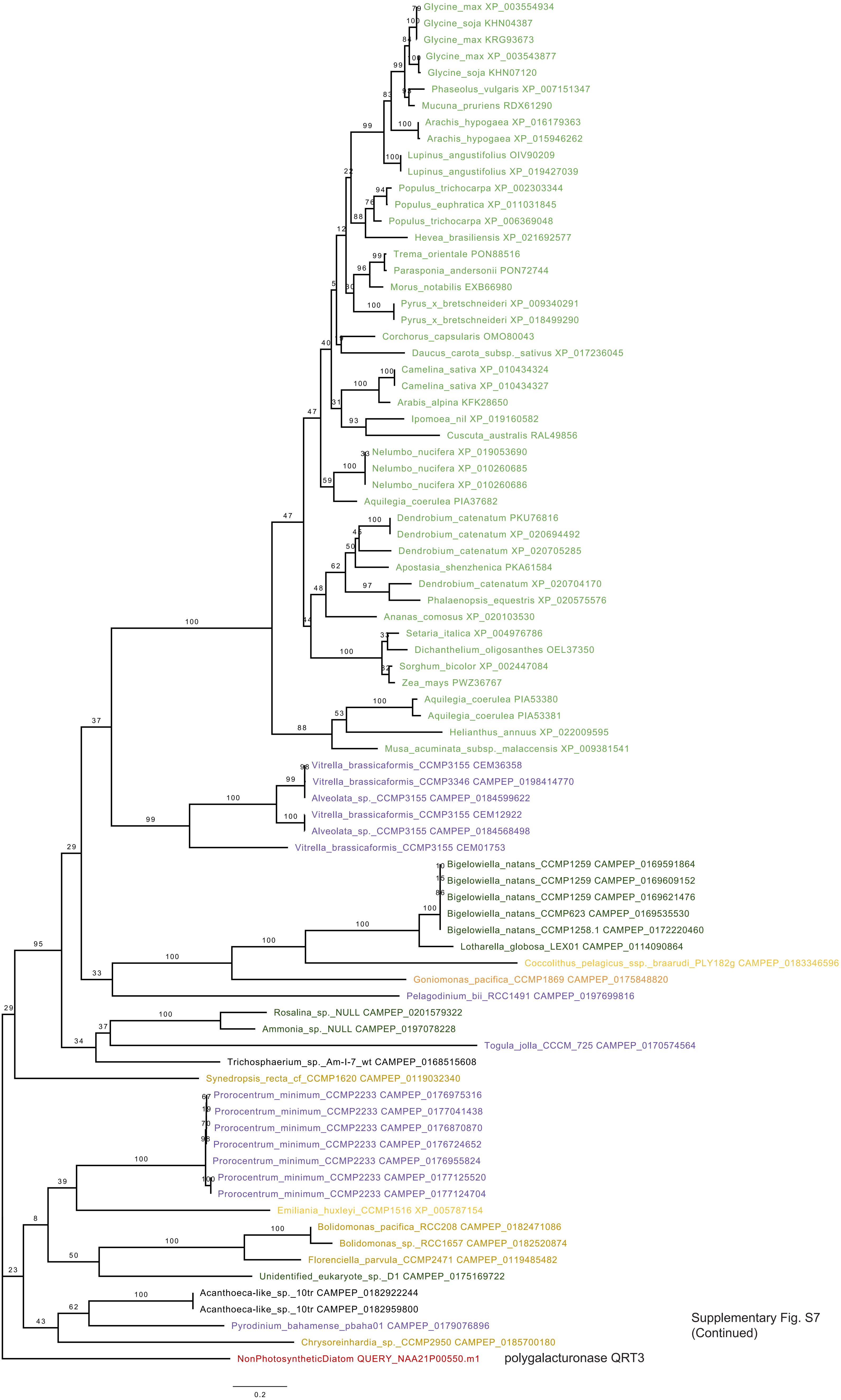

Supplementary Fig. S7
(Continued)

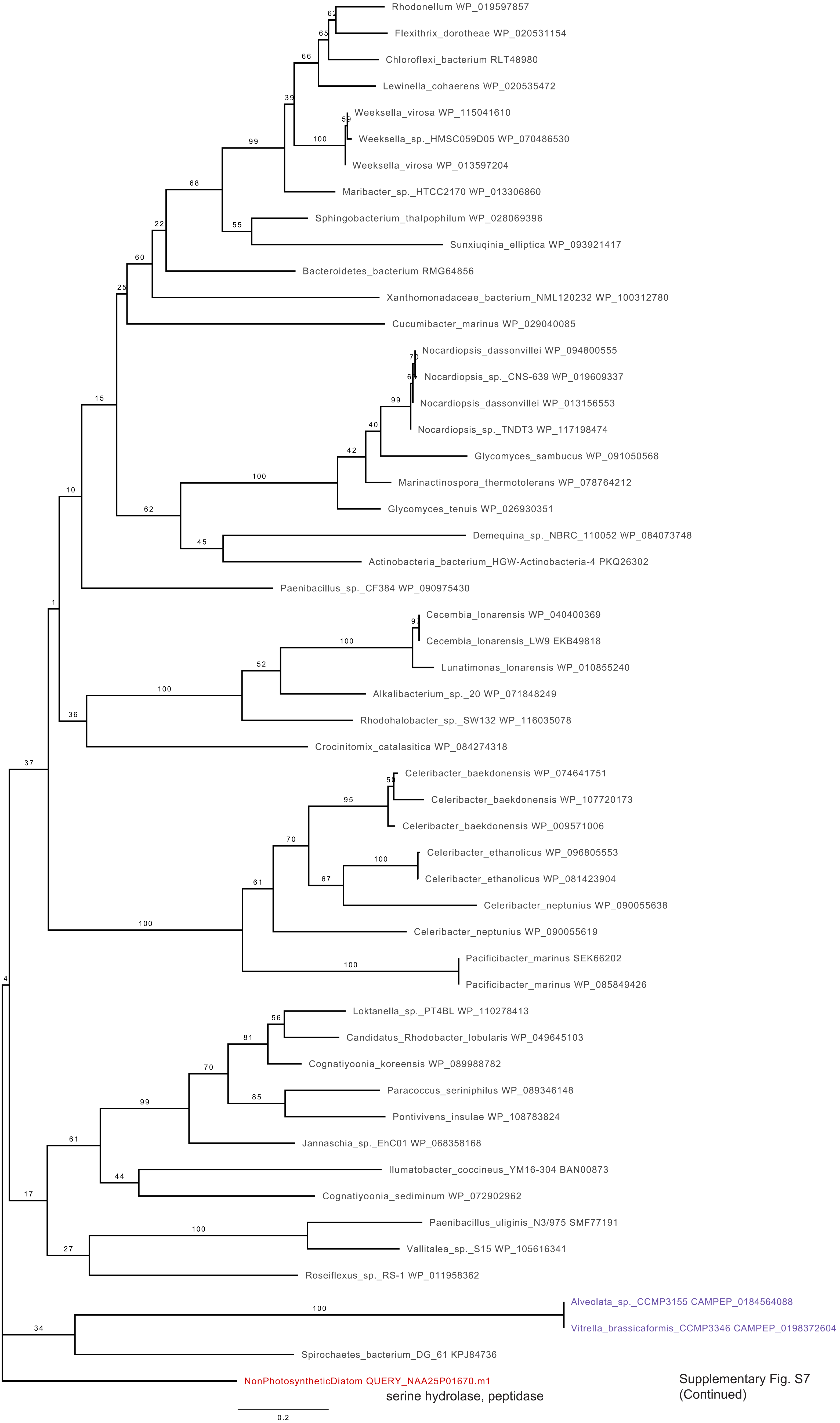

Supplementary Fig. S7
(Continued)

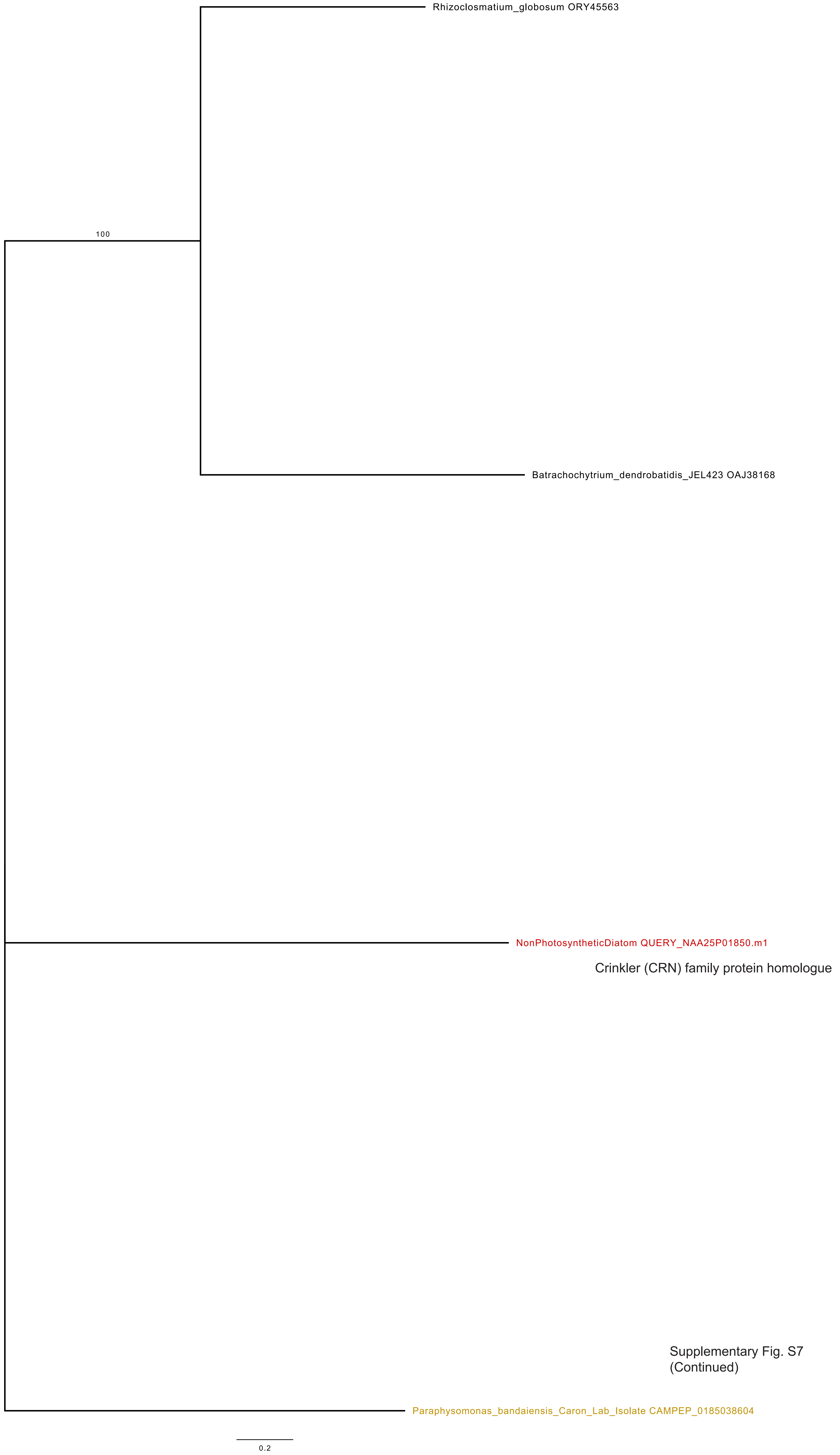

Supplementary Fig. S7
(Continued)

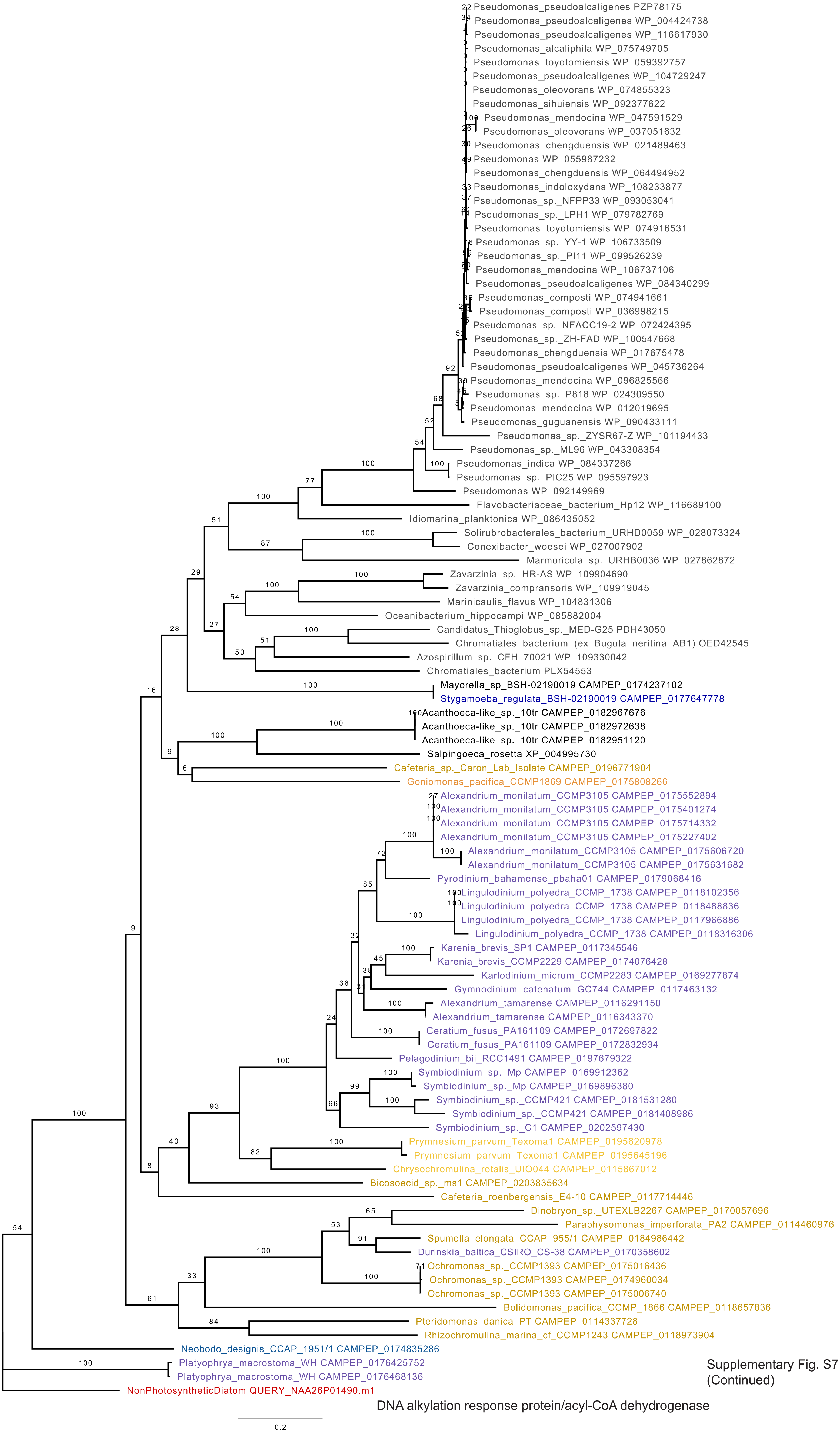

Supplementary Fig. S7
(Continued)

DNA alkylation response protein/acyl-CoA dehydrogenase

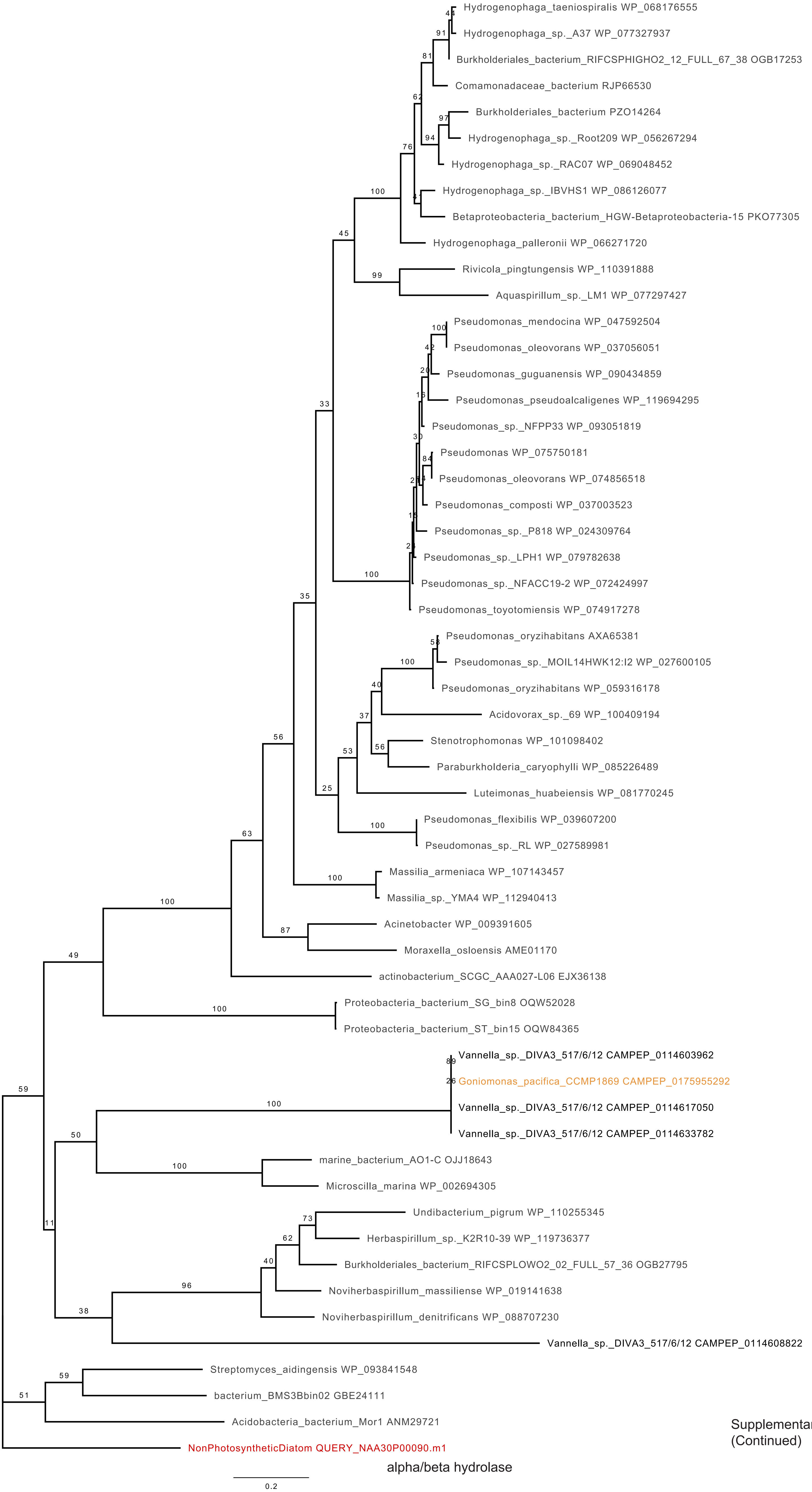

alpha/beta hydrolase

Supplementary Fig. S7
(Continued)

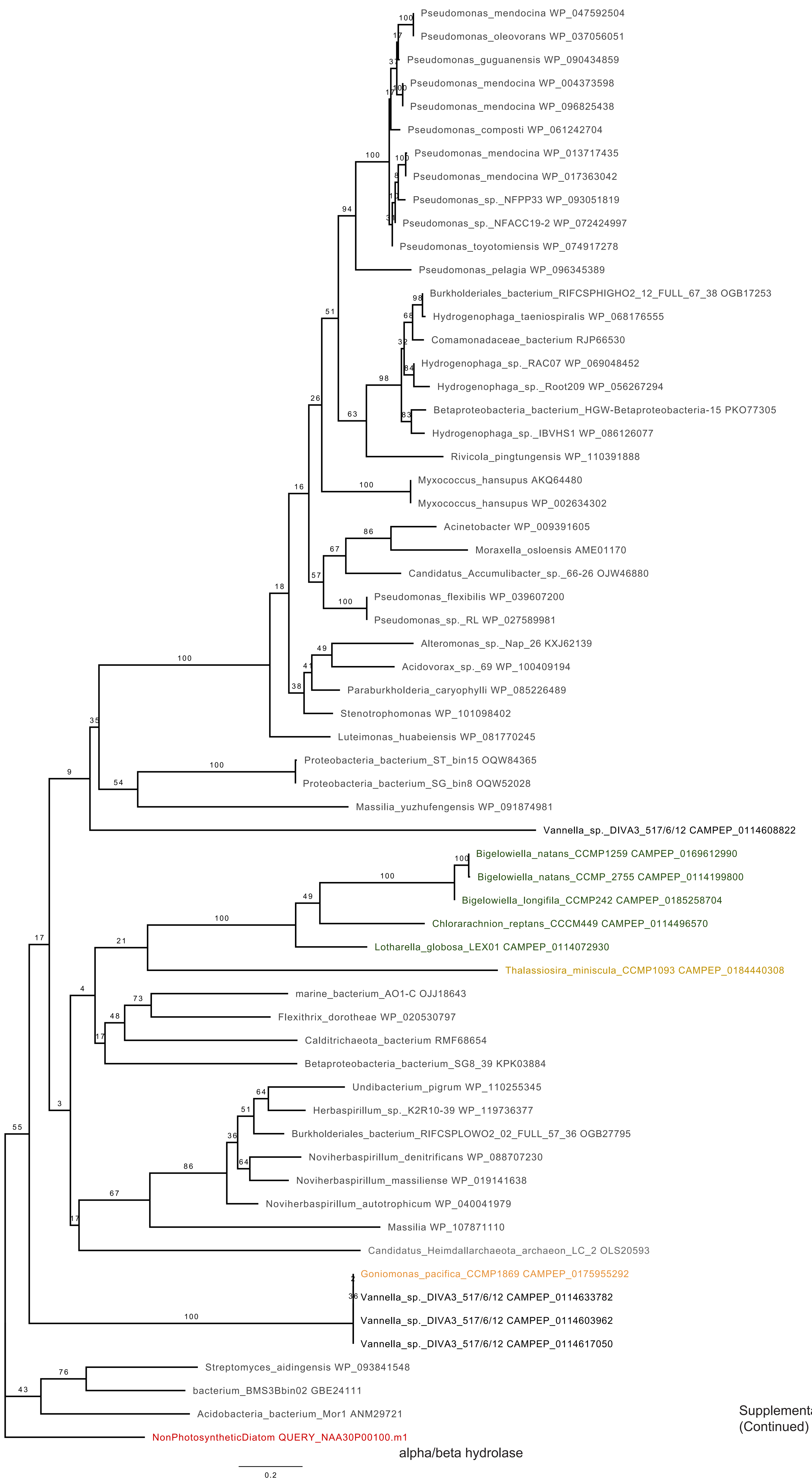

alpha/beta hydrolase

Supplementary Fig. S7
(Continued)

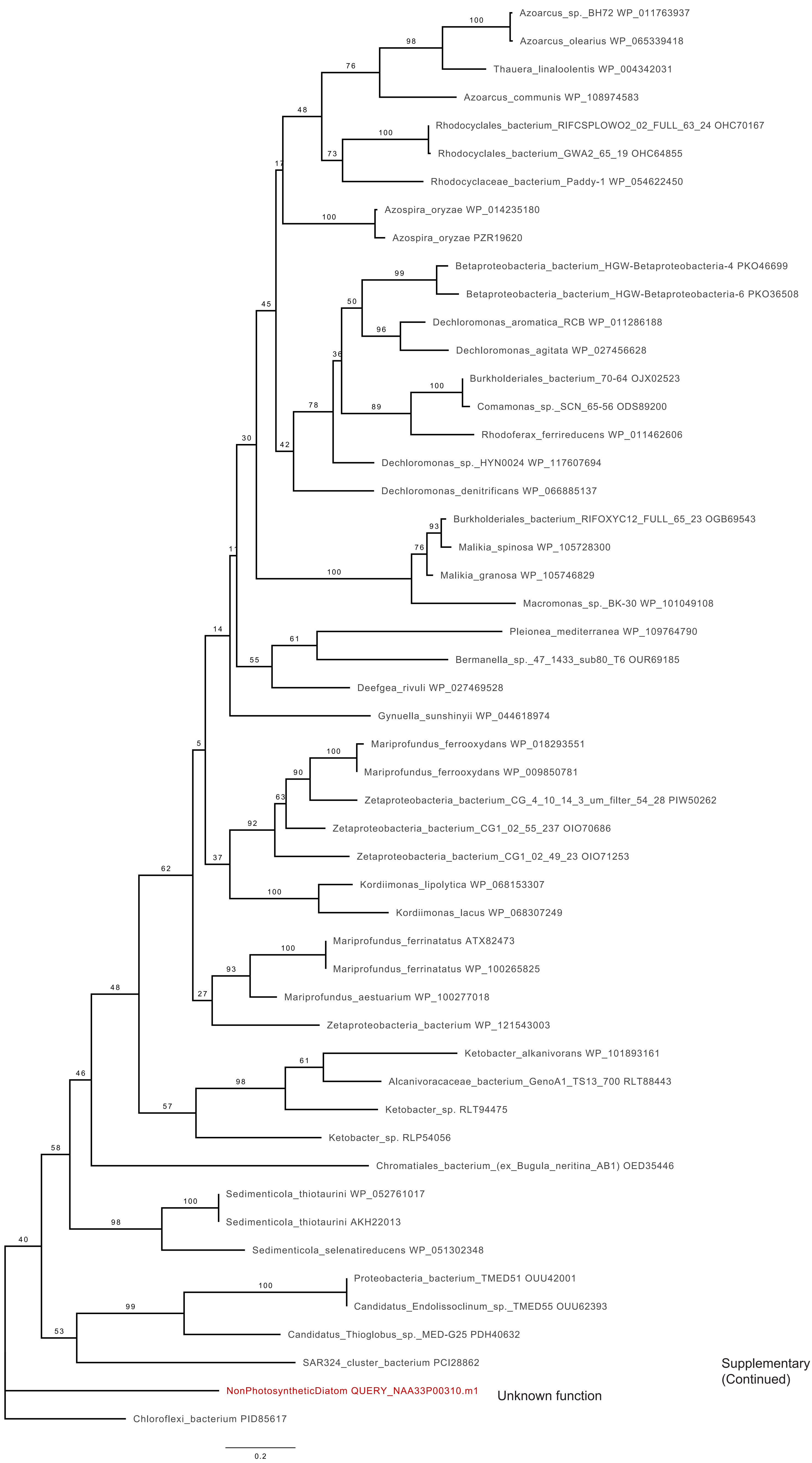

Supplementary Fig. S7
(Continued)

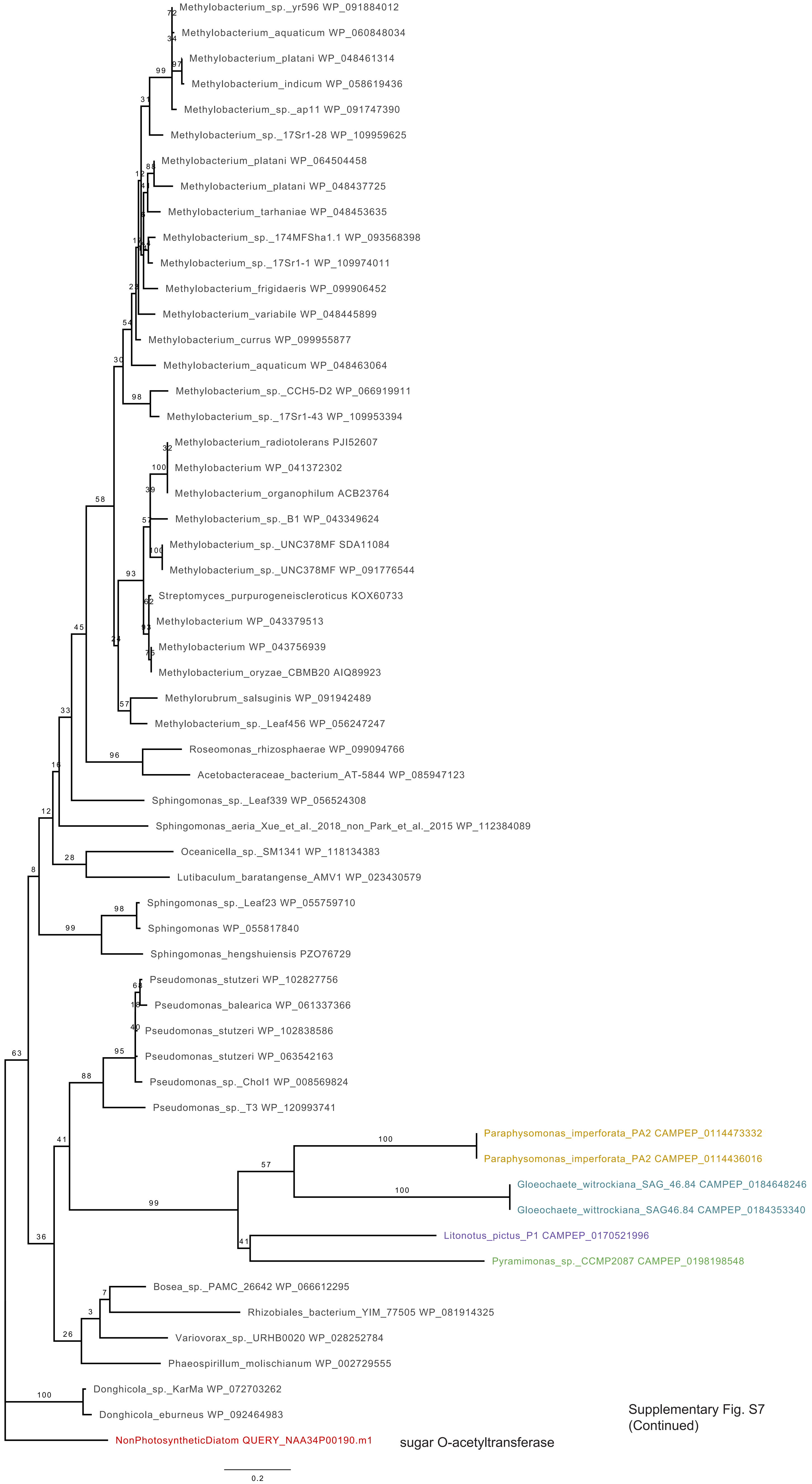

Supplementary Fig. S7
(Continued)

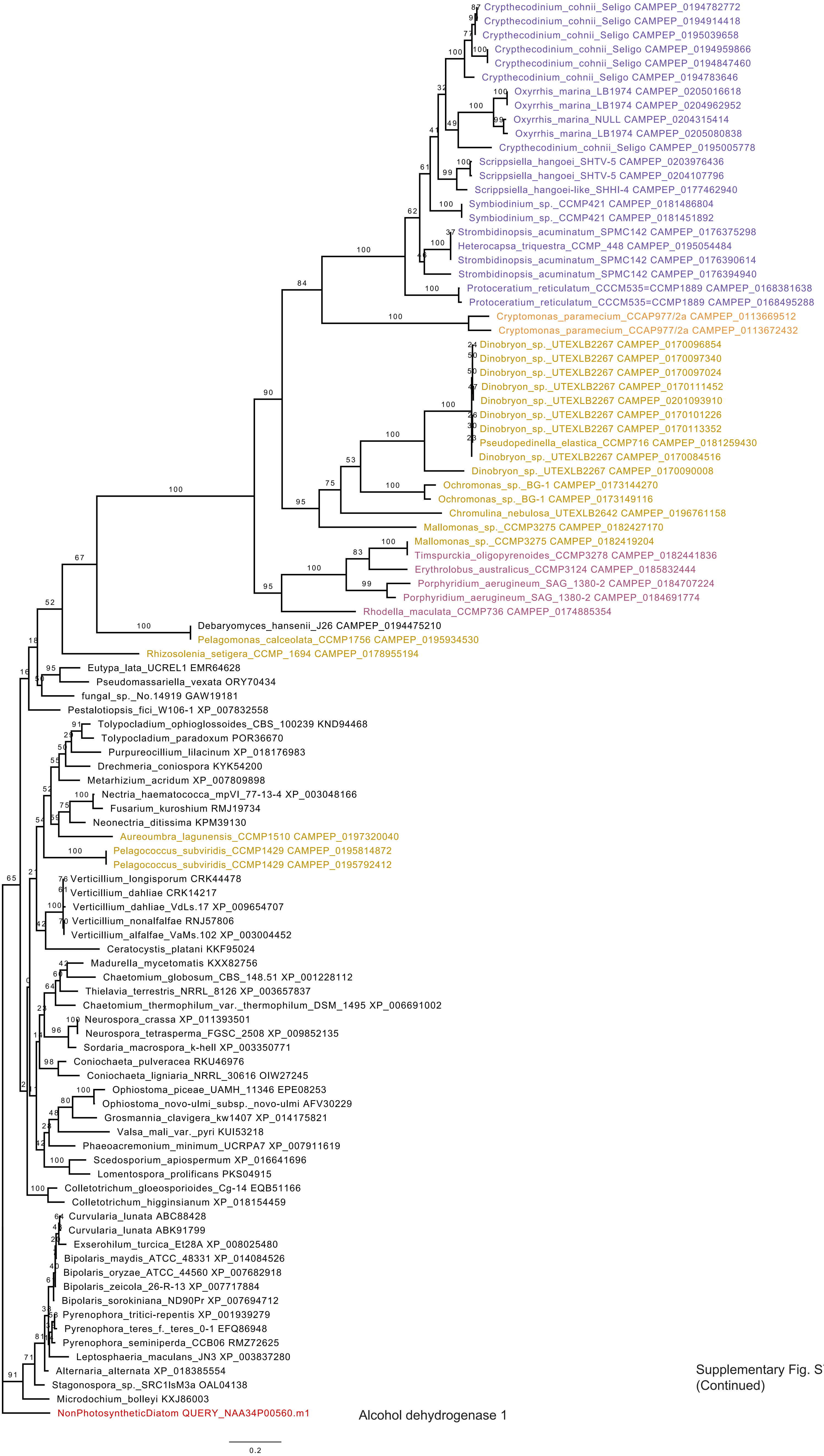

Supplementary Fig. S7
(Continued)

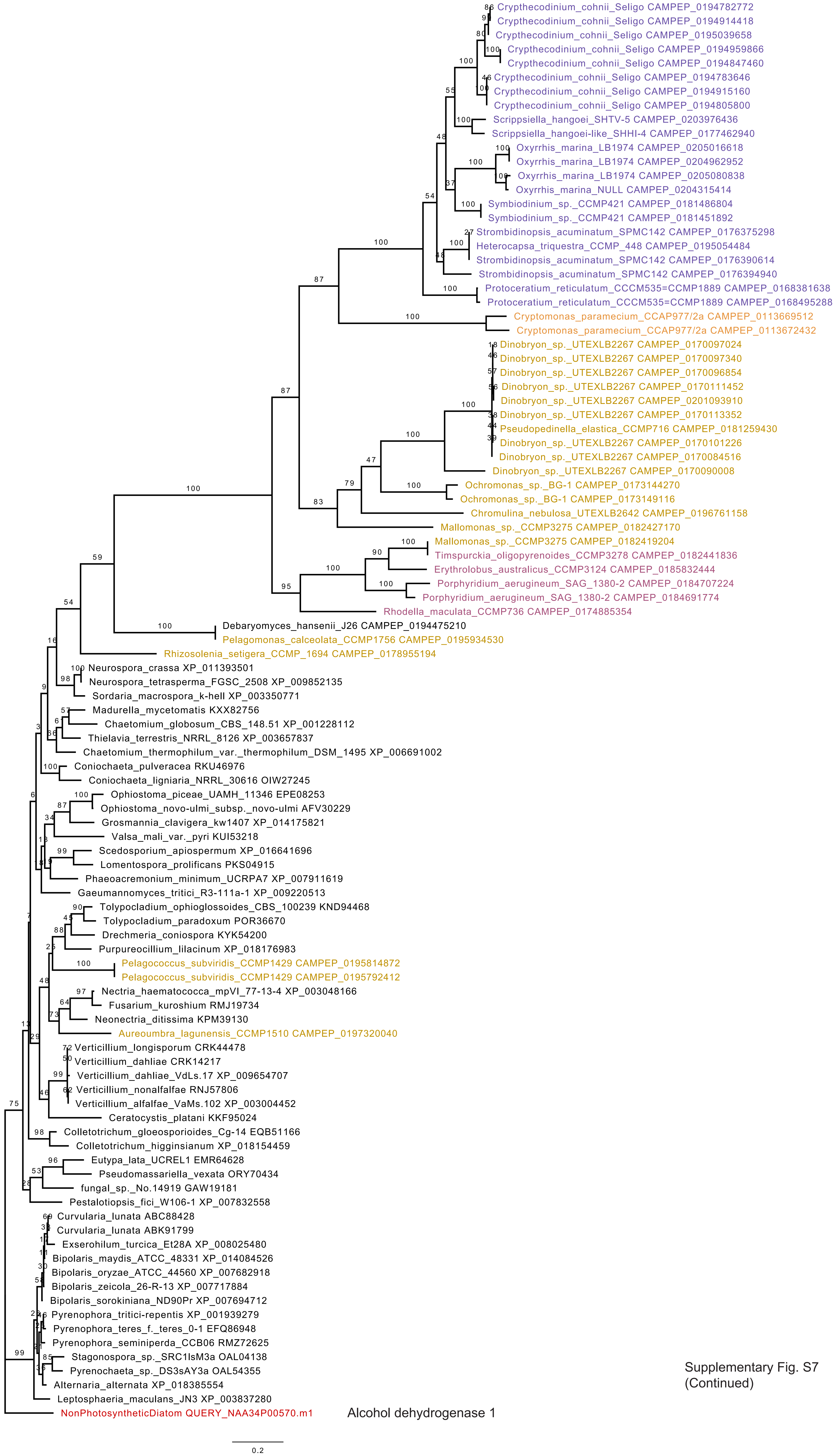

Supplementary Fig. S7
(Continued)

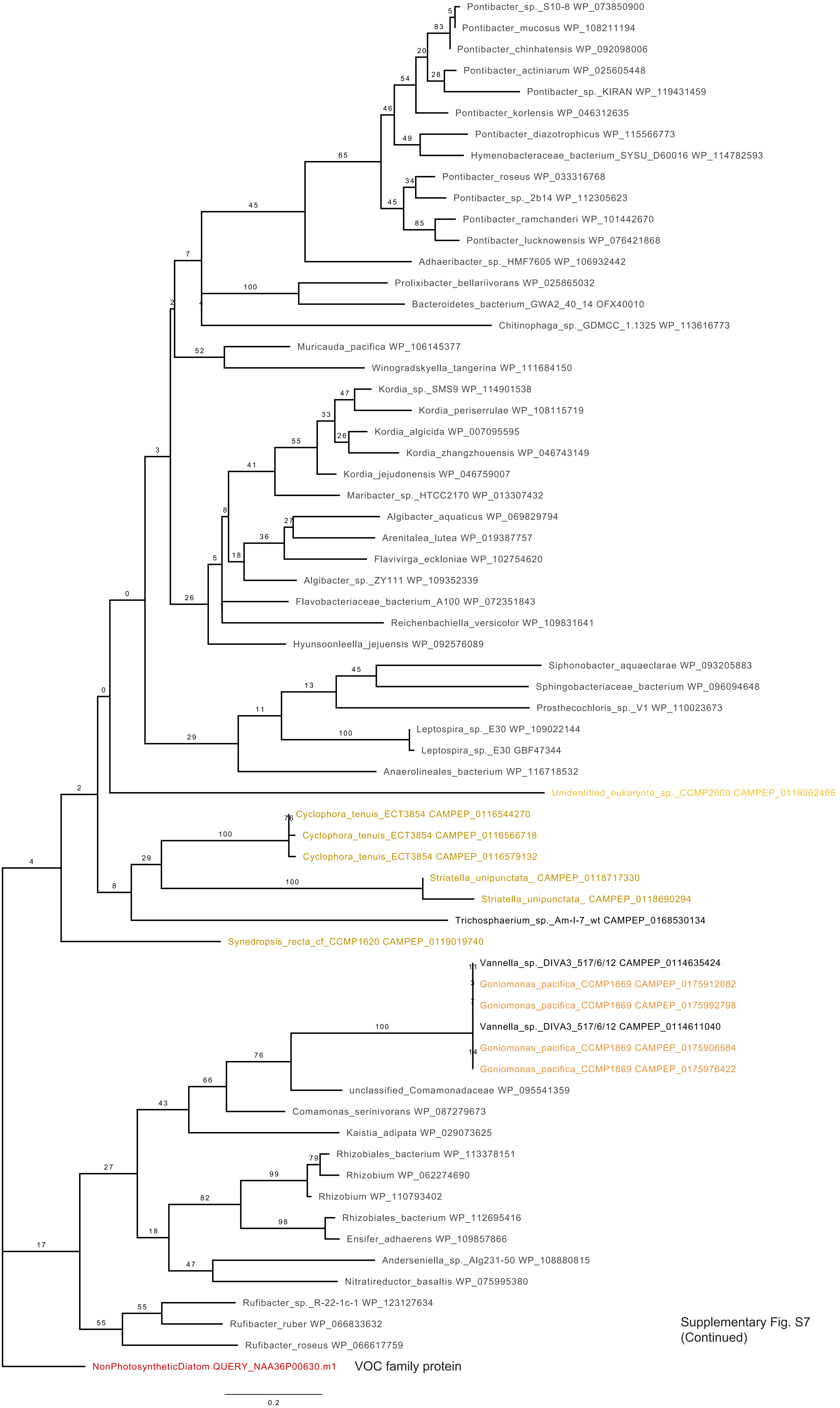

Supplementary Fig. S7
(Continued)

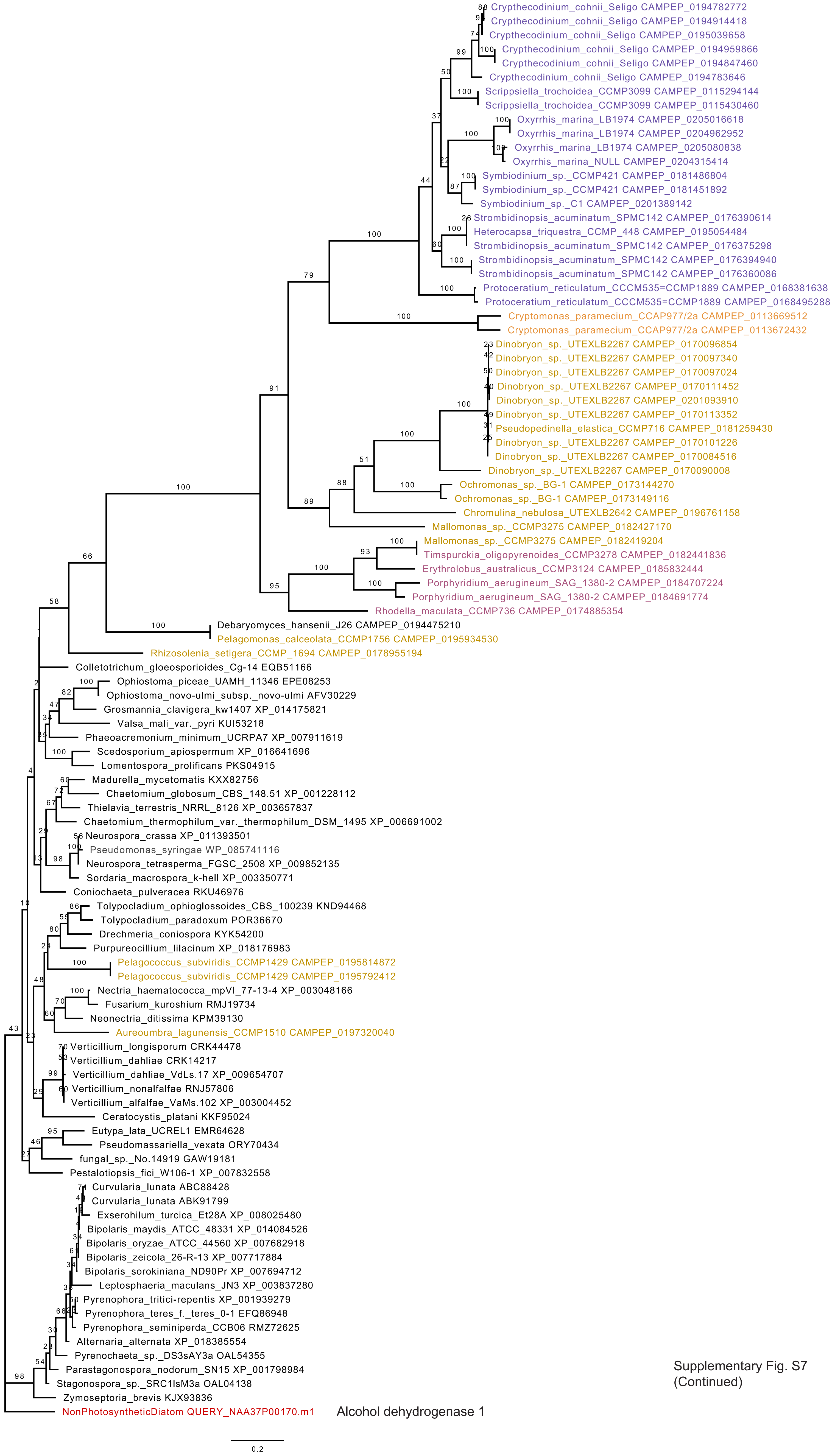

Supplementary Fig. S7 (Continued)

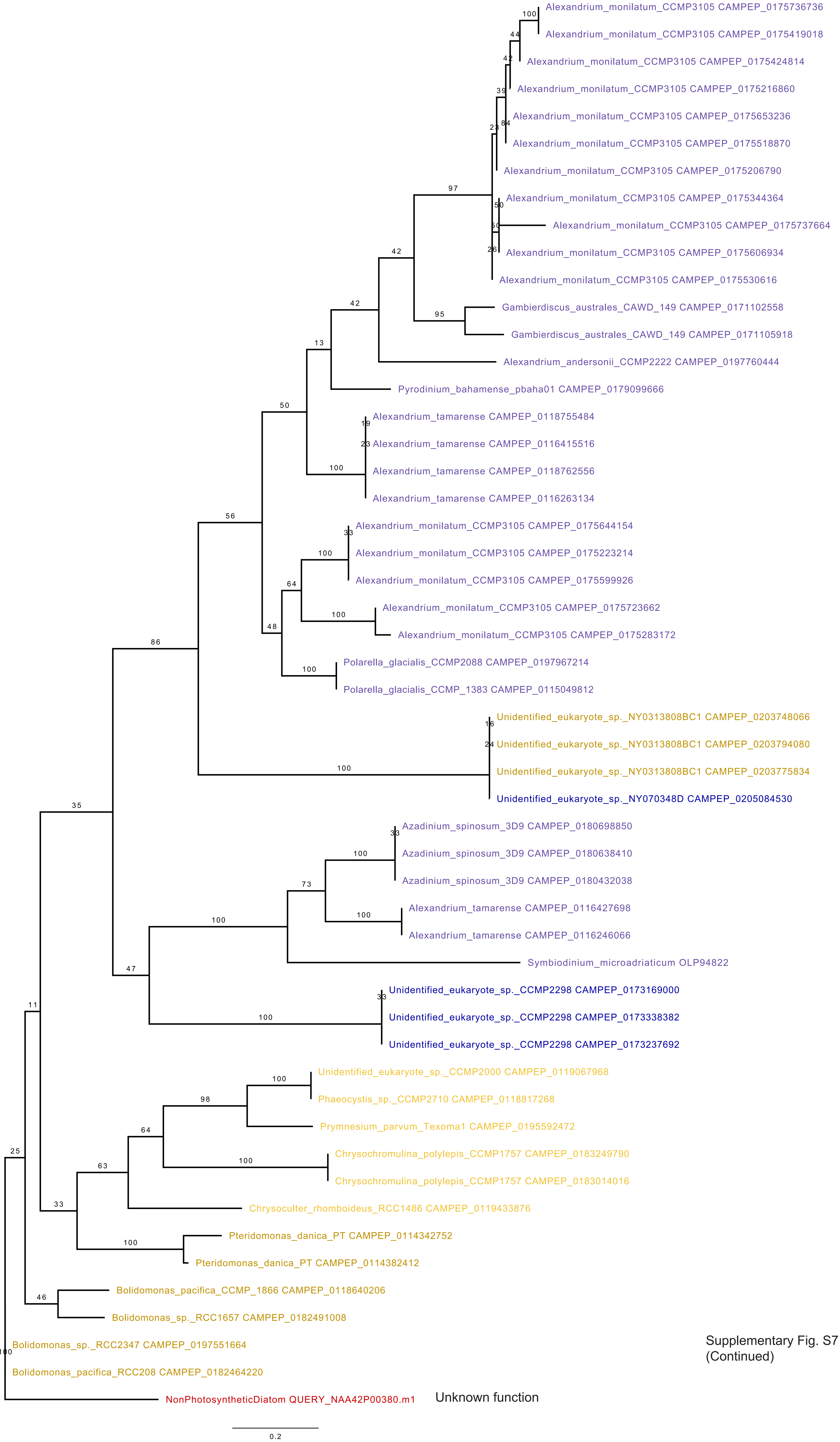

Supplementary Fig. S7
(Continued)

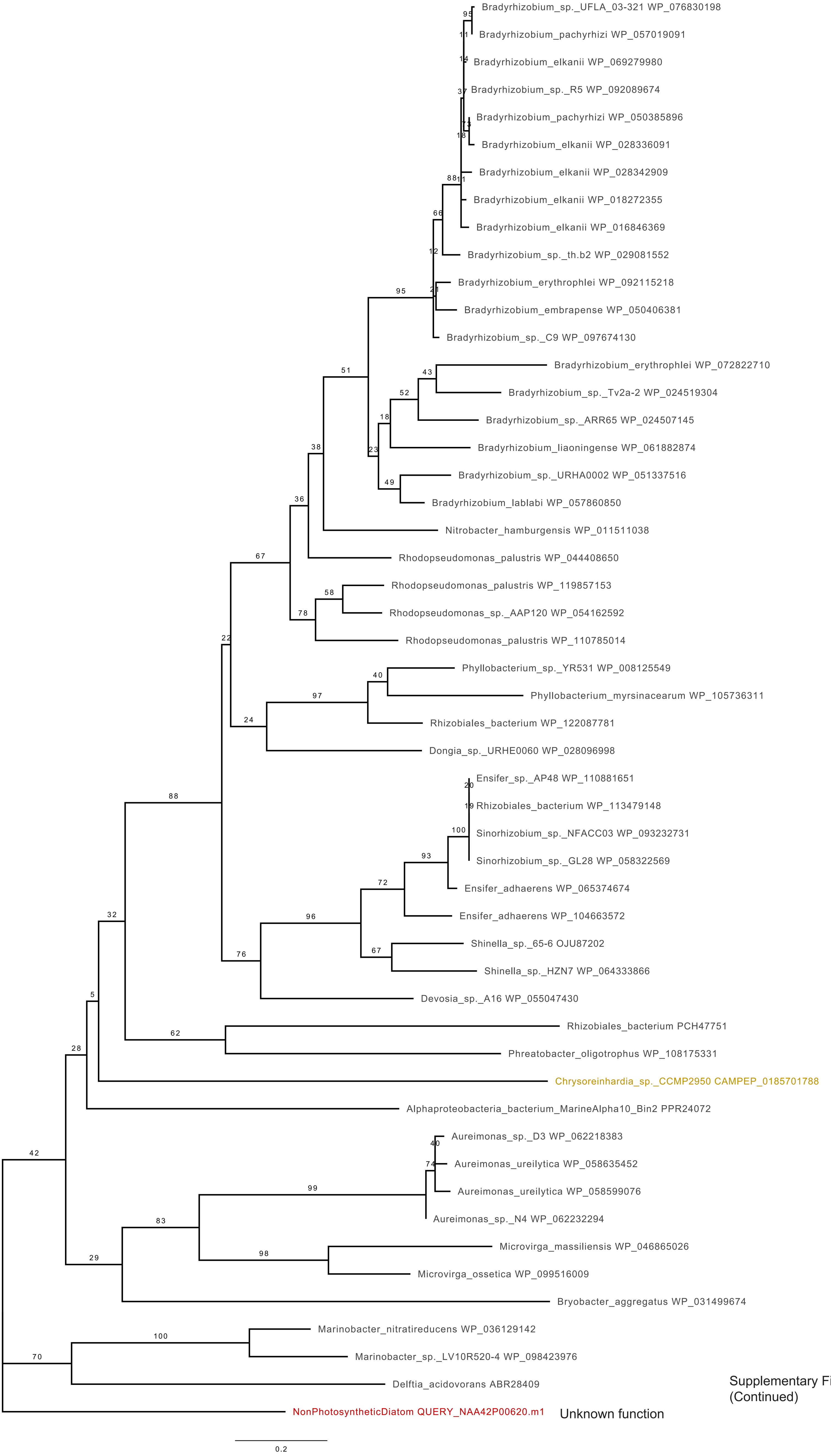

Supplementary Fig. S7
(Continued)

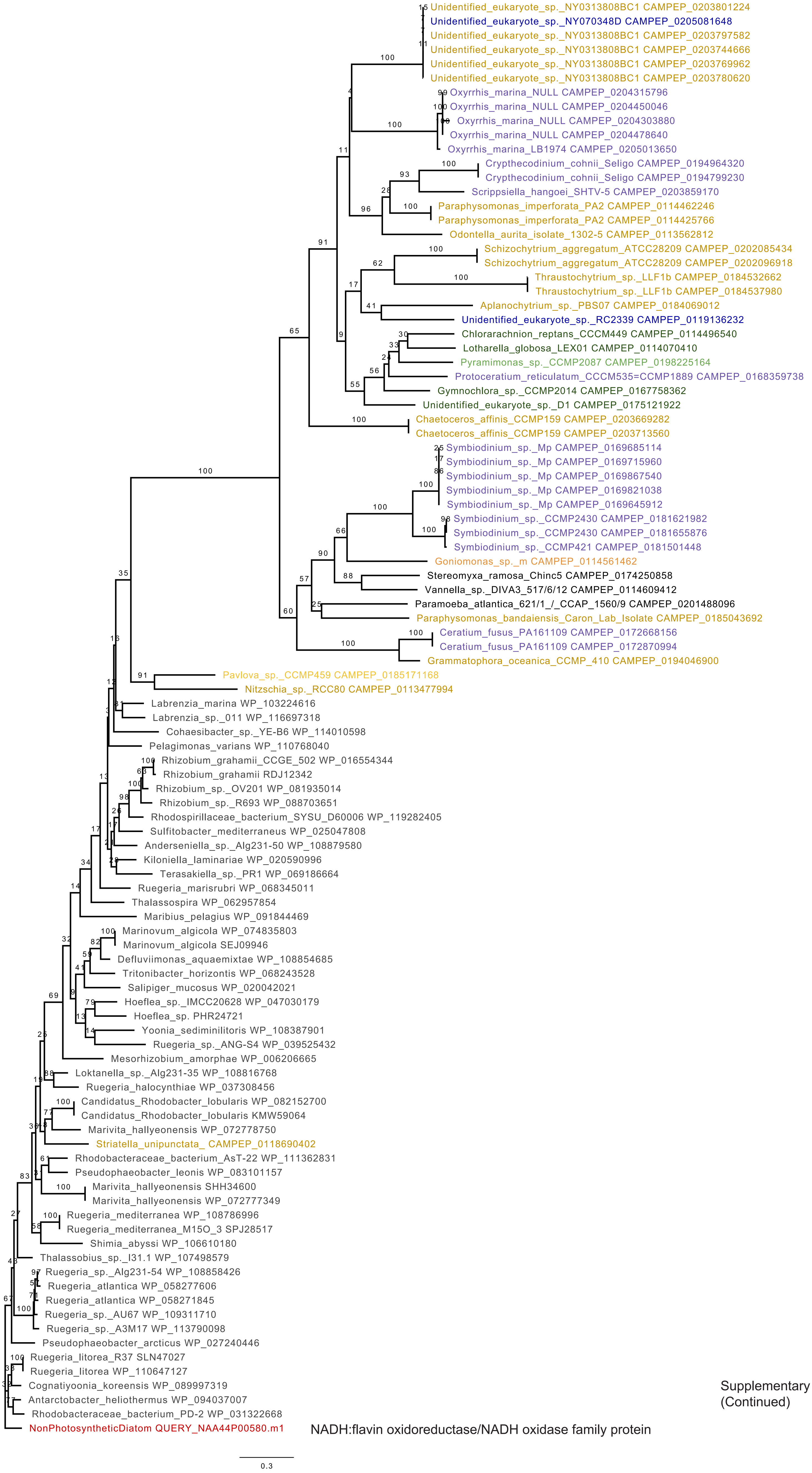

NADH:flavin oxidoreductase/NADH oxidase family protein

Supplementary Fig. S7
(Continued)

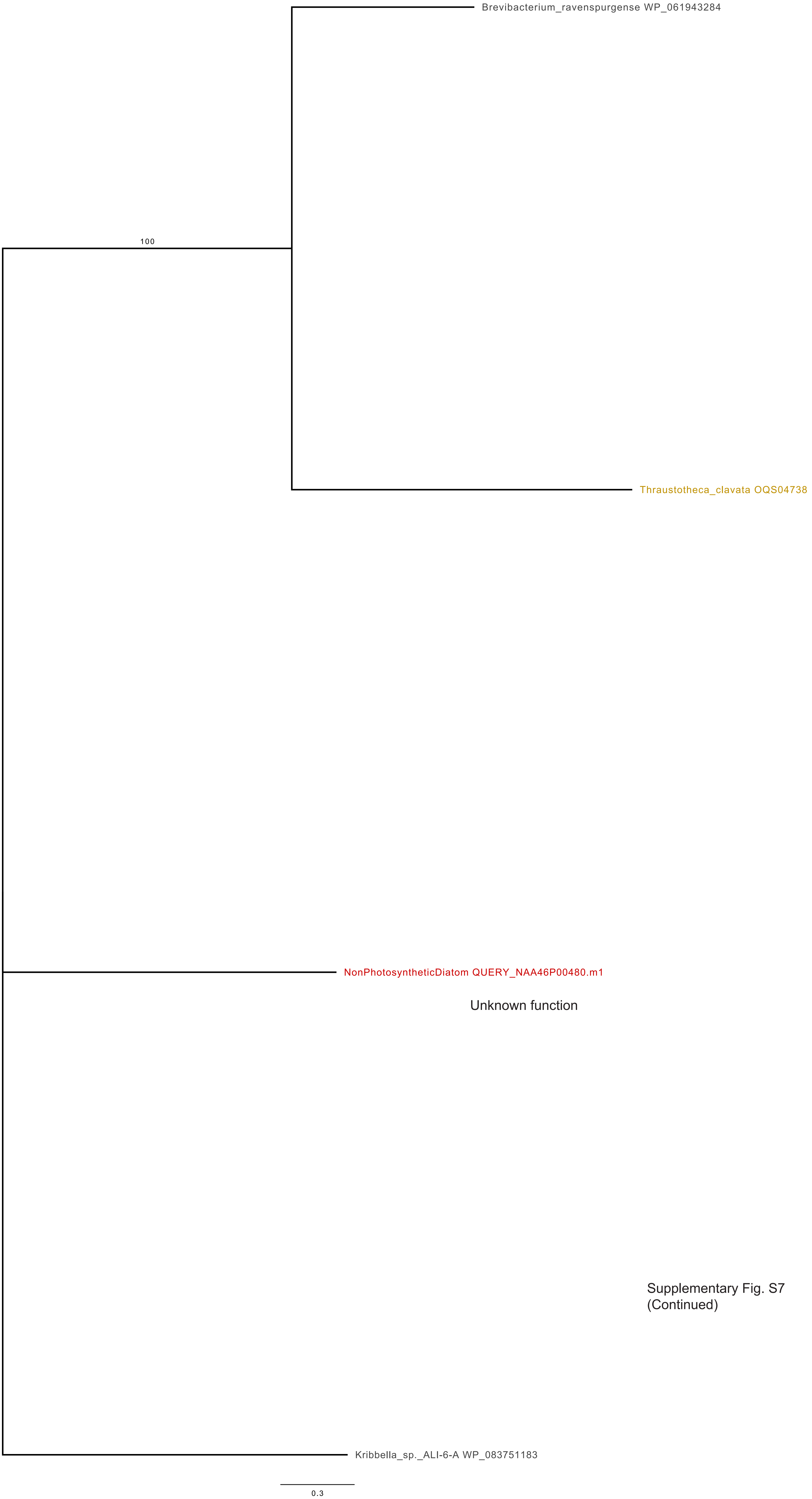

Supplementary Fig. S7
(Continued)

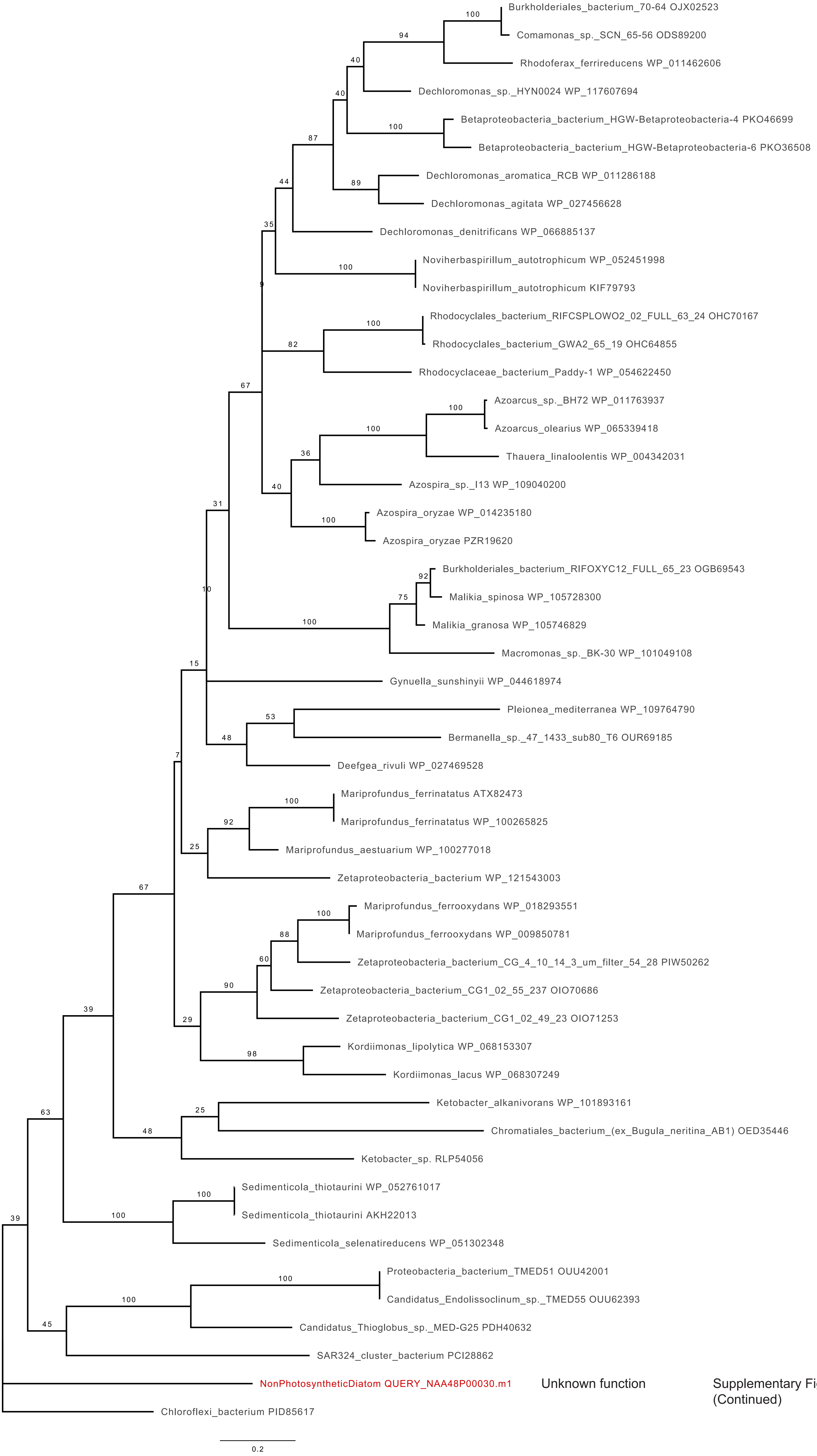

Supplementary Fig. S7
(Continued)

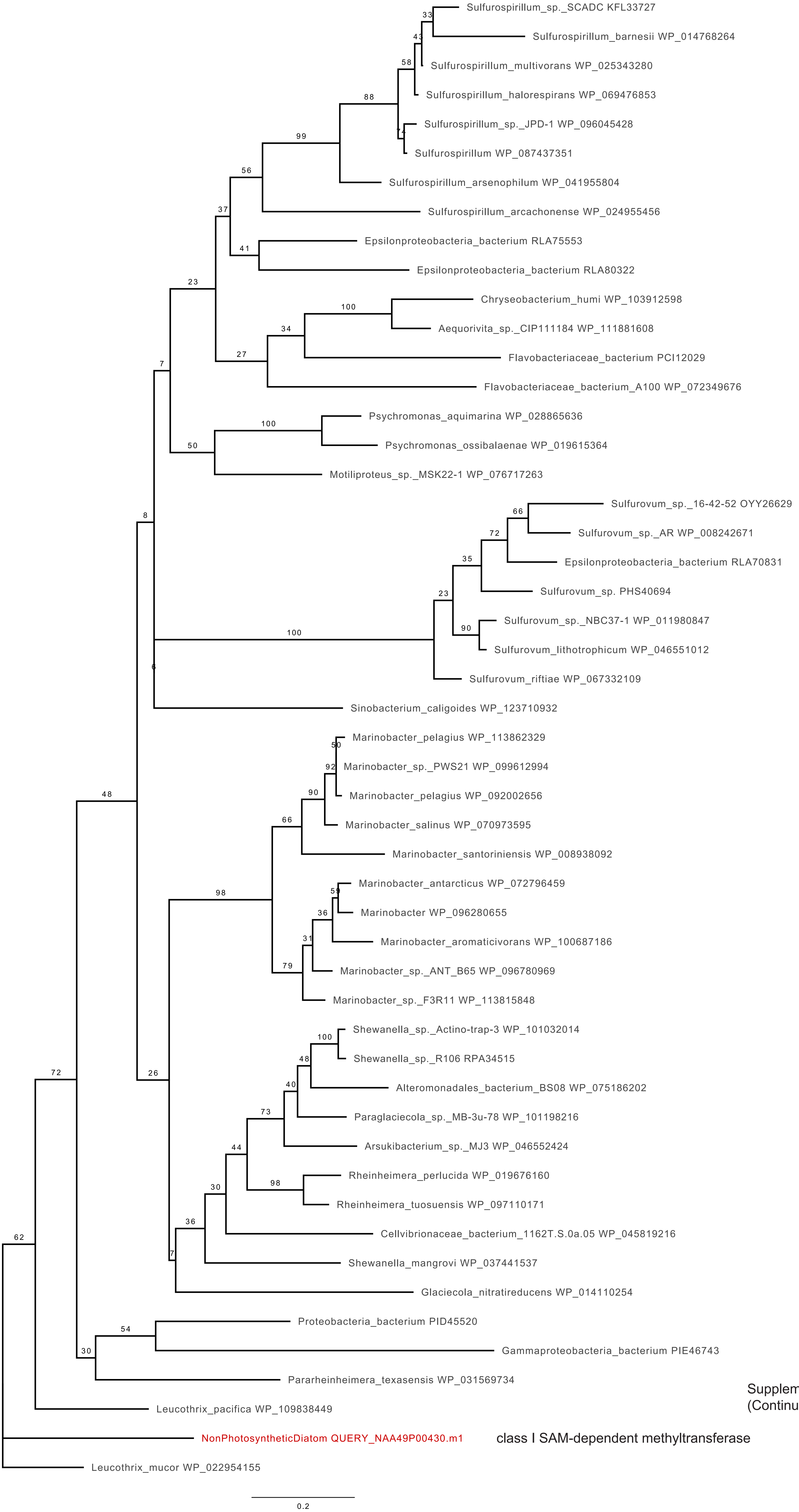

Supplementary Fig. S7
(Continued)

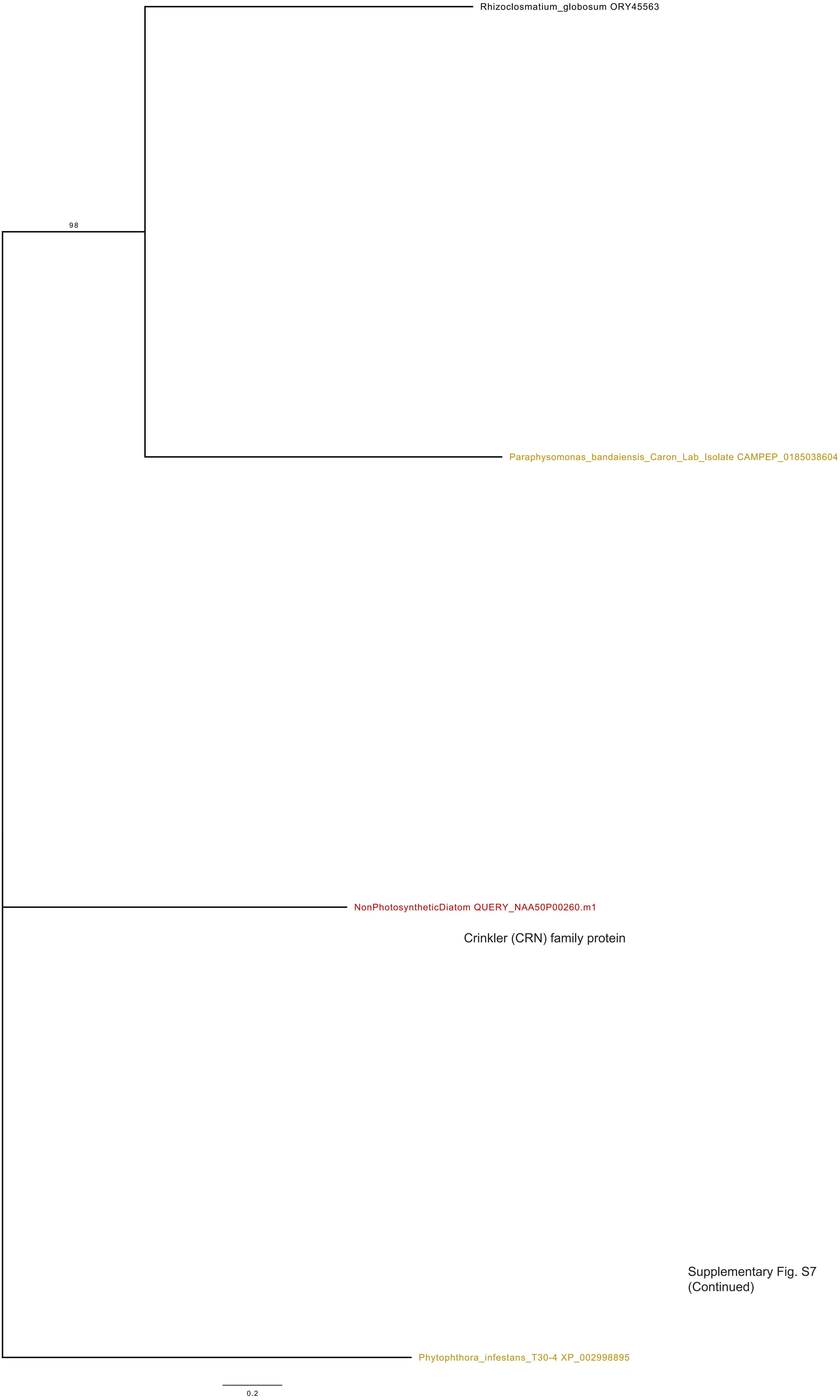

Crinkler (CRN) family protein

Supplementary Fig. S7
(Continued)

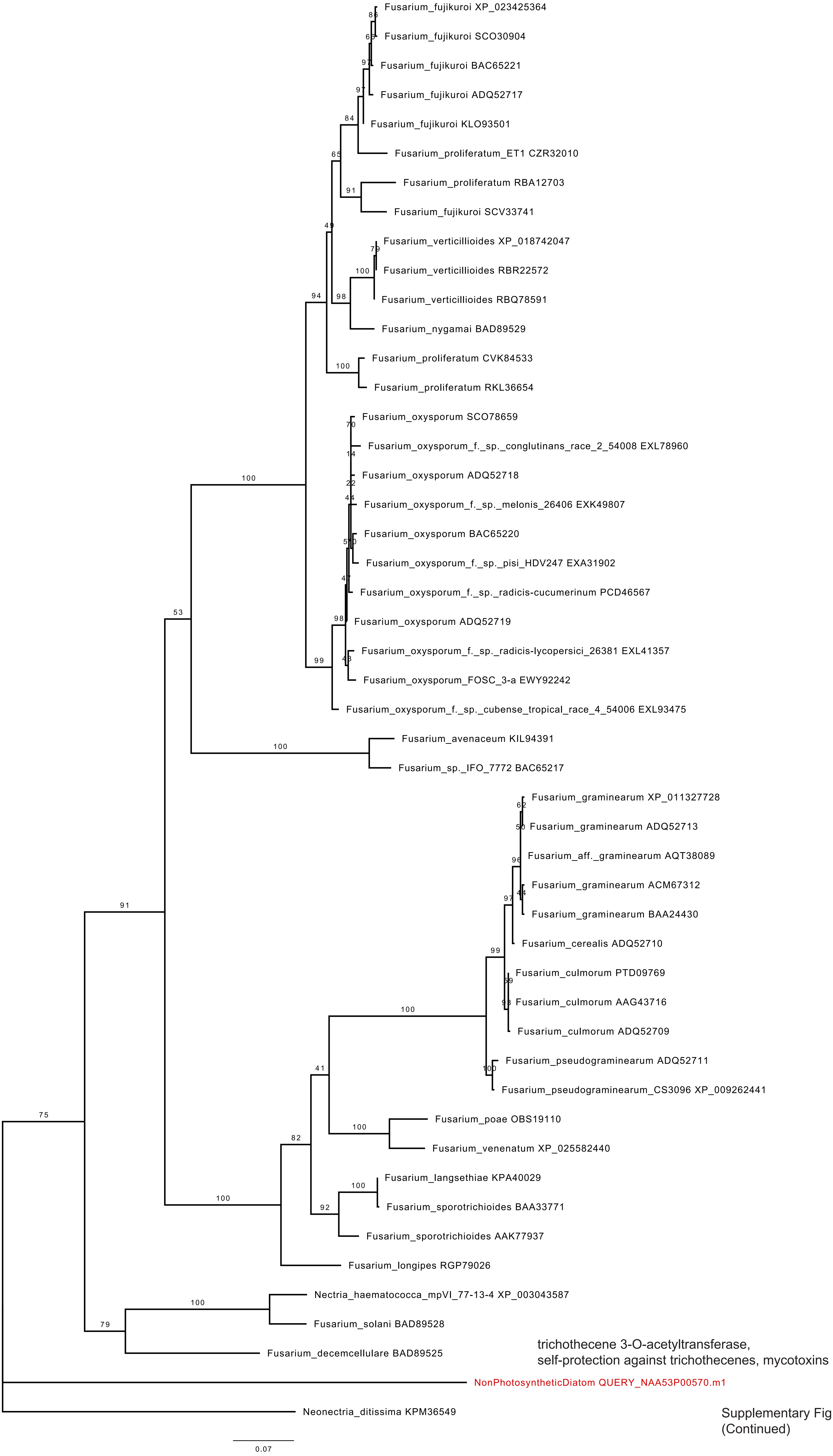

Supplementary Fig. S7
(Continued)

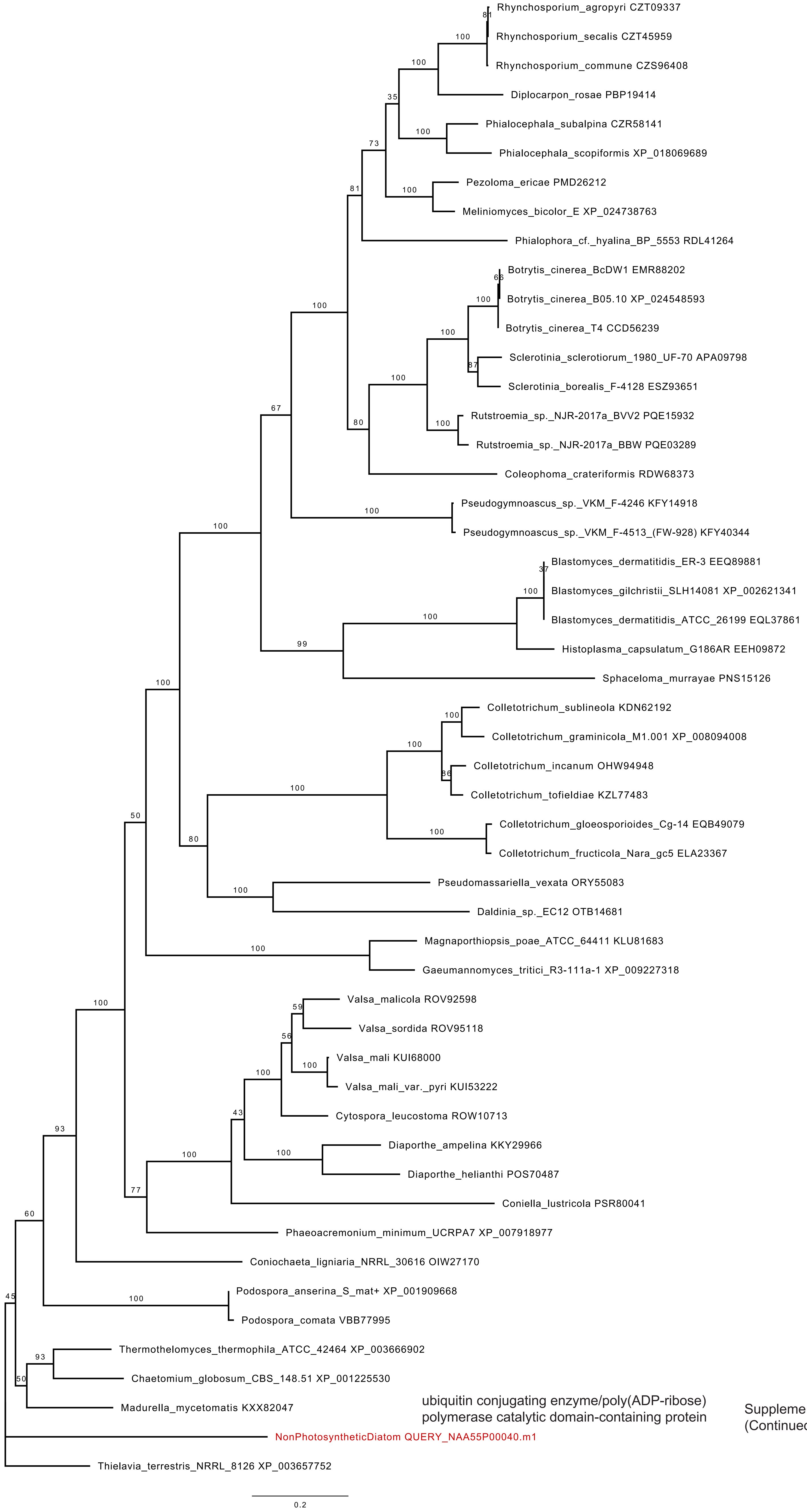

ubiquitin conjugating enzyme/poly(ADP-ribose) polymerase catalytic domain-containing protein

Supplementary Fig. S7 (Continued)

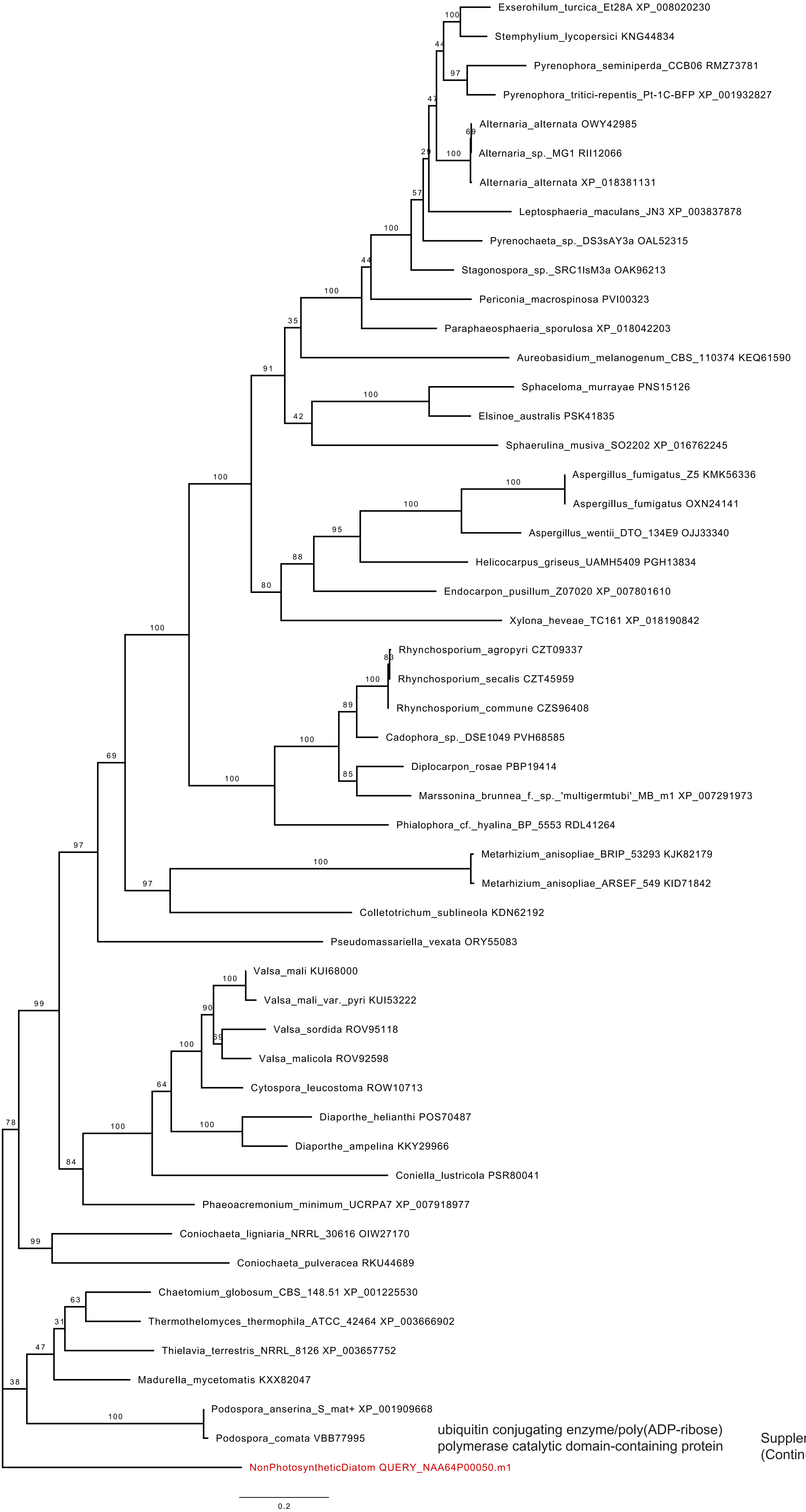

ubiquitin conjugating enzyme/poly(ADP-ribose) polymerase catalytic domain-containing protein

Supplementary Fig. S7 (Continued)

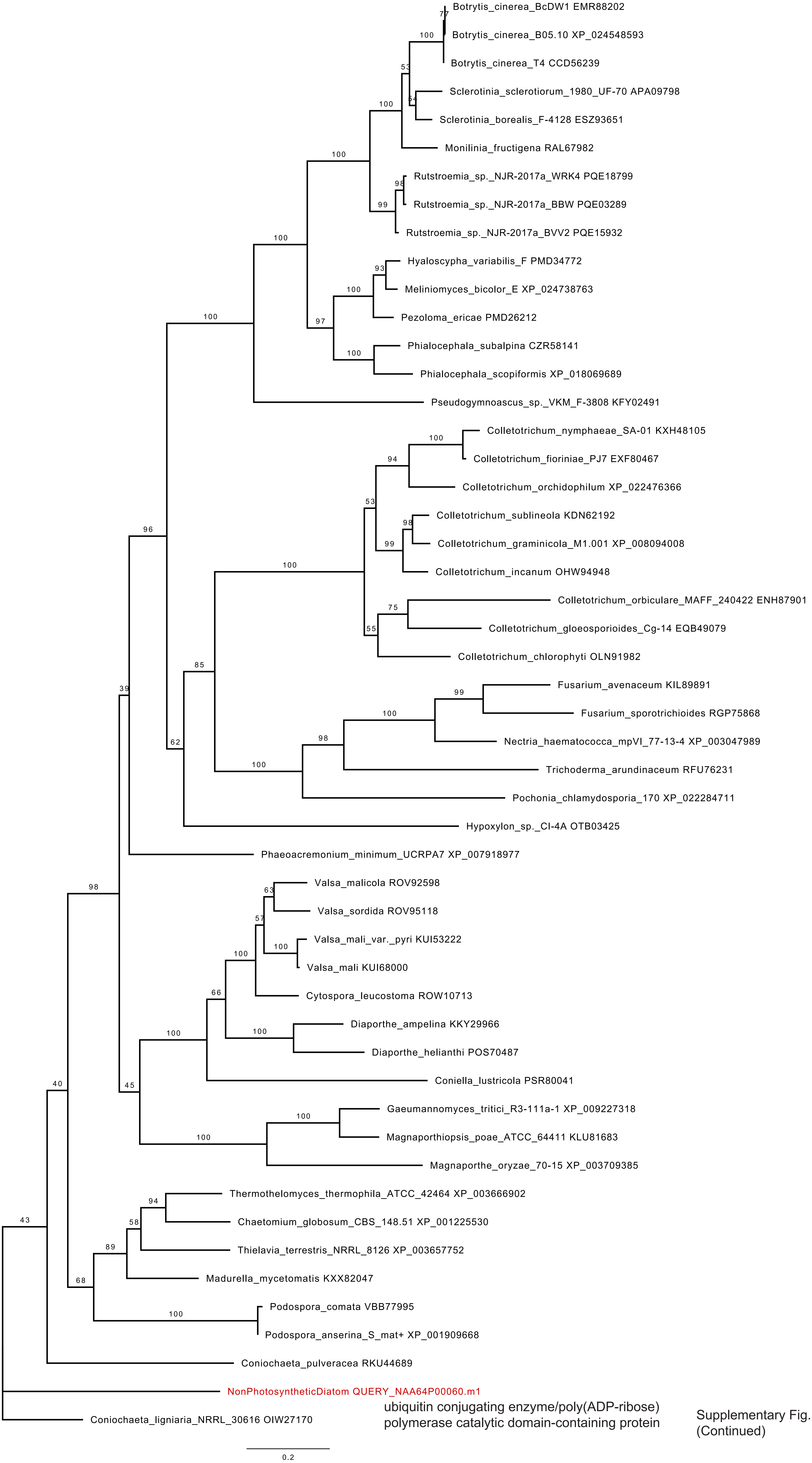

ubiquitin conjugating enzyme/poly(ADP-ribose) polymerase catalytic domain-containing protein

Supplementary Fig. S7 (Continued)

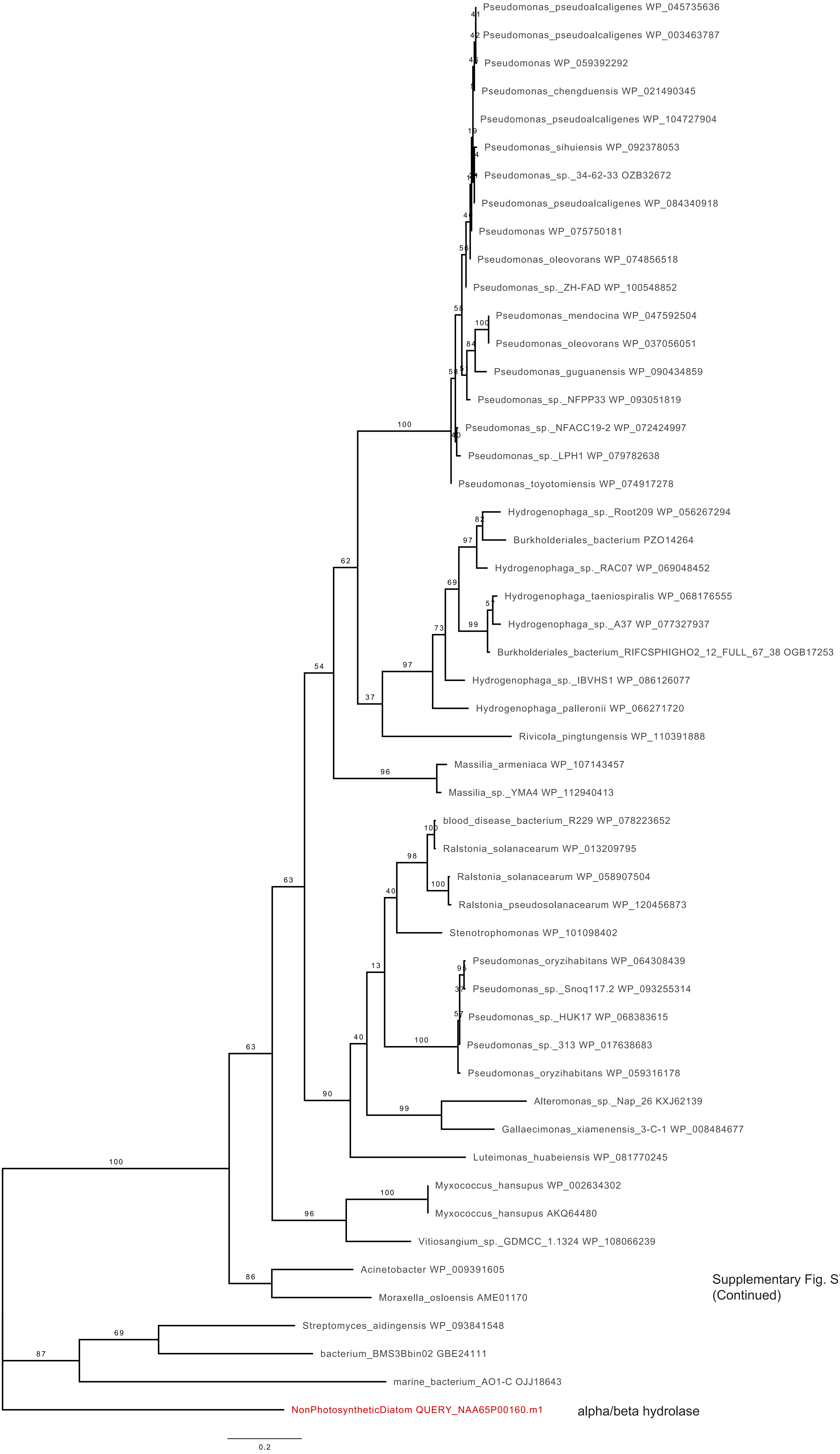

Supplementary Fig. S7
(Continued)

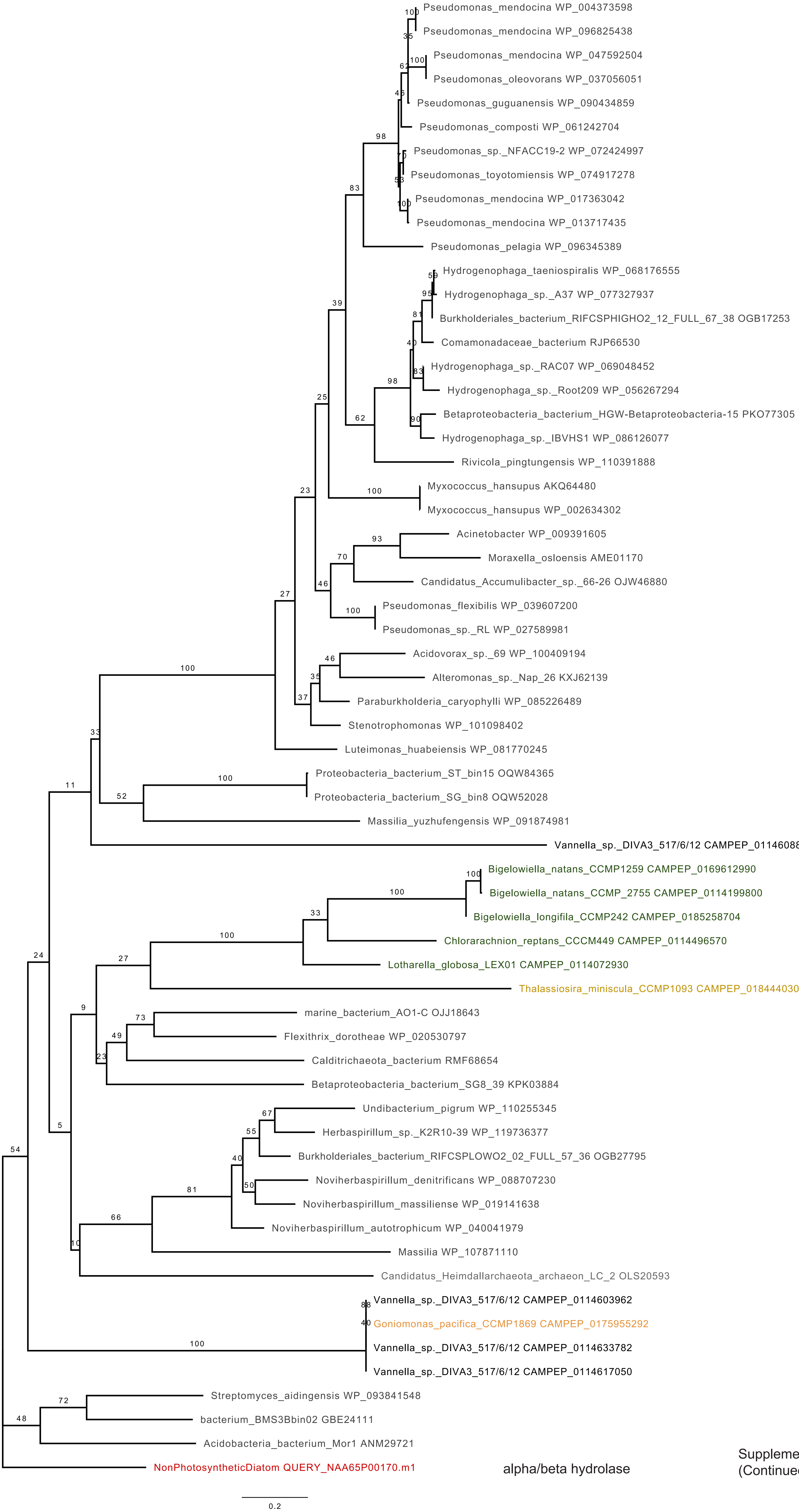

Supplementary Fig. S7 (Continued)

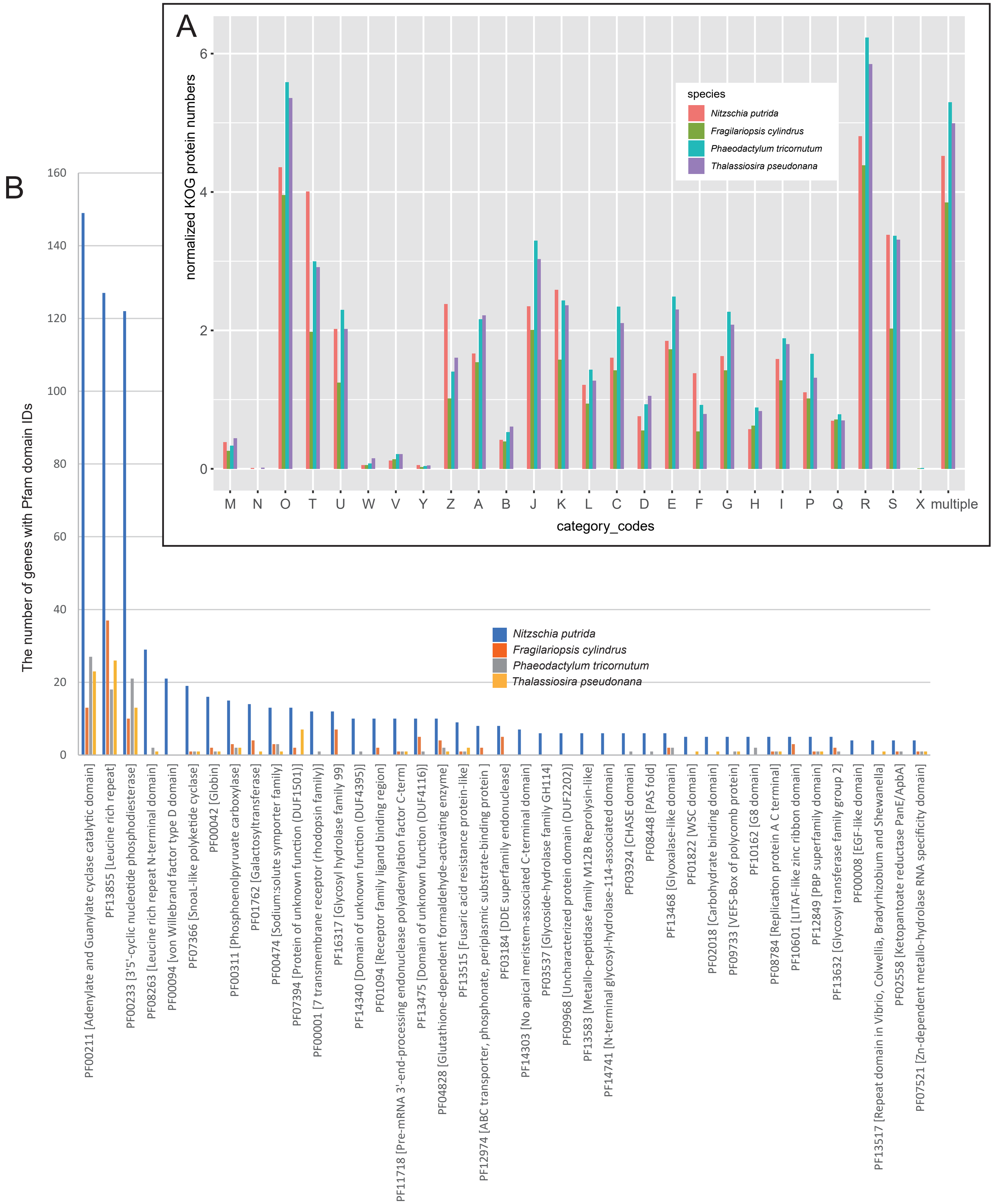

**Supplementary Fig. S8 The number of genes to which 41 Pfam domain IDs are assigned in *Nitzschia putrida*.** A. Comparison of the number of genes assigned to each KOG category, normalized by the total gene number of each genome. Other details are described Fig. 3F. B. Here chosen are Pfam IDs containing at least 4 *Nitzschia* sequences and of which assigned sequences in *Nitzschia* are as 4 or more times large as the mean number of sequences with the same Pfam IDs in the 3 photosynthetic diatoms.

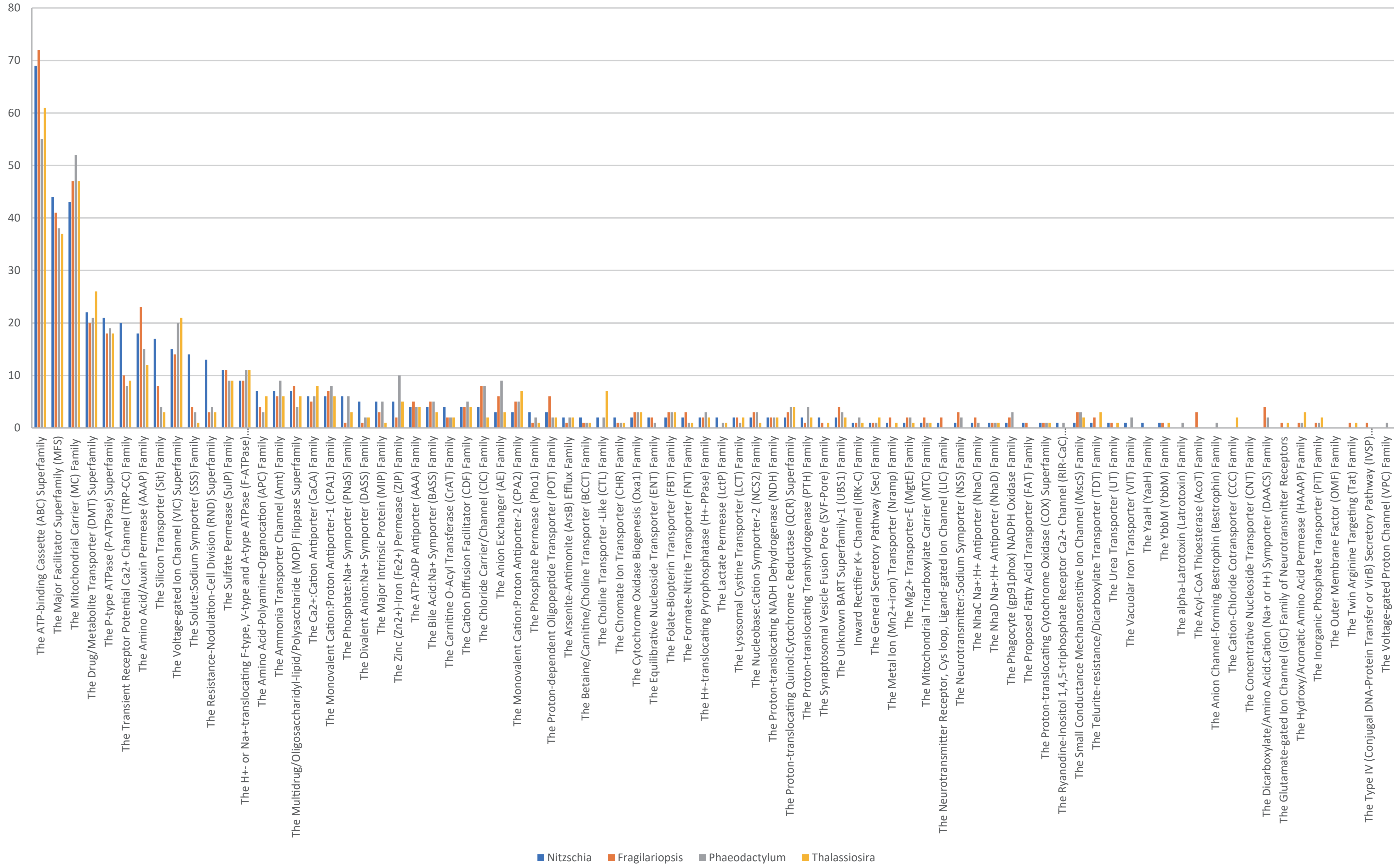

**Supplementary Fig. S9 Transporter genes in the four diatoms.** Transporters were identified by TransportTP (http://bioinfo3.noble.org/transporter/) (Li et al. 2009).

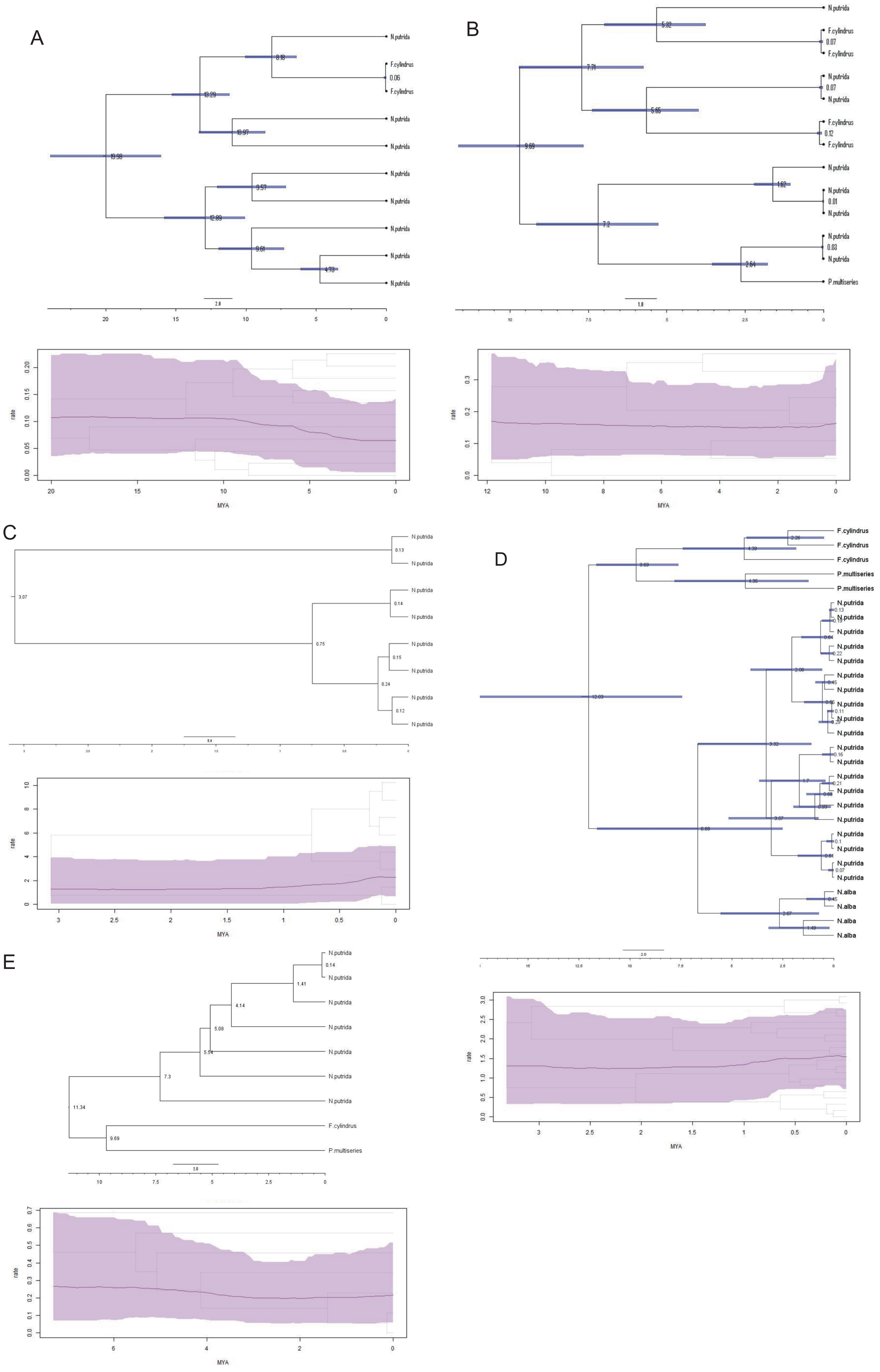

A
N.putrida
F.cylindrus
rate
MYA
B
N.putrida
F.cylindrus
P.multiseries
rate
MYA
C
N.putrida
rate
MYA
D
F.cylindrus
P.multiseries
N.putrida
N.alba
rate
MYA
E
N.putrida
F.cylindrus
P.multiseries
rate
MYA

*Supplementary Fig. S10 Expansion rates and divergence estimates of transporter gene families.* Phylogenetic Maximum Clade Credibility (MCC) tree summarised by TreeAnnotator v2.6.1. Expansion rate was calculated using the Bayesian inference lineage diversification rate analysis tool TESS for R (Höhna, May and Moore, 2016) using the MCC phylogenetic tree produced by BEAST v2.6.1 (Bouckaert et al., 2019). Divergence estimates were obtained using Bayesian Markov Chain Monte Carlo (MCMC) analyse using corresponding sequences from related species implemented in Beast v2.6.1. (Bouckaert et al., 2019). Divergence estimates for all nodes are given in Millions of years (Myr) before present. A. Myosin (control) for *Fragilariopsis cylindrus* (n=2) and the largest cluster of expanded Myosin gene family of *Nitzschia putrida* NIES-4235 (n=8). There is no evidence for a significant change in the expansion rate in the past 20 million years. B. Ammonium transporters (control) for *Pseudo-nitzschia multiseries* (n=1), *Fragilariopsis cylindrus* (n=4) and the largest cluster of expanded NH+ gene family of *Nitzschia putrida* NIES-4235 (n=8). There is no evidence for a significant change in the expansion rate in the past 12 million years. C. Resistance-nodulation-cell division superfamily for the largest cluster of expanded RND gene family of *Nitzschia putrida* NIES-4235 (n=8). There is no evidence for a significant change in the expansion rate in the past 3 million years. D. Silicon transporters for *Fragilariopsis cylindrus* (n=3), *Pseudo-nitzschia multiseries* (n=2), *Nitzschia alba* (n=4) and the expanded SIT gene family of *Nitzschia putrida* NIES-4235 (n=20). There is no evidence for a significant change in the expansion rate in the past 3.3 million years. E. Solute:sodium symporters for *Pseudo-nitzschia multiseries* (n=1), *Fragilariopsis cylindrus* (n=1) and the largest cluster of expanded SST gene family of *Nitzschia putrida* NIES-4235 (n=7). There is no evidence for a significant change in the expansion rate in the past 7.3 million years.

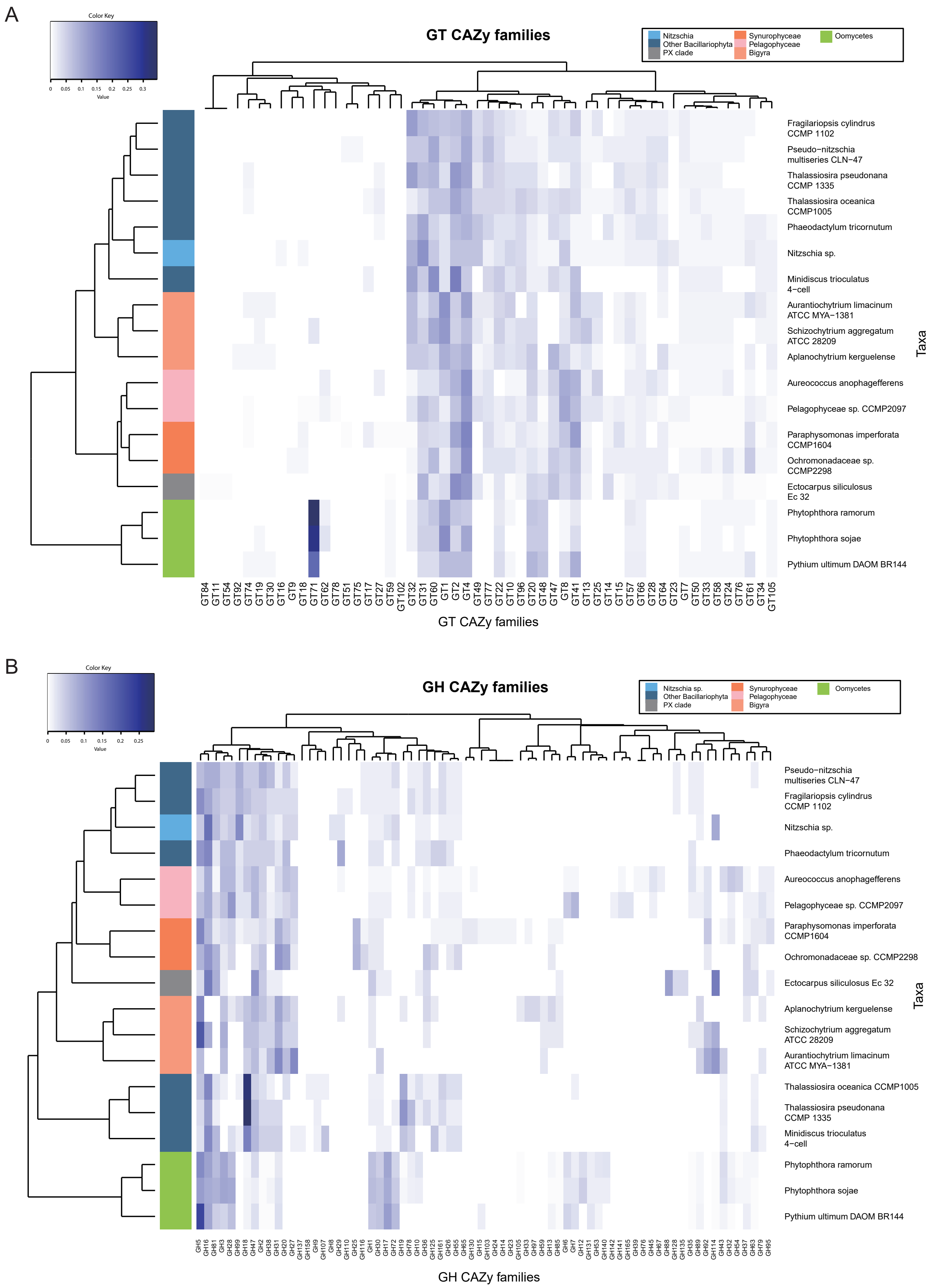
A
Color Key
0 0.05 0.1 0.15 0.2 0.25 0.3
Value
GT CAZy families
Nitzschia
Other Bacillariophyta
PX clade
Synurophyceae
Pelagophyceae
Bigyra
Oomycetes
Fragilariopsis cylindrus CCMP 1102
Pseudo−nitzschia multiseries CLN−47
Thalassiosira pseudonana CCMP 1335
Thalassiosira oceanica CCMP1005
Phaeodactylum tricornutum
Nitzschia sp.
Minidiscus trioculatus 4−cell
Aurantiochytrium limacinum ATCC MYA−1381
Schizochytrium aggregatum ATCC 28209
Aplanochytrium kerguelense
Aureococcus anophagefferens
Pelagophyceae sp. CCMP2097
Paraphysomonas imperforata CCMP1604
Ochromonadaceae sp. CCMP2298
Ectocarpus siliculosus Ec 32
Phytophthora ramorum
Phytophthora sojae
Pythium ultimum DAOM BR144
Taxa
GT84 GT11 GT54 GT92 GT74 GT19 GT30 GT16 GT9 GT18 GT71 GT62 GT78 GT51 GT75 GT17 GT27 GT59 GT102 GT32 GT31 GT60 GT1 GT2 GT4 GT49 GT77 GT22 GT10 GT96 GT20 GT48 GT47 GT8 GT41 GT13 GT25 GT14 GT15 GT57 GT66 GT28 GT64 GT23 GT7 GT50 GT33 GT58 GT24 GT76 GT61 GT34 GT105
GT CAZy families
B
Color Key
0 0.05 0.1 0.15 0.2 0.25
Value
GH CAZy families
Nitzschia sp.
Other Bacillariophyta
PX clade
Synurophyceae
Pelagophyceae
Bigyra
Oomycetes
Pseudo−nitzschia multiseries CLN−47
Fragilariopsis cylindrus CCMP 1102
Nitzschia sp.
Phaeodactylum tricornutum
Aureococcus anophagefferens
Pelagophyceae sp. CCMP2097
Paraphysomonas imperforata CCMP1604
Ochromonadaceae sp. CCMP2298
Ectocarpus siliculosus Ec 32
Aplanochytrium kerguelense
Schizochytrium aggregatum ATCC 28209
Aurantiochytrium limacinum ATCC MYA−1381
Thalassiosira oceanica CCMP1005
Thalassiosira pseudonana CCMP 1335
Minidiscus trioculatus 4−cell
Phytophthora ramorum
Phytophthora sojae
Pythium ultimum DAOM BR144
Taxa
GH5 GH16 GH81 GH3 GH28 GH99 GH18 GH47 GH2 GH38 GH31 GH20 GH27 GH137 GH158 GH9 GH107 GH8 GH29 GH110 GH25 GH116 GH1 GH30 GH17 GH72 GH19 GH78 GH10 GH36 GH125 GH161 GH26 GH55 GH65 GH130 GH15 GH103 GH24 GH14 GH23 GH105 GH33 GH97 GH59 GH13 GH85 GH6 GH7 GH12 GH131 GH53 GH140 GH142 GH141 GH165 GH39 GH76 GH45 GH67 GH88 GH128 GH135 GH35 GH89 GH92 GH114 GH43 GH32 GH54 GH37 GH63 GH79 GH95
GH CAZy families

**Supplementary Fig. 11 Carbohydrate-active enzyme (CAZyme) in *N. putrida*.** A. Glycosyltransferases (GT) families from the CAZy database classification (www.cazy.org) in various Stramenopiles. The diagram shows a heatmap of CAZyme prevalence in each taxon (number of a particular CAZy family divided by the total number of CAZymes in the organism); the white to blue color scheme indicates low to high prevalence, respectively. Dendrograms (left and top of the figure) show respectively the relative proximity of taxa with respect co-occurrence of CAZyme families and the co-occurrence of CAZy families with one another within genomes (for details see methods described in Cenci et al. 2018). Nitzschia is in light blue while other Bacillariophyta are in dark blue. We observe that all Bacillariophyta group together indicating a close GT repertoire. B. Glycoside Hydrolases (GH) families in various Stramenopiles. The diagram shows a heatmap of CAZyme prevalence in each taxon (number of a particular CAZy family divided by the total number of CAZyme in the organism); the white to blue colour scheme indicates low to high prevalence, respectively. Dendrograms (left and top) show respectively the relative taxa proximity with respect co-occurrence of CAZyme families and the co-occurrence of CAZyme families with one another within genomes (for details see methods described in Cenci et al. 2018). The same color code as in Figure 3 is used. Bacillariophyta are split into two groups indicating differences in the GH repertoire of *Nitzschia*, *F. cylindrus*, *Pseudo-nitzschia* and *P. tricornutum*, on one hand, and on the other hand *Thalassiosira* spp. and *M. trioculatus.*

A

| | Nitzschia | Fragilariopsis | Phaeodactylum | Thalassiosira |
|---|---|---|---|---|
| total (%) | 978 (6.5) | 998 (5.5) | 596 (5.7) | 718 (6.1) |
| Peptide degradation | 87 | 53 | 44 | 46 |
| Carbohydrate metabolisms | 101 | 89 | 74 | 52 |
| Extracellular matrix/adhesion/cell surface associated proteins | 127 | 42 | 19 | 23 |
| Substrate modification | 96 | 58 | 52 | 49 |
| Redox | 16 | 14 | 7 | 13 |
| Others including unknown functions | 551 | 742 | 400 | 535 |

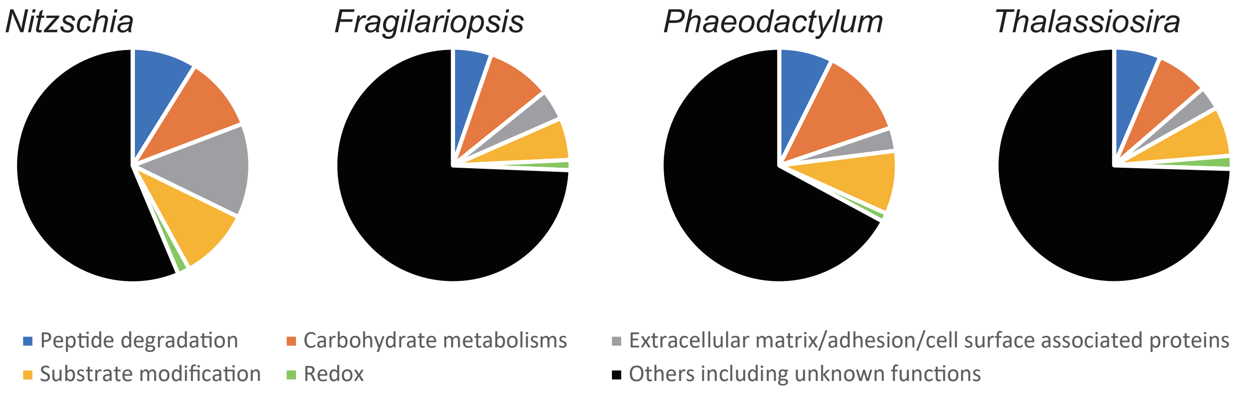

B

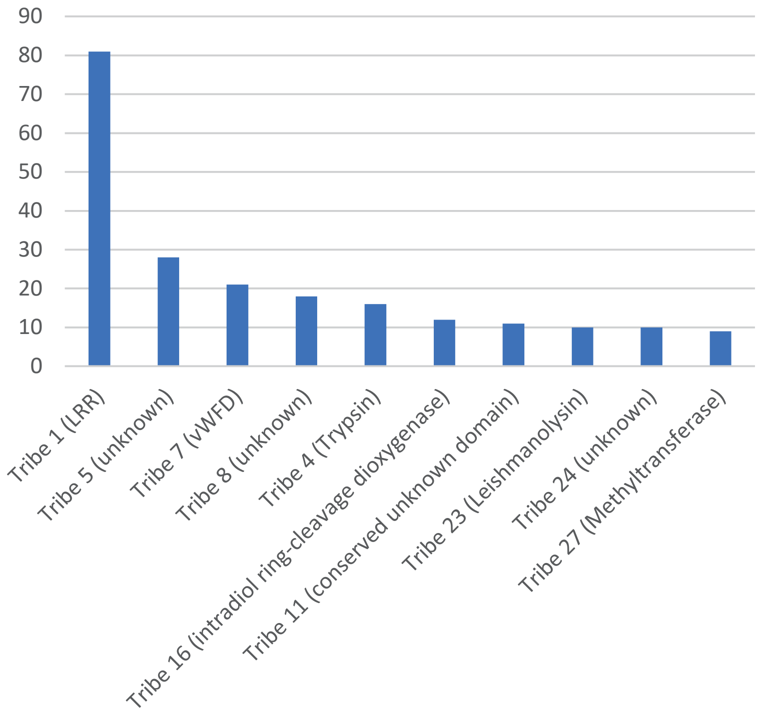

**Supplementary Fig. 12 Secretomes in diatoms.** A. The number of genes for predicted secreted proteins in diatoms and their annotated functions. B. Ten most abundant secretome sequences in *N. putrida*. Assigned protein domains or functions are described in parentheses. Tribes with no parenthesis are those of unknown functions.

Table S1 Genomes of non-photosynthetic and photosynthetic diatoms

| **Species** | ***Nitzschia putrida*** | ***Fragilariopsis cylindrus*** | ***Phaeodactylum tricornutum*** | ***Thalassiosira pseudonana*** |
|---|---|---|---|---|
| **Genome size (Mb)** | 35 (47) | 61 | 27 | 32 |
| **N50 (Mb)** | 0.86 (0.55) | 1.3 | 0.945 | 1.9 |
| **G+C (%)** | 47.6 (47.6) | 39.8 | 48.8 | 47 |
| **Gene count** | 15,003 (20,461) | 21,066[*1] | 10,402 | 11,776 |
| **Av gene length (bp)** | 1611 (1619) | 1575 | 1511 | 1553 |
| **Av exon length (bp)** | 901 (907) | 625 | 842 | 612 |
| **Av intron length (bp)** | 122 (121) | 245 | 135 | 124 |
| **Av exon number/gene** | 1.69 (1.68) | 2.08 | 1.79 | 2.49 |

[*1] including diverged alleles. Numbers in parentheses are of total numbers in primary contigs and haplotigs.

**Table S2: Tests for positive selection among Silicon Transporter (SIT) codons using Codeml site models.**

| Model | dN/dS[a] | lnL | 2ΔL[b] | Estimates of parameters | Positively selected sites (BEB)[c] |
|---|---|---|---|---|---|
| M0 (one ratio) | 0.1018 | -3893.91 | | ω=0.1018 | NA |
| M3 (discrete) | 0.1422 | -3804.88 | 178.05, df=4, p-value= < 2.2e-16 | p0=0.90449, p1=0.09137, p2=0.00413, ω0=0.02266, ω1=0.93222, ω2=8.85203 | 0 |
| M1a (neutral) | 0.1111 | -3813.95 | | p0=0.90976, p1=0.09024, ω0=0.02292, ω1=1.00000 | NA |
| M2a (selection) | 0.1452 | -3804.93 | 18.04, df=2, p-value=0.000121 | p0=0.90887, p1=0.08718, p2=0.00395, ω0=0.02394, ω1=1.00000, ω2=9.17486 | 8(2) |
| M7 (beta) | 0.1192 | -3815.95 | | p=0.03898, q=0.28910 | NA |
| M8 (beta & ω) | 0.1401 | -3805.67 | 20.57, df=2, p-value=3.42e-05 | p0= 0.99541, p=0.06255, q=0.53977, p1=0.00459, ω=8.13565 | 12(3) |

[a] Average dN/dS over all sites

[b] Likelihood ratio test statistic. Calculate with lr.test from extRemes package in R

[c] Inferred with Bayes Empirical Bayes at the 50% (95%) posterior probability cutoff
